# Cocaine use disorder is associated with lower brain state transition energy particularly in higher order and excitatory networks

**DOI:** 10.1101/2025.08.06.668699

**Authors:** Hussain Bukhari, Aliza Brzezinski-Rittner, S. Parker Singleton, Ceren Tozlu, Keith Jamison, Luis Concha, Eduardo Garza-Villarreal, Conor Liston, Amy Kuceyeski

**Author notes:** Senior Author (Conor Liston, Amy Kuceyeski). **Data Availability Statement** We used the Sudmex Conn data set, consisting of multimodal magnetic resonance imaging, CUD metrics, and demographics of 132 individuals (18-50 years), with 61 (11 female) non-user controls and 71 individuals with chronic CUD (9 females). The data is available publicly in raw form at https://openneuro.org/datasets/ds003346. This project used open-source code published by Cornblath et. al to deploy network control theory at https://github.com/ejcorn/brain_states.

## Abstract

Cocaine use disorder (CUD) detrimentally impacts personal health, social relationships, and economic opportunity. Here, we assess CUD-associated shifts in brain dynamics using Network Control Theory and examine how they align with previously identified changes in neurological systems and behavioral profiles of people with CUD. The SUDMEX CONN dataset consists of multi-modal MRI, cocaine use metrics, behavioral measures, and demographics of individuals with CUD (N=132, 71 CUD). We identified recurring brain activity states and used NCT to calculate the transition energy (TE) between pairs of states. ANCOVAs examined global and regional TE associations with drug use group (CUD vs controls (NC)), years of CUD, and risk-taking behaviors. We identified potential mechanisms driving the differences by correlating CUD-related regional TE effects with neurotransmitter/receptor systems. People with CUD had significantly lower global TE and default mode, dorsal attention and limbic network TE compared to non-user controls, particularly in regions enriched for noradrenaline and mu opioid receptors. Longer duration of CUD was associated with more decreased global TE, top-down TE, default mode, control and ventral attention network TE, and regional TE enriched for excitatory neurotransmitters and receptors. People with CUD needed to expend more global and top-down TE to perform better on a risk-taking task (the Iowa Gambling task), an effect which was not found in NCs. Our analysis of whole-brain activity dynamics provides a link between the effects of upstream glutamatergic excitotoxicity and/or opioid receptor dysfunction, and downstream weakening of inhibitory control that is central to CUD.

## 1. Introduction

Given that around 15% of casual users progress to long-term consumption, cocaine use disorder (CUD) poses significant public health challenges^1,2^. Reports of increased CUD make it critical to identify how brain dynamics shift as recreational use becomes pathological. Cocaine exposure alters brain excitatory and inhibitory balance by dysregulating glutamatergic and dopaminergic signaling. Changes in regions essential for response inhibition like the anterior and middle cingulate cortices^3–6^ are accompanied by differential expression of dopamine and glutamate receptors, impairing top-down control and interfering with GABAergic transmission and reducing inhibitory tone. This imbalance triggers neurotoxicity via excitotoxicity, causes cellular death, and reductions in gray and white matter volume^7^ that may lead to the loss of inhibitory control seen in CUD and explaining compulsive drug-seeking.

Multimodal imaging has revealed widespread alterations in the brains of people with CUD in circuits linked to cognitive control and reward processing^8^. Aberrant signaling in dopamine- and glutamate-rich circuits, particularly in response to drug cues^9–12^, accompanies altered prefrontal inhibition during cognitive tasks, notably in regions of the default mode network (DMN)^10^. At rest, functional connectivity weakens between the DMN and other systems, reflecting impaired integration across cortical and subcortical domains^13–15^, affecting sensitivity to salient stimuli, and increasing relapse risk. However, these changes cannot be attributed to FC but may reflect downstream effects of upstream dopamine receptor changes.

Network control theory (NCT) offers a dynamic systems-level framework to examine how brain state transitions are altered in neurological and psychiatric conditions as well as substance use disorders. By modeling how activation (measured via fMRI) flows across structural white matter pathways (measured via diffusion MRI), NCT estimates the control energy required to transition (TE) between patterns of brain activity, or states^16^. TE estimates reveal pathophysiological signatures of disease/disorder and recovery, which in turn allow identification of potential treatment targets. When paired with receptor and transporter maps, NCT can model specific neurotransmitter systems and their effect on neural dynamics in addiction^17^. Previously, we used NCT to understand shifts in brain dynamics in psychedelics like LSD, psilocybin, and DMT, GABAergic substances like alcohol, as well as the heritable effects of having a family history of substance use disorder^18–22^.

For this study, we apply NCT to understand brain dynamics in CUD. Using the SUDMEX-CONN dataset^23^ (n=132, 71 CUD, we identify CUD-related differences in brain state metrics and transition probabilities. We use NCT to compute TE at global, network, and regional scales, testing for CUD-related group differences and associations with length of CUD, while controlling for covariates. Then, we compare our CUD-related TE shifts to receptor/transporter density maps to identify possible mechanistic underpinnings of CUD-related differences in brain dynamics. Finally, we test if CUD changes the association between TE and performance on the Iowa Gambling Task (IGT), a decision-making task relevant to addiction^24^. We demonstrate the advantage of using NCT modeling of brain dynamics by associating our findings with both potential upstream molecular underpinnings and downstream behavioral changes that occur in CUD.

## 2. Methods

### 2.1 Subjects, MRI acquisition and preprocessing and connectome construction

The original dataset was collected in Mexico and approved by the Ethics Committee of the Instituto Nacional de Psiquiatría. Procedures were conducted in accordance with the Declaration of Helsinki, with 145 participants scanned using a Philips Ingenia 3T MRI system (Philips Healthcare, Best, The Netherlands, and Boston, MA, USA) with a 32-channel dS Head coil. Subjects underwent resting-state, T1-weighted, and High Angular Resolution Diffusion Imaging scans. Collection procedures and the diagnostic criteria for CUD are described previously^23^. Individuals were diagnosed with CUD using the Mini International Neuropsychiatric Interview – Plus, Spanish version 5.0.0, administered by trained psychiatrists. To be included, participants had to be actively consuming cocaine or within an abstinence period of less than two months prior to screening. The final sample consisted of 74 patients with CUD (9 female) and 64 healthy controls (6 female). Of these 138 individuals, 132 were included in the analysis. The excluded participants lacked the necessary behavioral and demographic data.

MRI preprocessing was conducted using fMRIPrep v22.0.1 and in-house code to parcellate the brain using the Schaeffer atlas (100 cortical regions)^25^ and the Tian atlas (16 subcortical regions)^26^. Regional fMRI time series were extracted using the Nilearn library and regressed for framewise displacement, white matter, cerebrospinal fluid, global signal, and anatomical and temporal components identified by CompCor. Time series were scrubbed for TRs where motion exceeded 0.5 mm and high-pass filtered using discrete cosine transform basis regressors from fMRIPrep to remove low-frequency signal drifts^23^. Structural connectivity matrices were constructed from DWI by denoising^27^, correcting for subject motion and geometric distortions, and correcting for bias field inhomogeneities^28^. Anatomically-constrained tractography^29^ computed from multi-shell multi-tissue constrained spherical deconvolution of the diffusion signals^30^. The SIFT algorithm was used to filtered tractograms (from 10 to 0.5 million tracks)^31^, and SIFT2 was used to construct weighted adjacency matrices for each subject^32^. Code available at https://github.com/lconcha/sudmex_conn.

### 2.2 Determining Brain States

Following Singleton et al. (2022), we used fMRI time series across all participants (Fig. 1)^19^. K-means clustering identified recurring patterns of brain activation, or “states”, using Pearson’s correlation as the distance metric. The clustering was repeated 100 times to identify the most stable and well-separated solution. The number of clusters was determined with the elbow criterion, and the elbow was observed between k=4 and k=8. (Supp Fig. S1.) We selected k=6, as it offered a balance between explanatory power and interpretability. We repeated the clustering process 10 times and computed adjusted mutual information (AMI) between all resulting partitions, before choosing the partition with the highest total AMI relative to the others for further analysis. Analyses were replicated with k=4 states, as this is a common number of states in previous work.

States (cluster centroids) were characterized using cosine similarity between the states and a binary representation of the eight Yeo resting-state networks (RSNs), see Fig. 1)^33^. Time series were demeaned, so positive values in the centroid represented high-amplitude activations exceeding the mean, while negative values indicated low-amplitude activity lower than the mean. Pearson correlation coefficients were computed between centroids, and anti-correlated pairs were grouped hierarchically into meta-states. Subject-specific states (cluster centroids) were calculated for use in the transition energy calculations described in Section 2.3.

**Figure 1.**
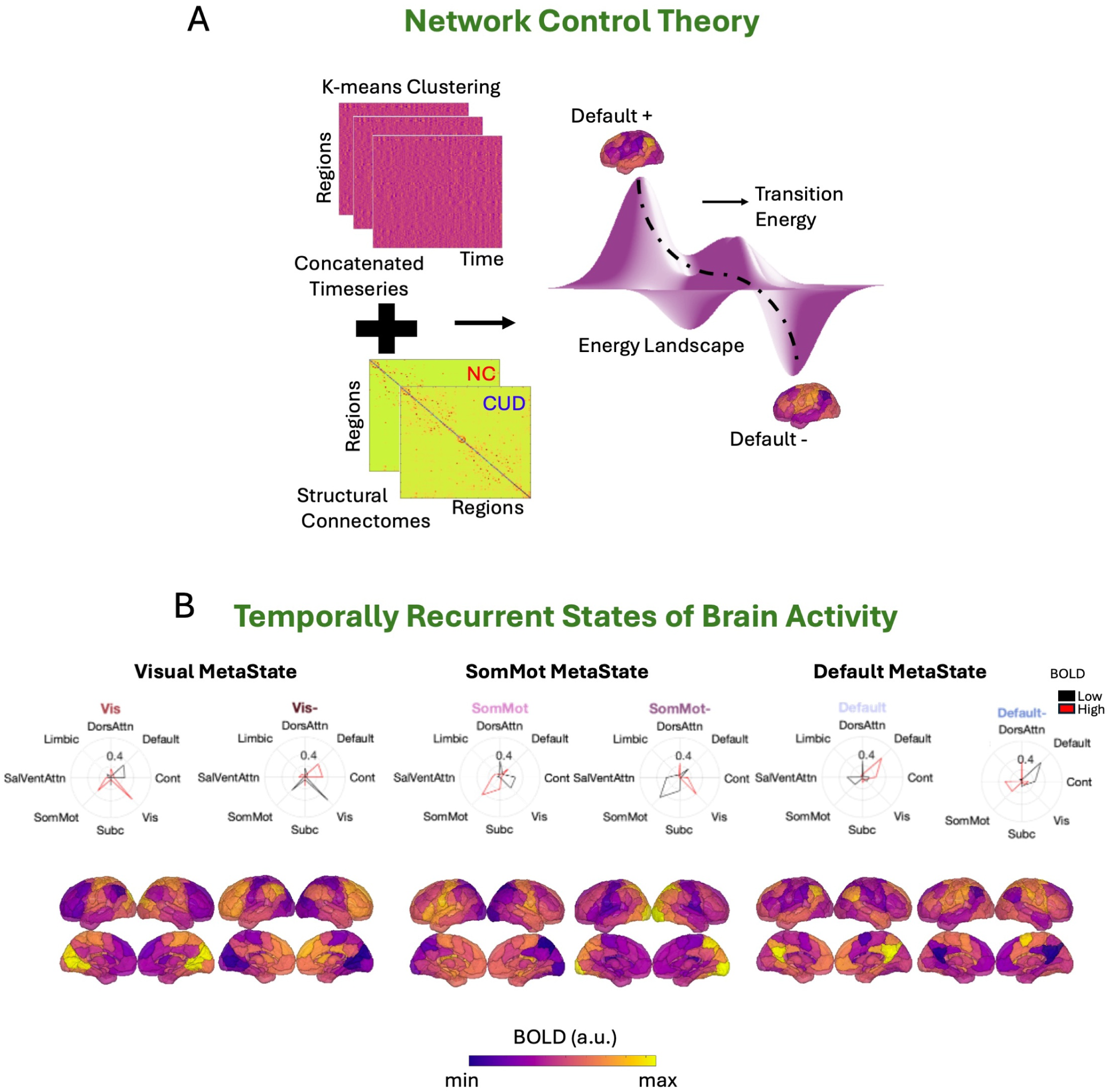
Recurrent States of Brain Activity. (**A**) K means clustering was conducted on regional fMRI time series data across all subjects (CUD and NC) to identify distinct brain activity states; an optimum of 6 brain states was identified. We subsequently utilized network control theory and group-specific structural connectomes (one for CUD and one for NC) to determine the energy required to complete state transitions. (**B**) Cosine similarity of each of the six states’ high-amplitude activity and low-amplitude activities to the Yeo resting-state networks (1 for regions in the network of interest and 0 elsewhere). Metastates (pairs of states with opposing activation) were hierarchically organized and consisted of the visual, somatomotor and default mode networks. The states are named based on the minimum distance to the Yeo canonical states.

Temporal dynamics of brain states were calculated, including fractional occupancy (FO), the number of TRs assigned to each cluster divided by the total number of TRs, dwell time, the average length of time spent in a cluster once transitioning to it and appearance rate, the total number of times a state was transitioned into per minute^19^ . Transition probabilities were also computed between each pair of ordered states, in addition to persistence probability (likelihood of staying in a state when currently in that state).

### 2.3 Network Control Theory and transition energy

As described in detail previously, we use a linear, time-invariant, controlled diffusion model of brain activity:

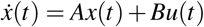

A is a representative group average NxN SC matrix (N = 116, one for NC and one for CUD), x is regional activity at time t, B is a NxN matrix of control input weights, and u(t) is the input. Group average SC matrices were used as dMRI data were not available for all subjects; global TE for individual level SCs and the group level SCs were highly correlated (r = 0.80, 1.0989*10 ^-15^). We used a time horizon of T = 1, with the strongest negative correlation for state-pair TP measures (across range T = 0.001 to 10), ensuring the least probable transition and largest TE^34,35^. By integrating u(t) over the time horizon for a specific transition, we determined total input needed in each region for state transitions. Summing regional inputs gives us whole brain TE. Doing so for ordered pair of states yields matrix of whole brain transition energies (TE); with the sum of entries in this k x k matrix being global TE. Ordering states by functional hierarchies (low to high), the upper triangular part of the pairwise TE matrix represents bottom-up transitions, and the lower triangular part of the matrix represents top-down transitions. Summing the pairwise TEs for each region gives us regional TEs, and summing regional TE values in each network gives us Network TEs. The effect of CUD-related changes in SC independent of any changes in fMRI dynamics is ascertained by using individual SCs (n = 32 NCs and 33 people with CUD) to compute TE between canonical activation states, i.e. binary state vectors of functional network (Yeo) as done previously^17,20^.

### 2.4 Statistical analyses

ANCOVA models including age, sex, tobacco use, mean frame-wise displacement during the fMRI, and IGT performance as covariates were used for group comparisons. Our primary ANCOVA sought to identify group differences (NC vs CUD) while the secondary ANCOVA tested the impact of length of CUD use normalized by age (analysis using CUD individuals only, median use = 10 years, interquartile range = 4-15 years). The full ANCOVA results are reported in the supplementary information. Unpaired post hoc t-tests were performed to determine the directionality of observed group effects, while post-hoc Spearman rank correlations were performed to determine the direction of the relationship between continuous variables. All p-values were corrected for multiple comparisons using the Benjamini-Hochberg false discovery rate (pFDR) procedure^36^.

### 2.5 NeuroMaps Analysis

To link regional TE effects (CUD vs NC differences and associations between TE and length of CUD) to nine different neurotransmitter systems, we used NeuroMaps, a toolbox that contains maps of receptor and transporter distributions from whole-brain, three-dimensional normative positron emission tomography (PET) atlases. The PET data used to create the atlases were collected from a total of 1,238 individuals (718 males and 520 females) across a variety of studies^24^. We computed Spearman rank correlations between regional TE effects (CUD vs NC differences and associations between TE and length of CUD) and receptor/transporter concentration levels for all 19 different receptor families available on neuromaps i.e. Acetylcholine - M_1_, *α*_4_*β*_2_, and VAChT, Cannabinoid - CB_1_, Dopamine - D_1_, D_2_, and DAT, GABA - GABA*_A/BZ_*, Glutamate - NMDA and mGluR_5_, Histamine - H_3_, Norepinephrine - NOR, Opioid - MOR, Serotonin - 5HT_1*A*,1*B*,2*A*,4,6_ and 5HTT.

## 3. Results

### 3.1 Brain (meta)states and their metrics

Recurrent states of brain activity were identified using k-means clustering and grouped into meta-states, each comprising two states with opposing activation patterns. Meta-states reflected opposing activation patterns of the visual, somatomotor, and DMN network, as seen in previous publications^20,21^. Fractional occupancy or dwell time did not differ between groups but appearance rates of the high amplitude somatomotor state in CUD were significantly higher (Supp Fig. S2). TP between pairs of states did not show significant CUD group differences, see Supp Fig. S4. However, there was one significant interaction term (CUD by IGT performance) for TP between low amplitude somatomotor state and high amplitude somatomotor states; post hoc tests revealed significant positive correlations between this TP and IGT performance in CUD, see Fig. S5. Females having CUD showed significantly increased global TE, but this finding is complicated by the low number of females in the study. (Supp Fig. S7).

**Table 1.**
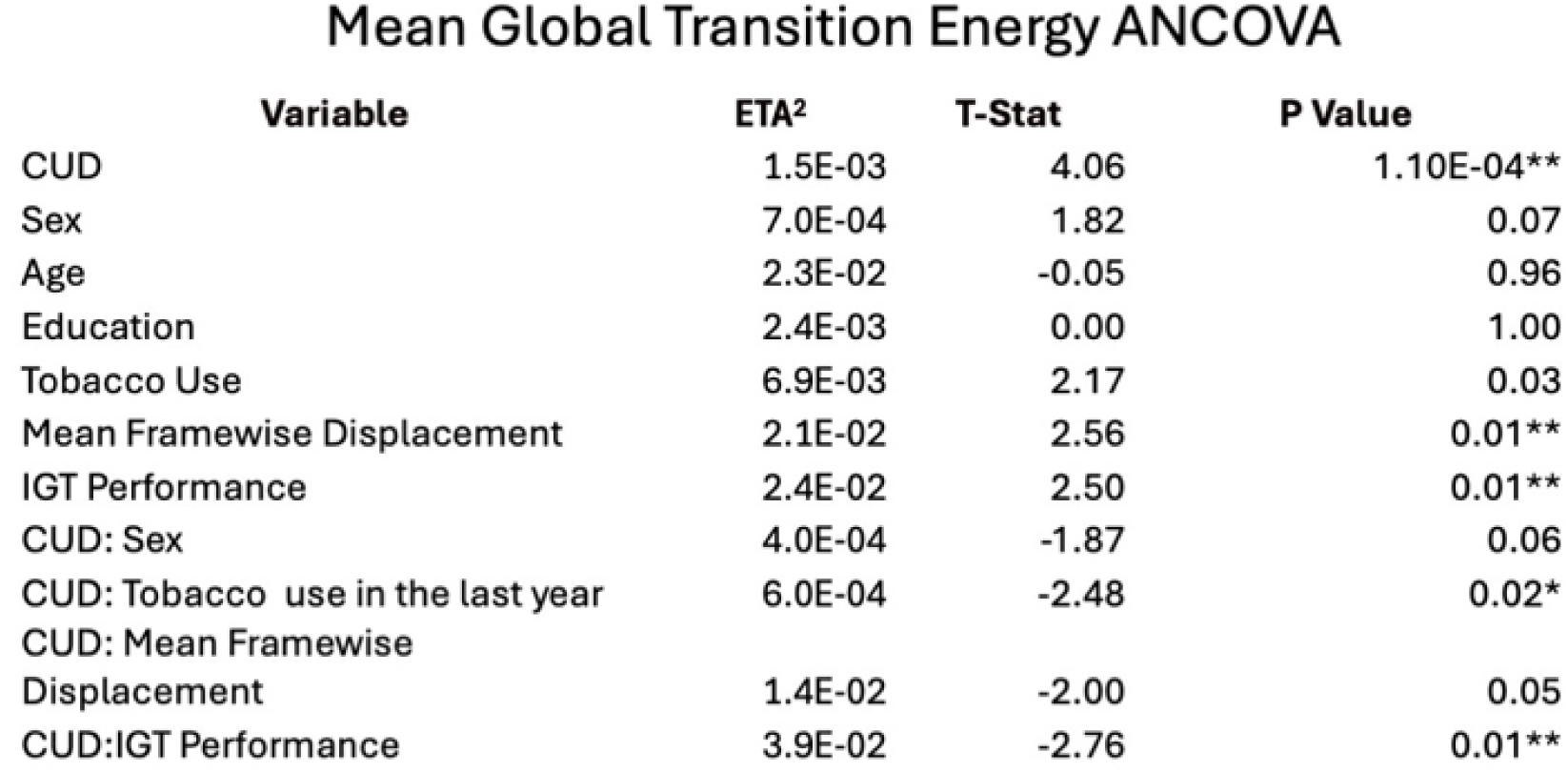
ANCOVA results showing effect sizes and t-stats for global transition energies reveal that people with CUD have lower global transition energies compared to NC. We find that both in-scanner motion (framewise displacement) and the predisposition for risky behaviors, measured using performance on the Iowa gambling task, are associated with global transition energies. Post-hoc testing revealed significant, positive correlations between IGT performance and global TE in CUD only (not NC). **corrected p < 0.05.

### 3.2 People with CUD have lower global and top-down transition energies that associate with risk-taking behaviors

For global and top-down TE, ANCOVAs showed significantly lower values in CUD compared to NCs (Table 1, Fig. S6), with smaller but significant effects of frame-wise displacement, IGT performance, and the interactions of IGT by CUD diagnosis and CUD diagnosis by tobacco use. A linear model was fit and all terms subtracted except for CUD group membership (linear model coefficient for CUD group was *β* = *−*34.1, CI = -17.4 to -50.8); marginalized residual histogram shown in Figure 2A. Frame-wise displacement was not significantly higher in CUD and global TE was not correlated with in-scanner motion (r = -0.0901, p = 0.3037). Pairwise state TE was lower in individuals with CUD between visual and DMN meta-states, and to/from the somatomotor states to the visual states, see Fig. 2B. When TE was averaged across top-down (lower triangular part of the TE matrix) or bottom-up (upper triangular part) transitions, both were lower for CUD and had similar significance patterns of associations compared to global TE, see Supplementary Table 2. Post-hoc testing revealed a positive, significant correlation between IGT performance and global and top-down TE for individuals with CUD (not NCs), see Figure 2C, D and E.

### 3.3 People with CUD have lower TE in higher-to-mid level networks, and in regions with excitatory receptors

Individuals with CUD exhibited significantly lower dorsal attention TE, and trends for lower TE in the DMN and limbic networks (see Fig. 3A). The interaction term of CUD group and IGT performance was significant after corrections in dorsal attention network TE, with post hoc tests revealing a trend toward a significant positive correlation for CUD only (see Fig. 3B). Regional TEs were significantly reduced primarily in frontal and dorsal cortical regions (see Fig. 4A), and the largest group difference effects were in regions with receptor expression of the noradrenergic and opioid systems (see Fig. 4B). Regions with higher concentrations of opioid receptors have larger decreases in TE in CUD (positive t-stats), while regions with high noradrenergic receptor concentrations have higher TE in CUD vs NC (negative t-stats).

### 3.4 Longer CUD duration relates to lower global, top-down, and excitatory system TE

We found that global TE, whole brain TE for top-down state transitions, and TE within the DMN, control and ventral attention networks were significantly negatively associated with years duration of CUD (normalized by age at the time of scanning) (Fig. 5A and Supplementary Table 5). We also investigated how various neurotransmitter receptor systems were associated with the regional t-stats for the association between duration of CUD and global TE (Fig. 5B). Regional t-stats capture spatial variations in the strength of the relationship between CUD duration and brain dynamics, allowing us to localize effects rather than relying solely on global averages and by aligning with specific neurotransmitter receptor distributions, provide mechanistic insight into altered brain state energetics. Our analysis reveals how regions with the strongest associations between duration of CUD and global TE are enriched most for D_1_ dopamine, H_3_ histamine, NMDA, GABA*_A/BZ_*inhibitory and glutamatergic receptors (mGluR5).

**Figure 2.**
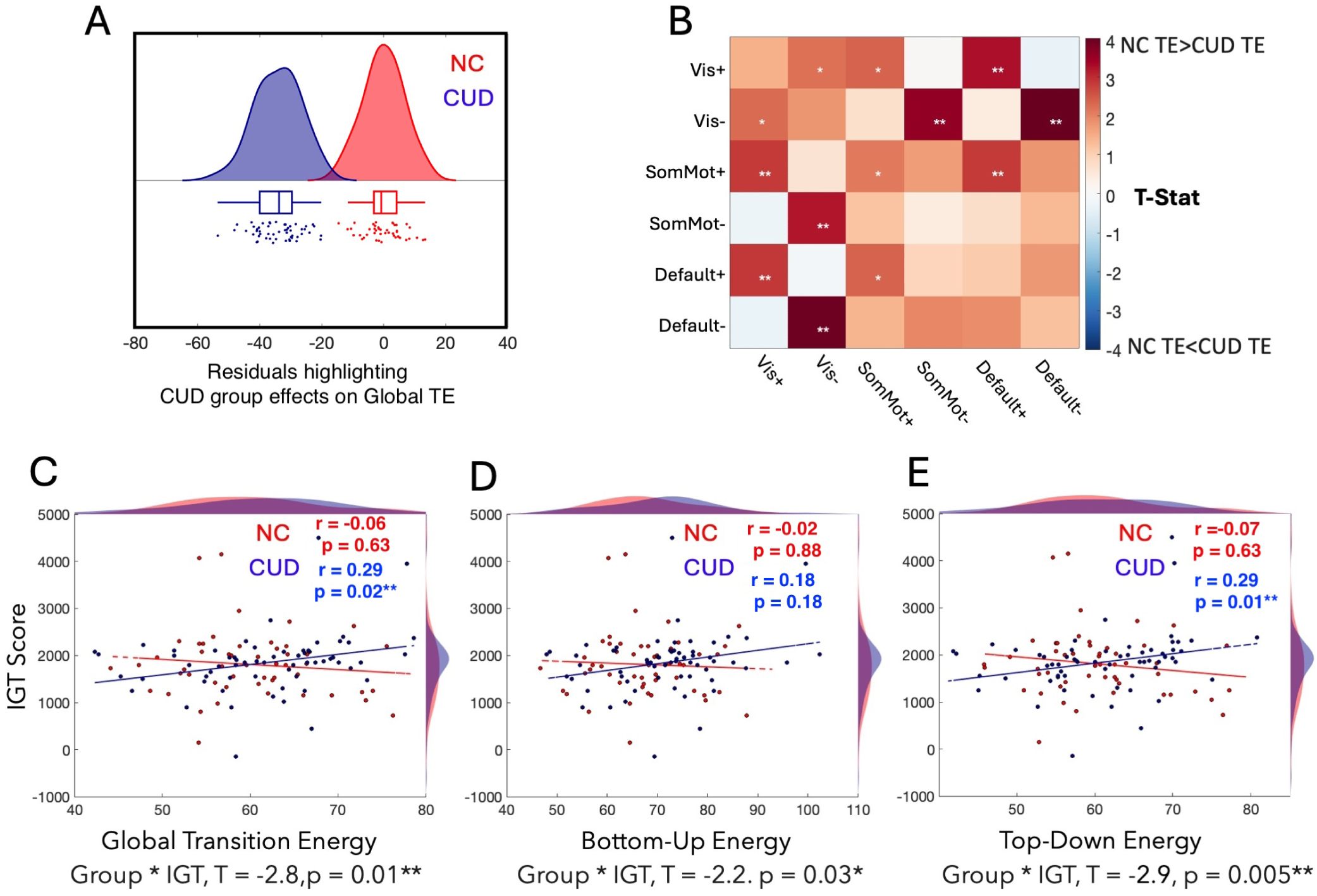
Brain state energy dynamics differ in people with Cocaine Use Disorder (CUD) and Non-User Controls (NC). (**A,B**) Histogram and boxplots of residuals showing CUD group effects on global transition energy. Heatmap of T-statistics contrasting pairwise transition energies, indicating where CUD requires less energy than NC to switch between brain states.. (**C,D,E**) Plots showing relationships between Iowa Gambling Task (IGT) performance and global (C), bottom-up (D), and top-down (E) transition energies, with group-specific regression lines (n = 132, 71 CUD). (pFDR <0.05**). *uncorrected p < 0.05, **corrected p < 0.05.

### 3.5 Structural connectome-based TE differences correlate with risk-taking behavior in people with CUD

We utilized individual level structural connectomes for those who had available data (n = 67, CUD = 33) and analyzed the energy needed to complete transitions between canonical functional resting states to isolate what TE shifts are driven by SC differences rather than functional activation pattern differences^20^. Our analysis found trends for CUD group-by-IGT performance interaction effect for TEs between the DMN network state and multiple other network states wherein the TE was correlated with IGT performance in individuals with CUD, but not in NCs (Fig. 6 and Supp Fig. S9). The CUD group-by-IGT performance interaction term for TE from the somatomotor network to the DMN network remained significant after correction. Specifically, individuals with CUD required significantly greater TE to perform better on the IGT task.

**Figure 3.**
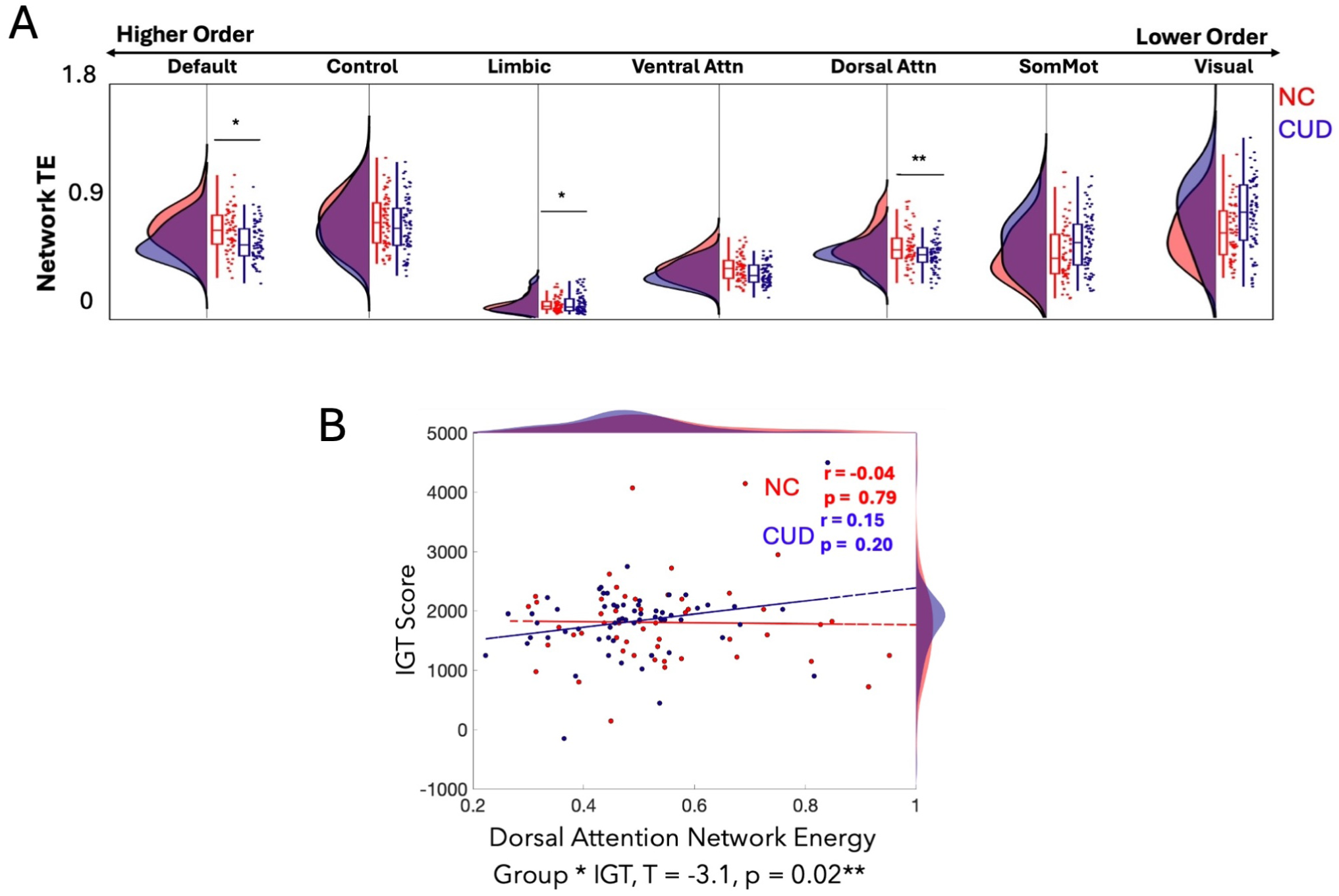
Network TEs are lower in people with CUD, and relate to risk-taking behaviors. (**A**) Network TEs were lower in people with CUD in the default mode network (pFDR <0.05, before correction), limbic and dorsal attention networks (pFDR <0.05, after correction) compared to NC. (**B**) Dorsal attention network TE showed a positive trend with performance on the IGT in people with CUD. p-values were corrected for multiple comparisons (Benjamini-Hochberg) where indicated. **corrected p < 0.05.

## 4. Discussion

In this work, we apply Network Control Theory to test for differences in brain dynamics of individuals with CUD compared to those without CUD using the SUDMEX-CONN dataset. Here we find that people with CUD exhibit whole-brain shifts in brain dynamics compared to those without CUD, specifically, they have lower global transition energy needed to complete transitions between recurring patterns of brain activity. The lower whole-brain transition energies appear to be driven by higher to mid-level networks (default, limbic, and dorsal attention), as well as regions that are enriched for excitatory receptors, whose overactivation has been shown to associate with the emergence of compulsive, drug-seeking behaviors typical of CUD. Lower global, top-down and dorsal attention transition energies are also correlated with poorer performance on a risk-taking behavioral task only in individuals with CUD. We interpret this finding as individuals with CUD expending more transition energy, particularly for top-down inhibitory transitions, in order to perform better on the gambling task - whereas NCs have no association, revealing a potential compensatory mechanism in CUD. Furthermore, there was a dose-dependence of this lower transition energy in CUD - the longer someone has CUD, the lower their global, top-down, and higher order transition energies. Overall, these findings reveal robust and widespread decreases in transition energy demand in people with CUD.

### 4.1 Whole brain energy dynamics are reduced in people with CUD

Our findings of the change in global and top down energy align with established evidence of cortical dysfunction in CUD and its comorbidity with other psychiatric disorders^9,14,37,38^. We also find higher and mid-level network involvement, with regions in the frontal lobe and medial surface particularly requiring significantly lower TE in CUD vs NCs. Areas like the dorsolateral prefrontal cortex or orbitofrontal cortex exhibit altered dynamics that could be attributed due to dopamine-mediated disruptions in striato-thalamo-cortical circuits, which contribute to the neuropathology of CUD^18,20^.

**Figure 4.**
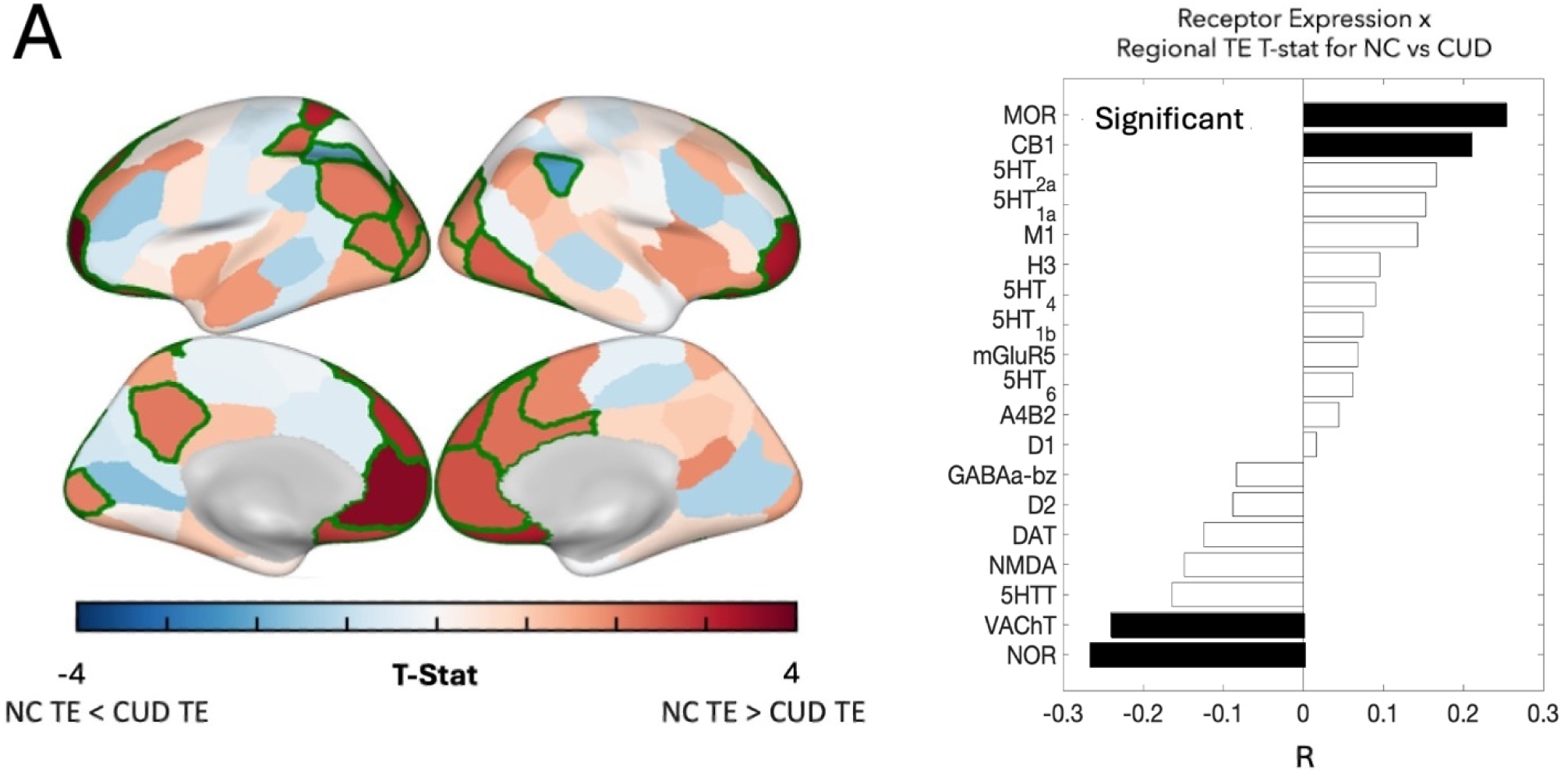
Regional TE. (**A**) ANCOVAs showed lower TEs in CUD vs NC for regions within higher order networks that are enriched for receptors like norepinephrine. Receptor families are categorized accordingly: Acetylcholine - M_1_, *α*_4_*β*_2_, and VAChT, Cannabinoid - CB_1_, Dopamine - D_1_, D_2_, and DAT, GABA - GABA*_A/BZ_*, Glutamate - NMDA and mGluR_5_, Histamine - H_3_, Norepinephrine - NOR, Opioid - MOR, Serotonin - 5HT_1*A*,1*B*,2*A*,4,6_ and 5HTT. p-values were corrected for multiple comparisons (Benjamini-Hochberg). *uncorrected p < 0.05, **corrected p < 0.05.

Our findings suggest that even in the absence of explicit task demands, individuals with CUD require less energy in dorsal, default mode and limbic systems to complete state transitions. These lower TEs could be attributed to differences that exist before CUD and predispose them to developing CUD, or are a result of CUD given that lower TEs correlate signficantly with duration of CUD. We observe lower energetic demands in the dorsal attention networks in particular. Damage to dopamine-rich motor circuitry, along with adjacent regions within the dorsal attention network that typically operate at higher metabolic energy levels, contributes substantially to the dysfunction induced by chronic cocaine use and its disruption of signaling dynamics^39,40^. These impairments affect a broad range of cognitive and motor functions^41–43^. Sensorimotor states that engage the dorsal attention network are known to require elevated metabolic energy during task performance in order to process sensory input and coordinate successful motor and cognitive execution^44^. Previous work in people with epilepsy revealed a possible correspondence in NCT-based TE and metabolic activity^45^. Here, we propose that the reduced metabolic energy demands observed in individuals with CUD within these circuits may translate to the lower transition energies we find, account for the deficits in task performance shown by people with CUD, and may also reveal a compensatory mechanism in relation to performance on a risk taking behavior task.

### 4.2 Length of CUD impacts energy dynamics

CUD duration was negatively correlated with global TE, particularly for energy to complete top-down transitions, as well as having a trend toward being negatively correlated with TE in control, default mode and salience/ventral attention networks. The fact that longer duration of CUD was related to lower TE may indicate that the lower TE found in CUD vs NC may more likely a result of cocaine use rather than a pre-existing mechanism existing before cocaine use. Networks like the salience and default mode networks are essential for suppressing drug-related urges and maintaining inhibitory control over reward-driven impulses^46,47^. Our results indicate that regional and global energy expenditures—particularly top-down signaling originating from frontal regions that suppress bottom-up drive toward drug use—are diminished in individuals with CUD. This reduction worsens with longer durations of cocaine use, possibly reflecting progressive neuronal damage to dopaminergic pathways and a shift toward intensified craving and increased drug consumption over time (Fig. 3)^48^.

Prolonged cocaine use has been associated with reduced dopamine reuptake in dopamine receptor-rich regions such as the ACC. The resulting excess of postsynaptic dopamine leads to receptor down-regulation, reduced metabolic activity, and impaired signaling across circuits involving these regions. In task-based fMRI studies, regions like the ACC demonstrate stronger deactivation in individuals with CUD relative to healthy controls—a consequence of long-term cocaine exposure^10^. Resting-state studies similarly implicate aberrant functional activity in these regions in disorders comorbid with addiction, such as depression and mania^49^. The reduction in functional activation in these areas during task performance has been linked to diminished craving, which is necessary for improved behavioral control in individuals with CUD. Our findings of lower transition energies in these same regions underscore the regional impact of CUD and offer insight into the structural and functional alterations that impair frontal circuits responsible for regulating craving-related behaviors central to the pathology of CUD^50,51^. This link between task-based adaptations and altered resting-state control dynamics is further corroborated by recent findings showing that artificially perturbed state transitions in individuals CUD reveal persistent energy inefficiencies reflective of task-induced plasticity—that is, individuals with CUD rely on less efficient neural pathways for the same state transitions as NCs, highlighting the enduring impact of cocaine-induced neuroplasticity^52^.

**Figure 5.**
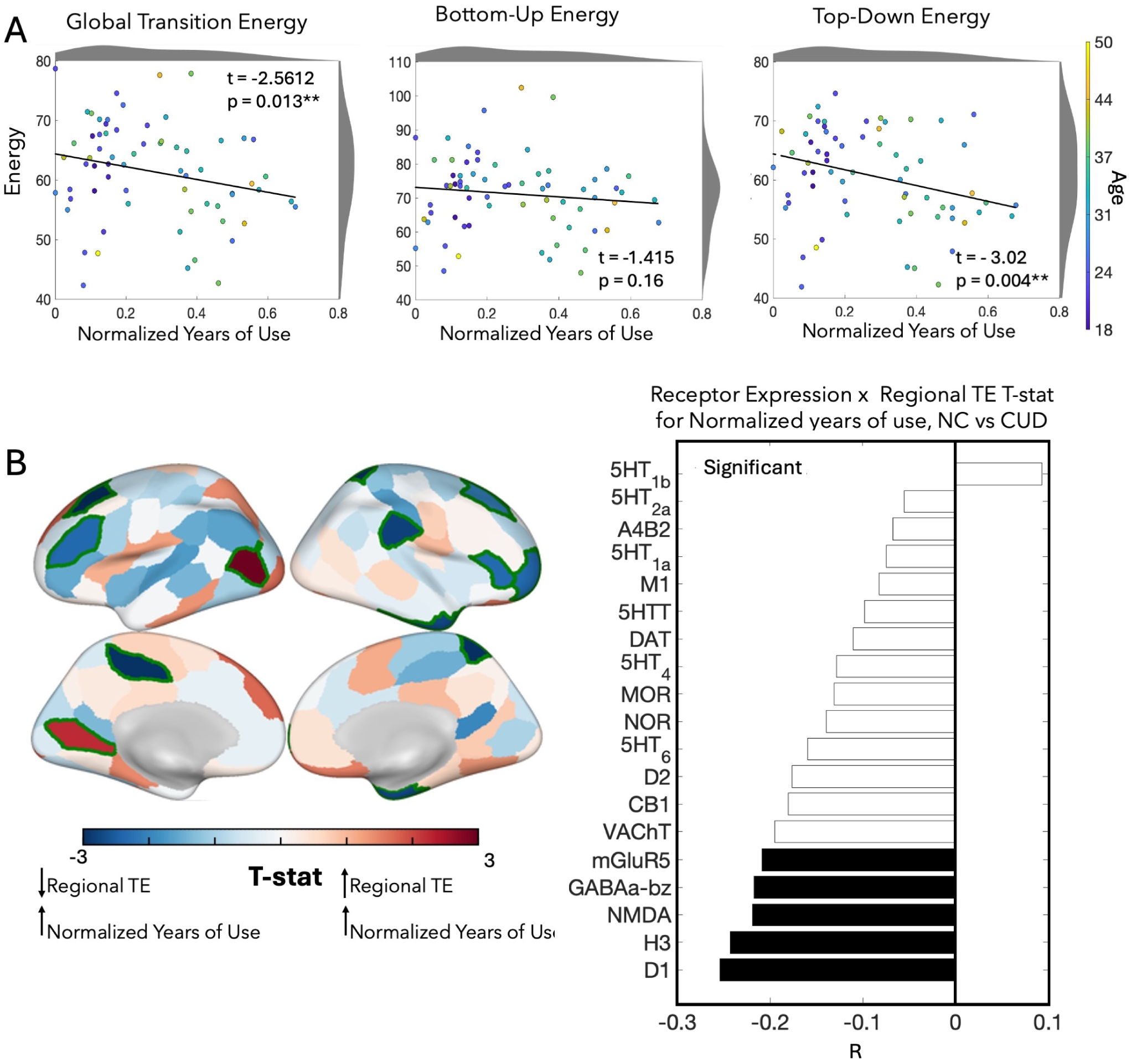
Global and top-down TE decreases with length of cocaine use, and correlate with regional differences in the expression of receptor families. (**A**) All TEs were negatively related to length of CUD, with global TE and top down TE having significant relationships after correction. (**B**) Some regional TEs had trends toward being positively or negatively associated with normalized years of use (regions with green outline; p < 0.05 before correction). Regional TE associations with length of use were correlated with various receptor/transporter concentrations and the largest overlap was with dopaminergic receptor maps whose dysfunction is implicated in the pathology central to cocaine use disorder. p-values were corrected for multiple comparisons (Benjamini-Hochberg) where indicated. **corrected p < 0.05.

**Figure 6.**
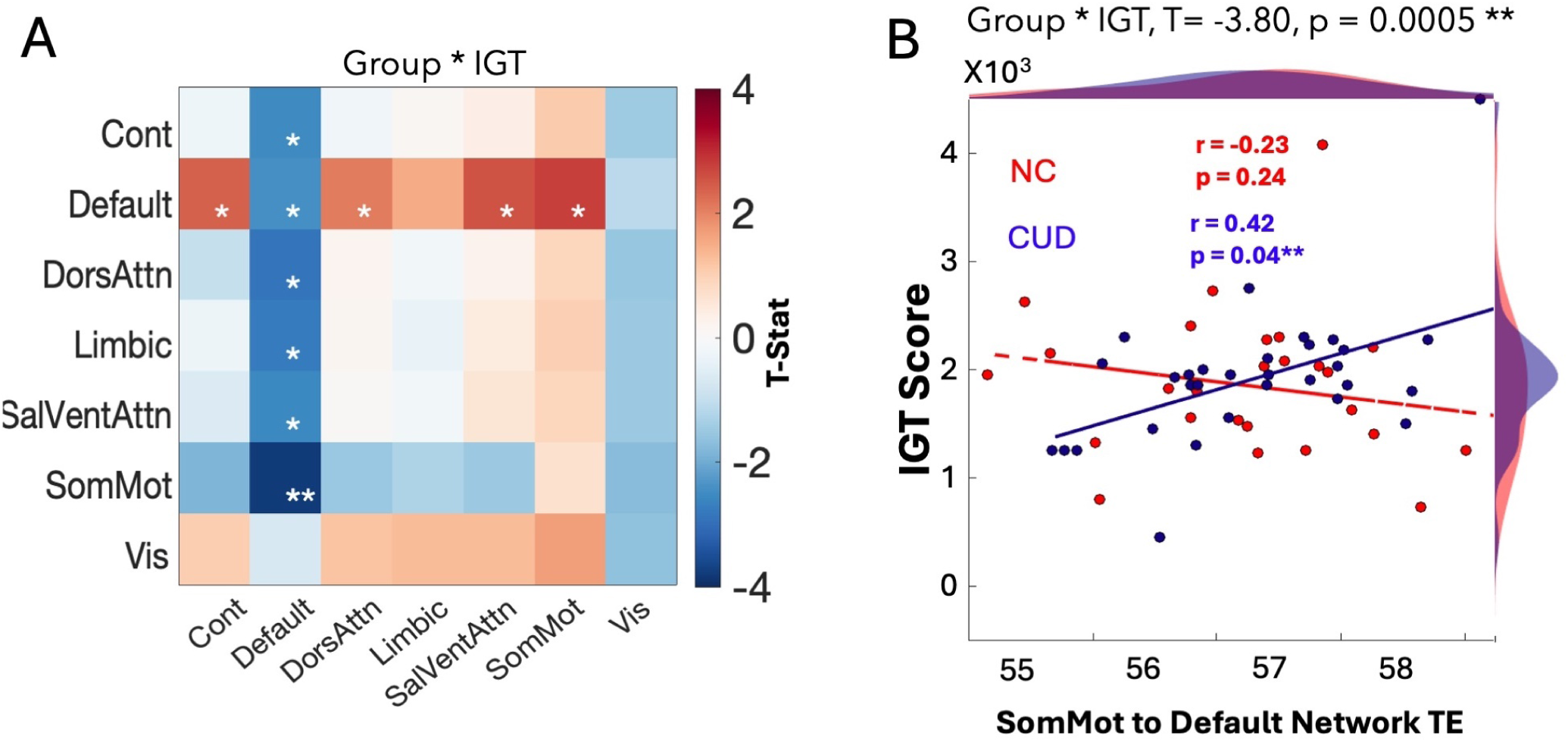
Canonical State Analysis. A subgroup analysis of individual-level SCs and canonical functional states showed that TE to/from the default mode network state had significant (before corrections) interaction effects of CUD group and performance on the Iowa Gambling task (IGT). One transition from somatomotor to default mode network had a significant interaction effect after correction wherein there was a significantly positive correlation of TE and IGT performance in CUD and not in NCs (CUD Group * IGT, t = -3.80, corrected p < 0.05). *uncorrected p < 0.05 and **corrected p < 0.05.

### 4.3 Increased energy demands link to better performance on tasks measuring risky behaviors

Our findings indicate that individuals with CUD having higher global TE, particularly for dorsal attention networks, perform better on the Iowa Gambling Task (IGT). We show that dynamics are particularly disrupted in part of an interhemispheric dorsal frontoparietal network that is central to top-down attention control and behavioral regulation relevant to craving^46^. Cocaine users consistently show altered activity in this dorsal attention network, possibly reflecting a maladaptive shift in attentional allocation. This dysfunction manifests behaviorally as poor performance on tasks requiring fine motor coordination, sustained attention, and real-time sensory feedback—capacities critical for inhibiting craving-related distractions^53–55^. Moreover, these performance deficits appear to worsen with longer duration of CUD, paralleling the progressive decrease in TE. Here we find that the dorsal attention network — along with structural connectivity differences that alter TE dynamics between somatomotor and default mode networks — appears to mitigate this known deficit in risk-related behavior associated with CUD (Fig. 3, 6)^56^.

### 4.4 Changes in transition energy relate to exctitaory receptors that cause glutamatergic excitotoxicity

Previous work identifying differences in neural systems between people with CUD and NCs shows a degradation of reward circuitry, particularly in regions characterized by noradrenergic and glutamatergic signaling^57–61^. These receptors play a crucial role in arousal, attention, stress response, and reward processing—functions that intersect significantly with the neuropathology of cocaine use. The decline in energetic expenditures across the course of CUD is closely linked to excitotoxic cell death in receptor systems such as dopamine and noradrenaline^58,59,62^. We demonstrate that changes in glutamatergic, dopaminergic, noradrenergic and serotonergic receptor expression significantly correlate with the association between lower regional TE and longer duration of CUD (Figs. 2, 3). These receptor-level alterations could reflect glutamatergic excitotoxicity, a mechanism central to the neurobiology of CUD^63^. As dopamine levels surge in response to cocaine intake, excitatory signaling in the prefrontal cortex and striatum becomes dysregulated. When this excitation exceeds physiological limits, it leads to overstimulation, circuit degradation, and ultimately, neuronal death. We posit that neuronal death that presumably increases with the duration of CUD, could be driving the regional decreases in energetic expenditures in these systems.

The neurodegenerative progression alters the system’s energetic demands, providing a mechanistic explanation for the shift from recreational use to compulsive drug-seeking behavior^7^. Once neuronal damage sets in, higher levels of energy are required for individuals with CUD to achieve cognitive performance comparable to that of healthy controls^64,65^. This finding may illuminate the pathological alterations in brain energy landscapes and underscore the compensatory processes necessary to preserve cognitive function in the face of chronic cocaine exposure.

### 4.5 Limitations

This analysis is subject to several limitations. The dataset used in this study includes individuals from a single population—Mexican adults—and is predominantly composed of male participants aged 18–50. As a result, our findings are geographically and demographically specific, limiting generalizability and precluding any analysis of potential sex-specific differences in energy dynamics. There was a suggestion of potential sex-mediated effects of CUD on global TE but future work with larger sample sizes are needed before any claims can be made. Although we observe an association between transition energy and the duration of CUD (normalized by age), the restricted age range prevents us from drawing conclusions about the long-term progression of CUD beyond the age of 50. Additionally, the dataset includes mostly individuals who consume cocaine in the form of crack cocaine, thereby constraining our ability to examine energy dynamics across varied forms of ingestion. Expanding these findings will require inclusion of recreational cocaine users and a more demographically diverse sample, spanning a broader range of ages and balanced sex representation.

### 4.6 Conclusion

Our results show that individuals with CUD, relative to healthy controls, exhibit reduced network control–based energetic demand globally and for top-down brain state transitions, particularly within higher-to-mid-order networks. This decline scales with the duration of CUD, reflecting a pathological shift in the brain’s functional architecture that is associated with both decreased TE and poorer performance on risk-sensitive cognitive tasks, such as the Iowa Gambling Task (IGT). Interestingly, a possible compensatory mechanism may involve increased energy expenditure in preserved network regions, which could enable some individuals with CUD to partially maintain or improve IGT performance despite overall network dysfunction. These findings underscore the potential of network control theory to elucidate systems-level alterations underlying cocaine addiction. Further investigation into pharmacological, behavioral, or non-invasive brain stimulation interventions targeting these disrupted energy dynamics may help optimize therapeutic strategies for individuals with CUD.

## Supporting information

Supplemental Image 1

Supplemental Image 2

Supplemental Image 3

Supplemental Image 4

Supplemental Image 5

Supplemental Image 6

Supplemental Image 7

Supplemental Image 8

Supplemental Image 9

Supplemental Image 10

Supplemental Image 11

Supplemental Image 12

Supplemental Image 13

Supplemental Image 14

Supplemental Image 15

Supplemental Image 16

Supplemental Image 17

Supplemental Image 18

## 4.7 Acknowledgements

We thank Prof Mallar Chakravarty (Director of the Computational Neuroanatomy Laboratory, Douglas Research Centre, Montreal, Canada) who provided access to computational tools from his group on the Niagara Compute Cluster from Compute Canada.

---

**Figure S1.**
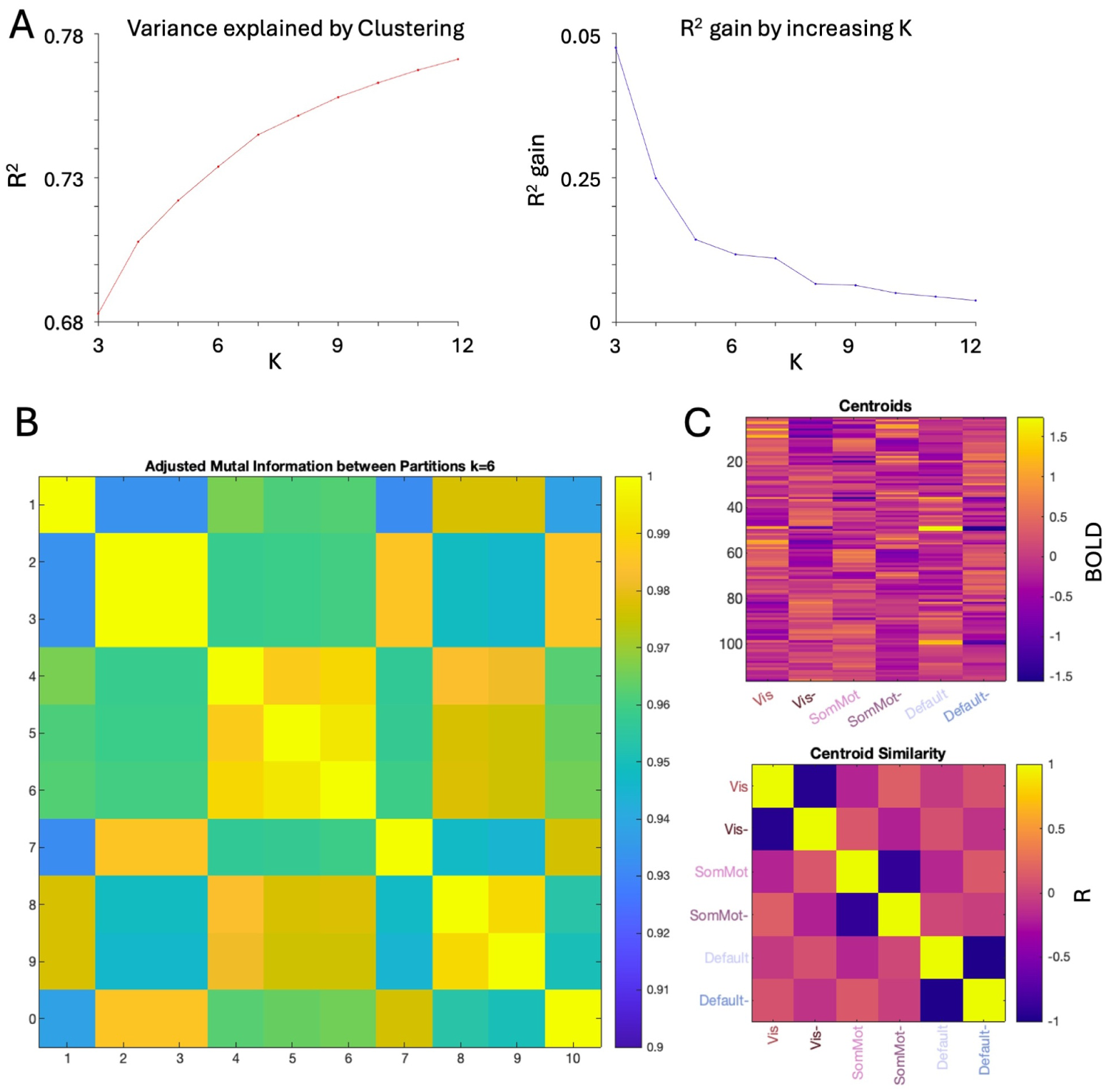
Determining k and cluster stability. (A) 50 repetitions of k-means clustering were done for a range of k from k = 2 to k = 14. Variance was quantified as the ratio of between cluster variance to complete variance. We plotted variance gained by increasing k, and observed a plateauing around k = 6. We analyzed results with k = 6, given that increase in explained variance after this k was negligible. (B) K-means clustering was repeated 50 times to find the lowest error solution. We sought to find a consistent global minimum through repeating k-means clustering 10 times, before comparing the adjusted mutual information (AMI) between the 10 partitions generated. Maximum sum of AMI score with all other partitions was used to guide the analysis, while allowing us to confirm that k-means clustering was consistent. Centroids produced were shown to have high similarity.

**Figure S2.**
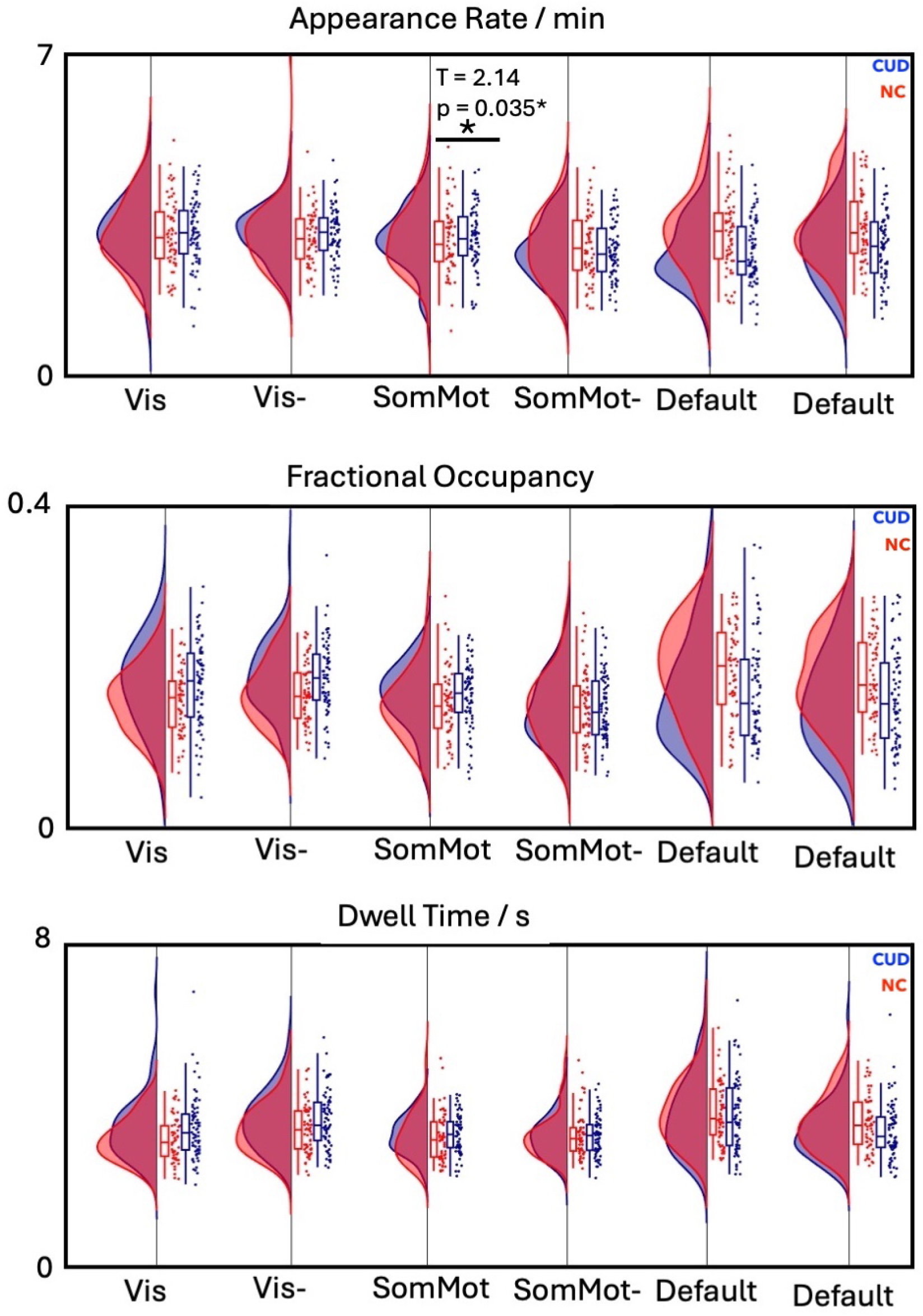
We calculated appearance rate, fractional occupancy, and dwell time. We observed that the somatomotor state had higher appearances in people with CUD before correction and found no significant differences between other metrics.

**Figure S3.**
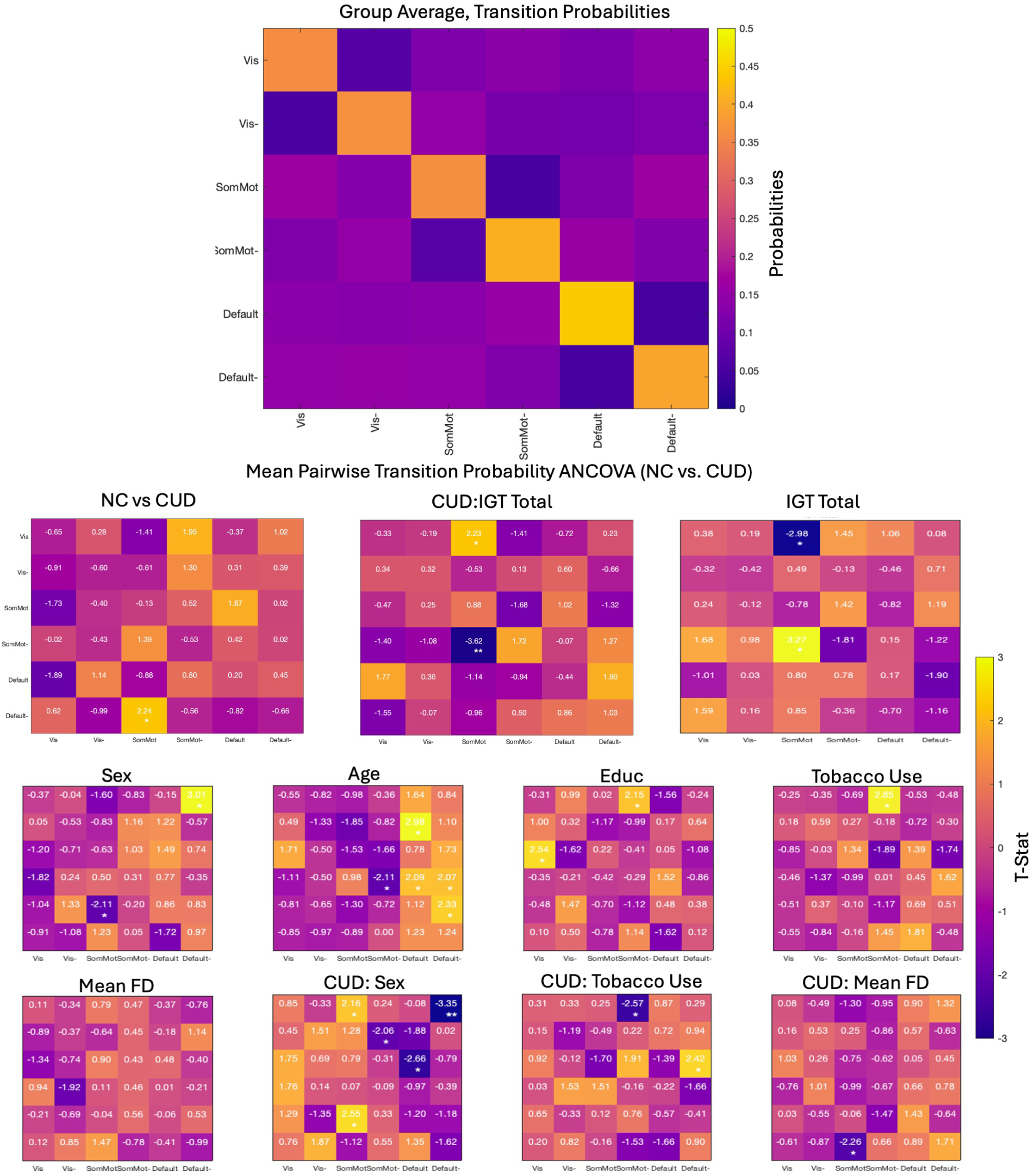
ANCOVA for transition probabilities did not find differences that survived correction between people with CUD and non user controls, while controlling for other covariates.

**Figure S4.**
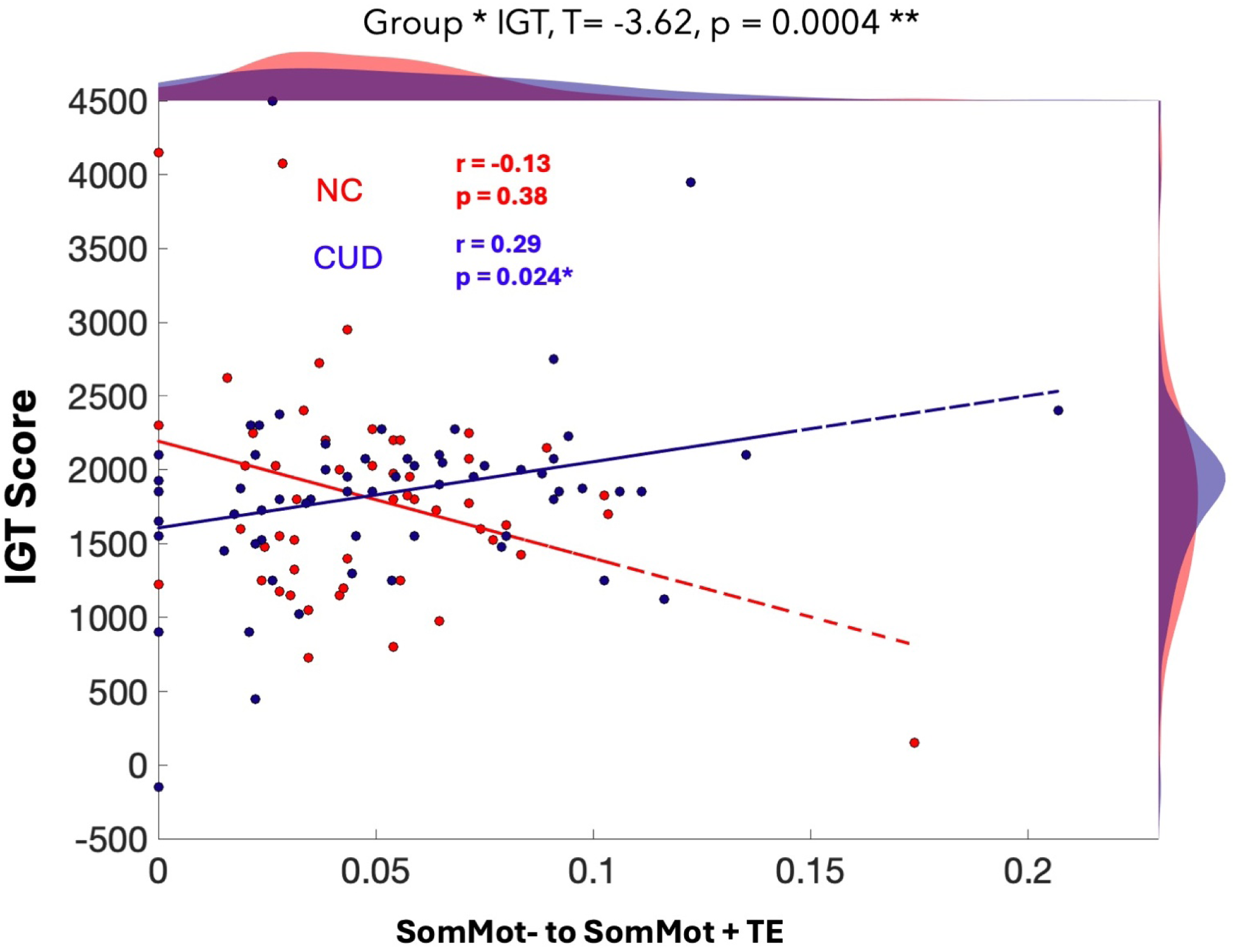
Post hoc testing for transition probabilities revealed a significant interaction between risky behaviors and transition probabilities for the transition from the SomMot low amplitude to the SomMot high amplitude state. *uncorrected p < 0.05, **corrected p < 0.05..

**Figure S5.**
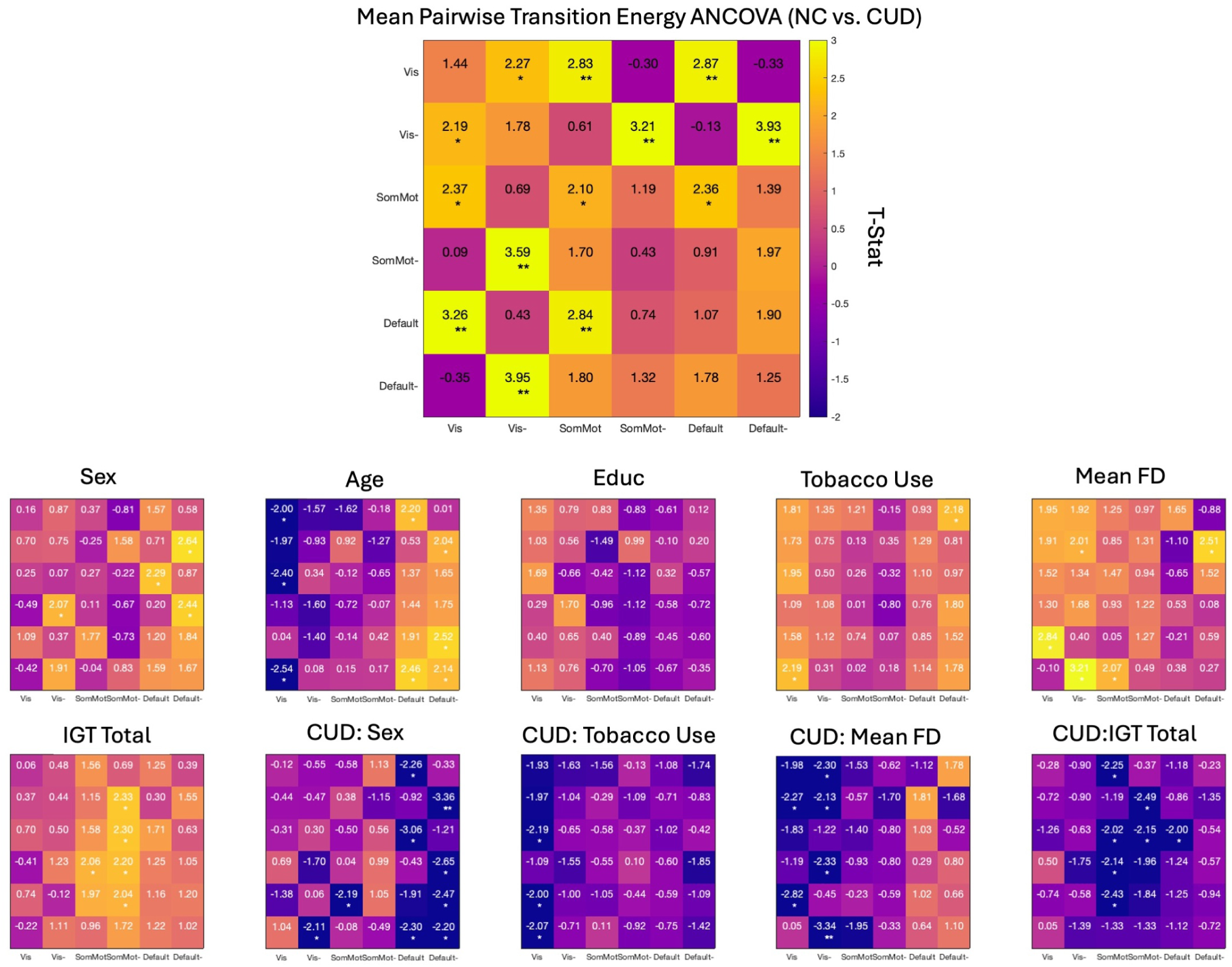
ANCOVA for comparing transition energies between groups indicated that the energetic landscape was lower for people with CUD, with these effects primarily being driven by the transition energies involving transitions to and from the default to the visual network. The impact of covariates, particularly performance on IGT scores, underscored the pathology of CUD and how it impacts risk taking social behaviors.

**Table 2.**
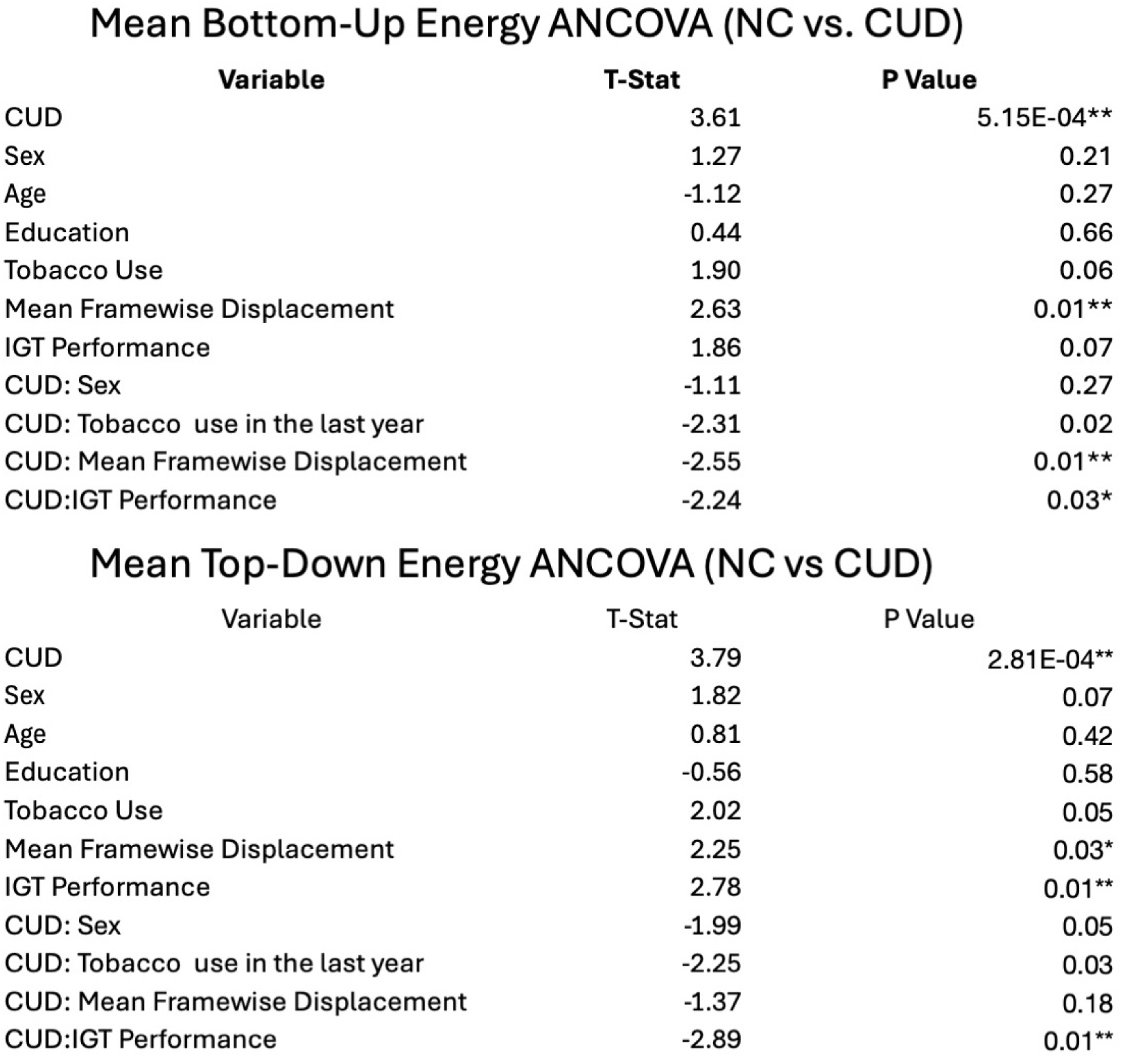
ANCOVA for comparing transition energies on a top down and bottom up balance of signaling.

Mean Bottom-Up Energy ANCOVA (NC vs. CUD)
| Variable | T-Stat | P Value |
| --- | --- | --- |
| CUD | 3.61 | 5.15E-04** |
| Sex | 1.27 | 0.21 |
| Age | -1.12 | 0.27 |
| Education | 0.44 | 0.66 |
| Tobacco Use | 1.90 | 0.06 |
| Mean Framewise Displacement | 2.63 | 0.01** |
| IGT Performance | 1.86 | 0.07 |
| CUD: Sex | -1.11 | 0.27 |
| CUD: Tobacco use in the last year | -2.31 | 0.02 |
| CUD: Mean Framewise Displacement | -2.55 | 0.01** |
| CUD:IGT Performance | -2.24 | 0.03* |

| Variable | T-Stat | P Value |
| --- | --- | --- |
| CUD | 3.79 | 2.81E-04** |
| Sex | 1.82 | 0.07 |
| Age | 0.81 | 0.42 |
| Education | -0.56 | 0.58 |
| Tobacco Use | 2.02 | 0.05 |
| Mean Framewise Displacement | 2.25 | 0.03* |
| IGT Performance | 2.78 | 0.01** |
| CUD: Sex | -1.99 | 0.05 |
| CUD: Tobacco use in the last year | -2.25 | 0.03 |
| CUD: Mean Framewise Displacement | -1.37 | 0.18 |
| CUD:IGT Performance | -2.89 | 0.01** |

**Figure S6.**
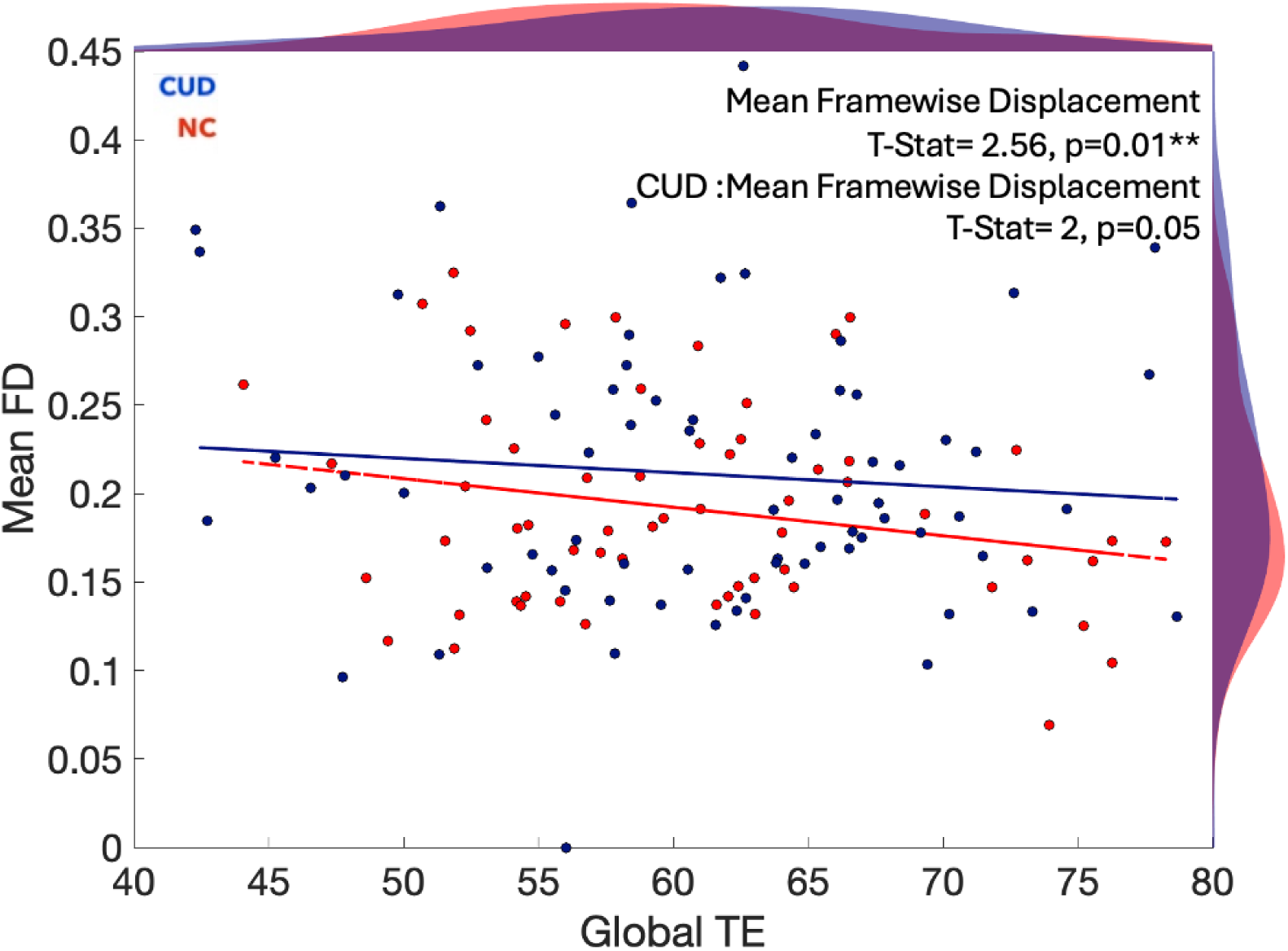
Global transition energies are impacted by mean framewise displacement across both groups

**Figure S7.**
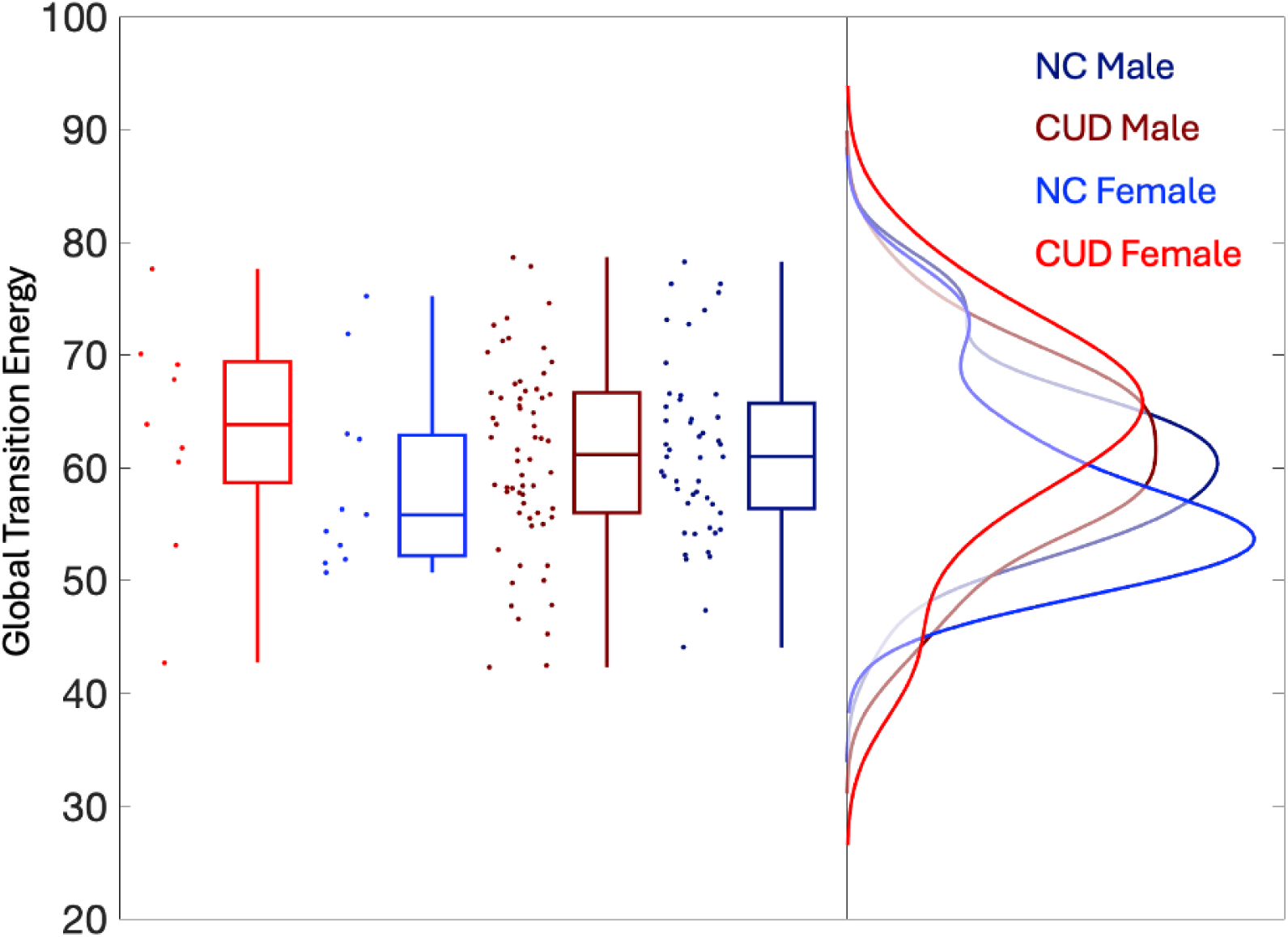
Global transition energies are impacted by sex, but this result is hampered by the low number of females in this cohort

**Table 3.**
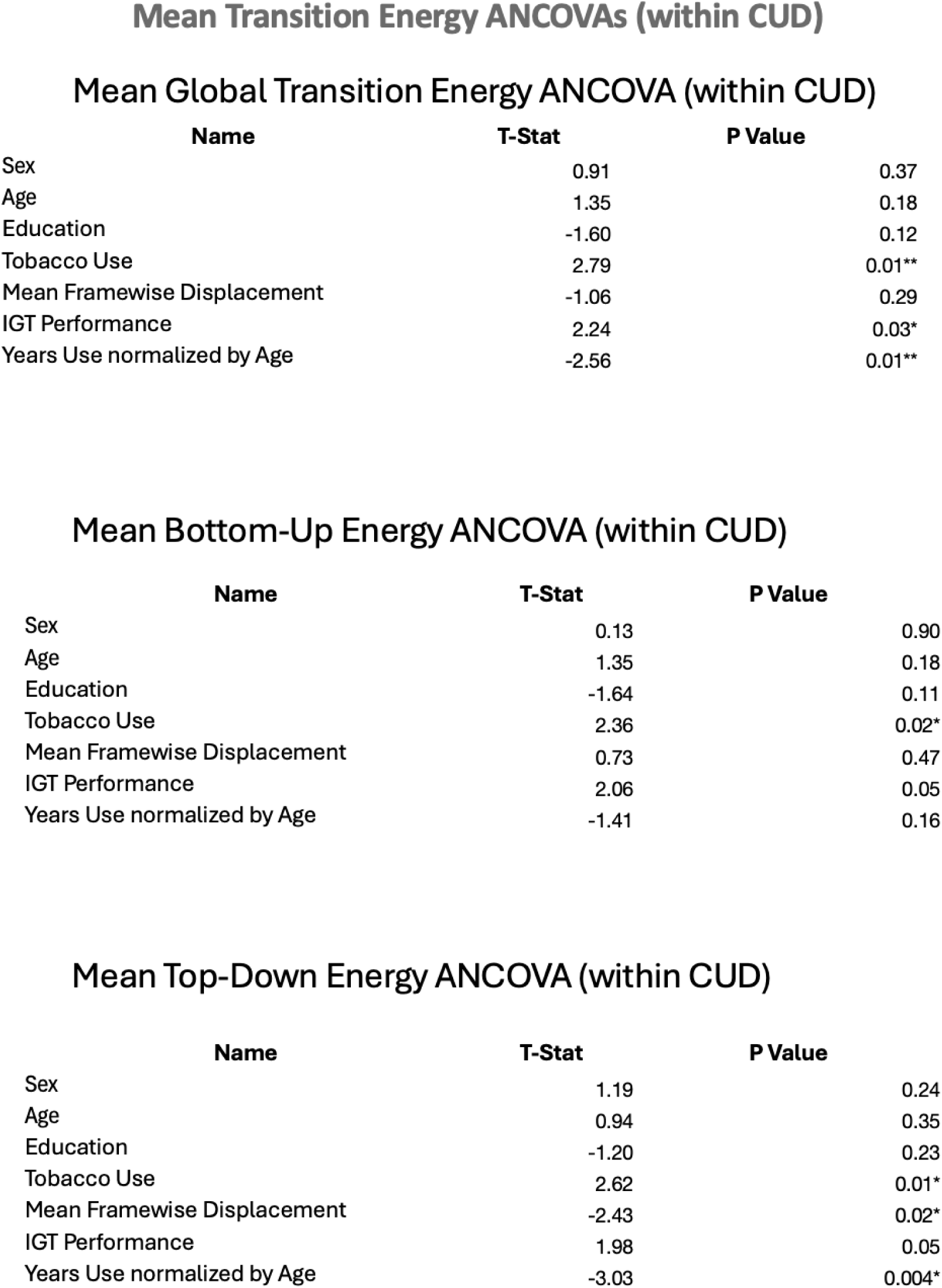
ANCOVA for understanding the impact of duration of cocaine use, normalized by age on global, top-down and bottom-up TE.

Mean Global Transition Energy ANCOVA (within CUD)
|  | Name | T-Stat | P Value |
| --- | --- | --- | --- |
| Sex |  | 0.91 | 0.37 |
| Age |  | 1.35 | 0.18 |
| Education |  | -1.60 | 0.12 |
| Tobacco Use |  | 2.79 | 0.01** |
| Mean Framewise Displacement |  | -1.06 | 0.29 |
| IGT Performance |  | 2.24 | 0.03* |
| Years Use normalized by Age |  | -2.56 | 0.01** |

Mean Bottom-Up Energy ANCOVA (within CUD)
|  | Name | T-Stat | P Value |
| --- | --- | --- | --- |
| Sex |  | 0.13 | 0.90 |
| Age |  | 1.35 | 0.18 |
| Education |  | -1.64 | 0.11 |
| Tobacco Use |  | 2.36 | 0.02* |
| Mean Framewise Displacement |  | 0.73 | 0.47 |
| IGT Performance |  | 2.06 | 0.05 |
| Years Use normalized by Age |  | -1.41 | 0.16 |

|  | Name | T-Stat | P Value |
| --- | --- | --- | --- |
| Sex |  | 1.19 | 0.24 |
| Age |  | 0.94 | 0.35 |
| Education |  | -1.20 | 0.23 |
| Tobacco Use |  | 2.62 | 0.01* |
| Mean Framewise Displacement |  | -2.43 | 0.02* |
| IGT Performance |  | 1.98 | 0.05 |
| Years Use normalized by Age |  | -3.03 | 0.004* |

**Table 4.**
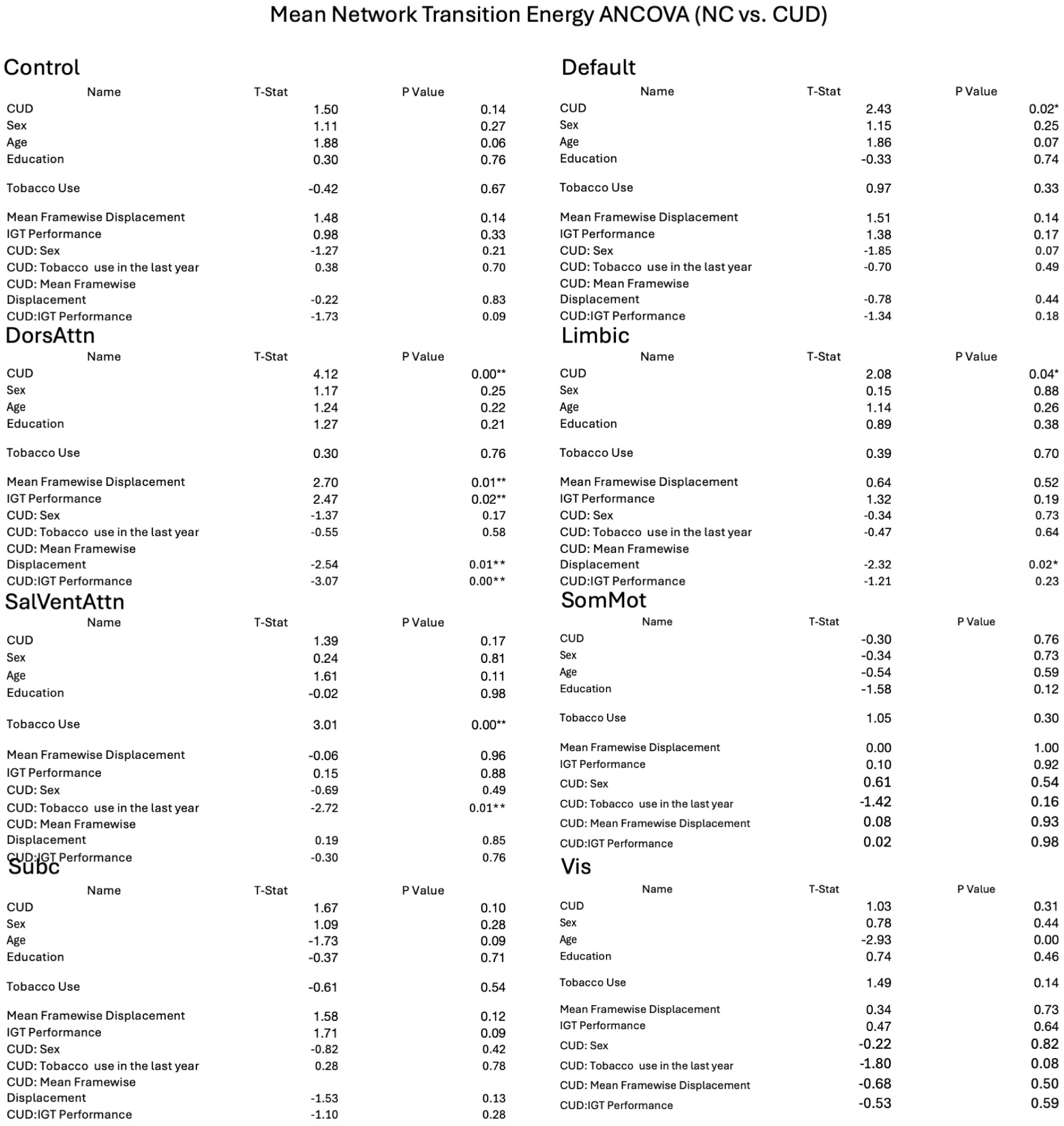
ANCOVA for comparing transition energies between groups for Yeo Networks.

Control
| Name | T-Stat | P Value |
| --- | --- | --- |
| CUD | 1.50 | 0.14 |
| Sex | 1.11 | 0.27 |
| Age | 1.88 | 0.06 |
| Education | 0.30 | 0.76 |
| Tobacco Use | -0.42 | 0.67 |
| Mean Framewise Displacement | 1.48 | 0.14 |
| IGT Performance | 0.98 | 0.33 |
| CUD: Sex | -1.27 | 0.21 |
| CUD: Tobacco use in the last year | 0.38 | 0.70 |
| CUD: Mean Framewise Displacement | -0.22 | 0.83 |
| CUD:IGT Performance | -1.73 | 0.09 |

DorsAttn
| Name | T-Stat | P Value |
| --- | --- | --- |
| CUD | 4.12 | 0.00** |
| Sex | 1.17 | 0.25 |
| Age | 1.24 | 0.22 |
| Education | 1.27 | 0.21 |
| Tobacco Use | 0.30 | 0.76 |
| Mean Framewise Displacement | 2.70 | 0.01** |
| IGT Performance | 2.47 | 0.02** |
| CUD: Sex | -1.37 | 0.17 |
| CUD: Tobacco use in the last year | -0.55 | 0.58 |
| CUD: Mean Framewise Displacement | -2.54 | 0.01** |
| CUD:IGT Performance | -3.07 | 0.00** |

SalVentAttn
| Name | T-Stat | P Value |
| --- | --- | --- |
| CUD | 1.39 | 0.17 |
| Sex | 0.24 | 0.81 |
| Age | 1.61 | 0.11 |
| Education | -0.02 | 0.98 |
| Tobacco Use | 3.01 | 0.00** |
| Mean Framewise Displacement | -0.06 | 0.96 |
| IGT Performance | 0.15 | 0.88 |
| CUD: Sex | -0.69 | 0.49 |
| CUD: Tobacco use in the last year | -2.72 | 0.01** |
| CUD: Mean Framewise Displacement | 0.19 | 0.85 |
| CUD:IGT Performance | -0.30 | 0.76 |

Subc
| Name | T-Stat | P Value |
| --- | --- | --- |
| CUD | 1.67 | 0.10 |
| Sex | 1.09 | 0.28 |
| Age | -1.73 | 0.09 |
| Education | -0.37 | 0.71 |
| Tobacco Use | -0.61 | 0.54 |
| Mean Framewise Displacement | 1.58 | 0.12 |
| IGT Performance | 1.71 | 0.09 |
| CUD: Sex | -0.82 | 0.42 |
| CUD: Tobacco use in the last year | 0.28 | 0.78 |
| CUD: Mean Framewise Displacement | -1.53 | 0.13 |
| CUD:IGT Performance | -1.10 | 0.28 |

Default
| Name | T-Stat | P Value |
| --- | --- | --- |
| CUD | 2.43 | 0.02* |
| Sex | 1.15 | 0.25 |
| Age | 1.86 | 0.07 |
| Education | -0.33 | 0.74 |
| Tobacco Use | 0.97 | 0.33 |
| Mean Framewise Displacement | 1.51 | 0.14 |
| IGT Performance | 1.38 | 0.17 |
| CUD: Sex | -1.85 | 0.07 |
| CUD: Tobacco use in the last year | -0.70 | 0.49 |
| CUD: Mean Framewise Displacement | -0.78 | 0.44 |
| CUD:IGT Performance | -1.34 | 0.18 |

Limbic
| Name | T-Stat | P Value |
| --- | --- | --- |
| CUD | 2.08 | 0.04* |
| Sex | 0.15 | 0.88 |
| Age | 1.14 | 0.26 |
| Education | 0.89 | 0.38 |
| Tobacco Use | 0.39 | 0.70 |
| Mean Framewise Displacement | 0.64 | 0.52 |
| IGT Performance | 1.32 | 0.19 |
| CUD: Sex | -0.34 | 0.73 |
| CUD: Tobacco use in the last year | -0.47 | 0.64 |
| CUD: Mean Framewise Displacement | -2.32 | 0.02* |
| CUD:IGT Performance | -1.21 | 0.23 |

SomMot
| Name | T-Stat | P Value |
| --- | --- | --- |
| CUD | -0.30 | 0.76 |
| Sex | -0.34 | 0.73 |
| Age | -0.54 | 0.59 |
| Education | -1.58 | 0.12 |
| Tobacco Use | 1.05 | 0.30 |
| Mean Framewise Displacement | 0.00 | 1.00 |
| IGT Performance | 0.10 | 0.92 |
| CUD: Sex | 0.61 | 0.54 |
| CUD: Tobacco use in the last year | -1.42 | 0.16 |
| CUD: Mean Framewise Displacement | 0.08 | 0.93 |
| CUD:IGT Performance | 0.02 | 0.98 |

| Name | T-Stat | P Value |
| --- | --- | --- |
| CUD | 1.03 | 0.31 |
| Sex | 0.78 | 0.44 |
| Age | -2.93 | 0.00 |
| Education | 0.74 | 0.46 |
| Tobacco Use | 1.49 | 0.14 |
| Mean Framewise Displacement | 0.34 | 0.73 |
| IGT Performance | 0.47 | 0.64 |
| CUD: Sex | -0.22 | 0.82 |
| CUD: Tobacco use in the last year | -1.80 | 0.08 |
| CUD: Mean Framewise Displacement | -0.68 | 0.50 |
| CUD:IGT Performance | -0.53 | 0.59 |

**Table 5.**
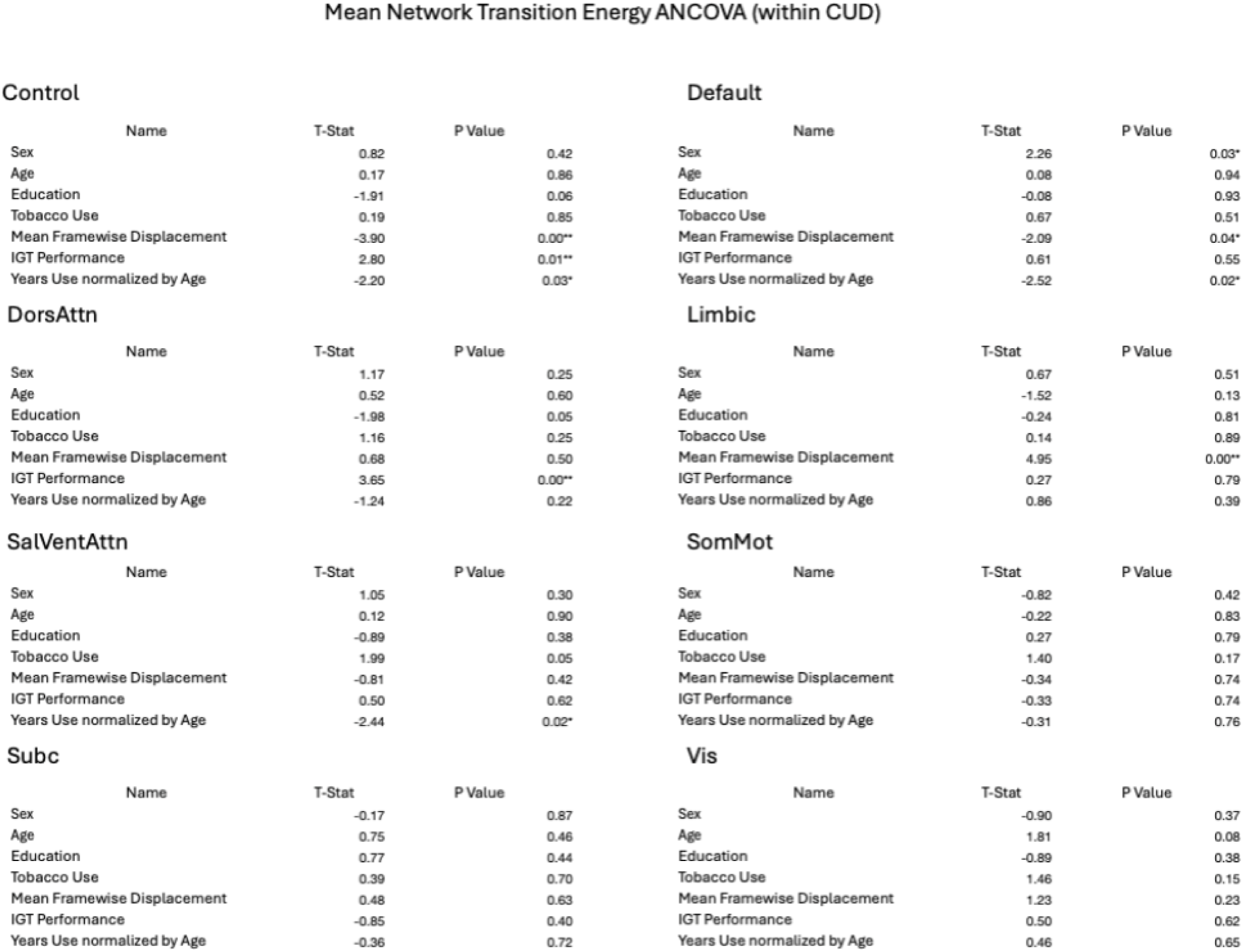
ANCOVA for understanding the impact of CUD duration on transition energies for canonical Yeo Networks.

Control
| Name | T-Stat | P Value |
| --- | --- | --- |
| Sex | 0.82 | 0.42 |
| Age | 0.17 | 0.86 |
| Education | -1.91 | 0.06 |
| Tobacco Use | 0.19 | 0.85 |
| Mean Framewise Displacement | -3.90 | 0.00** |
| IGT Performance | 2.80 | 0.01** |
| Years Use normalized by Age | -2.20 | 0.03* |

DorsAttn
| Name | T-Stat | P Value |
| --- | --- | --- |
| Sex | 1.17 | 0.25 |
| Age | 0.52 | 0.60 |
| Education | -1.98 | 0.05 |
| Tobacco Use | 1.16 | 0.25 |
| Mean Framewise Displacement | 0.68 | 0.50 |
| IGT Performance | 3.65 | 0.00** |
| Years Use normalized by Age | -1.24 | 0.22 |

SalVentAttn
| Name | T-Stat | P Value |
| --- | --- | --- |
| Sex | 1.05 | 0.30 |
| Age | 0.12 | 0.90 |
| Education | -0.89 | 0.38 |
| Tobacco Use | 1.99 | 0.05 |
| Mean Framewise Displacement | -0.81 | 0.42 |
| IGT Performance | 0.50 | 0.62 |
| Years Use normalized by Age | -2.44 | 0.02* |

Subc
| Name | T-Stat | P Value |
| --- | --- | --- |
| Sex | -0.17 | 0.87 |
| Age | 0.75 | 0.46 |
| Education | 0.77 | 0.44 |
| Tobacco Use | 0.39 | 0.70 |
| Mean Framewise Displacement | 0.48 | 0.63 |
| IGT Performance | -0.85 | 0.40 |
| Years Use normalized by Age | -0.36 | 0.72 |

Default
| Name | T-Stat | P Value |
| --- | --- | --- |
| Sex | 2.26 | 0.03* |
| Age | 0.08 | 0.94 |
| Education | -0.08 | 0.93 |
| Tobacco Use | 0.67 | 0.51 |
| Mean Framewise Displacement | -2.09 | 0.04* |
| IGT Performance | 0.61 | 0.55 |
| Years Use normalized by Age | -2.52 | 0.02* |

Limbic
| Name | T-Stat | P Value |
| --- | --- | --- |
| Sex | 0.67 | 0.51 |
| Age | -1.52 | 0.13 |
| Education | -0.24 | 0.81 |
| Tobacco Use | 0.14 | 0.89 |
| Mean Framewise Displacement | 4.95 | 0.00** |
| IGT Performance | 0.27 | 0.79 |
| Years Use normalized by Age | 0.86 | 0.39 |

SomMot
| Name | T-Stat | P Value |
| --- | --- | --- |
| Sex | -0.82 | 0.42 |
| Age | -0.22 | 0.83 |
| Education | 0.27 | 0.79 |
| Tobacco Use | 1.40 | 0.17 |
| Mean Framewise Displacement | -0.34 | 0.74 |
| IGT Performance | -0.33 | 0.74 |
| Years Use normalized by Age | -0.31 | 0.76 |

| Name | T-Stat | P Value |
| --- | --- | --- |
| Sex | -0.90 | 0.37 |
| Age | 1.81 | 0.08 |
| Education | -0.89 | 0.38 |
| Tobacco Use | 1.46 | 0.15 |
| Mean Framewise Displacement | 1.23 | 0.23 |
| IGT Performance | 0.50 | 0.62 |
| Years Use normalized by Age | 0.46 | 0.65 |

**Figure S8.**
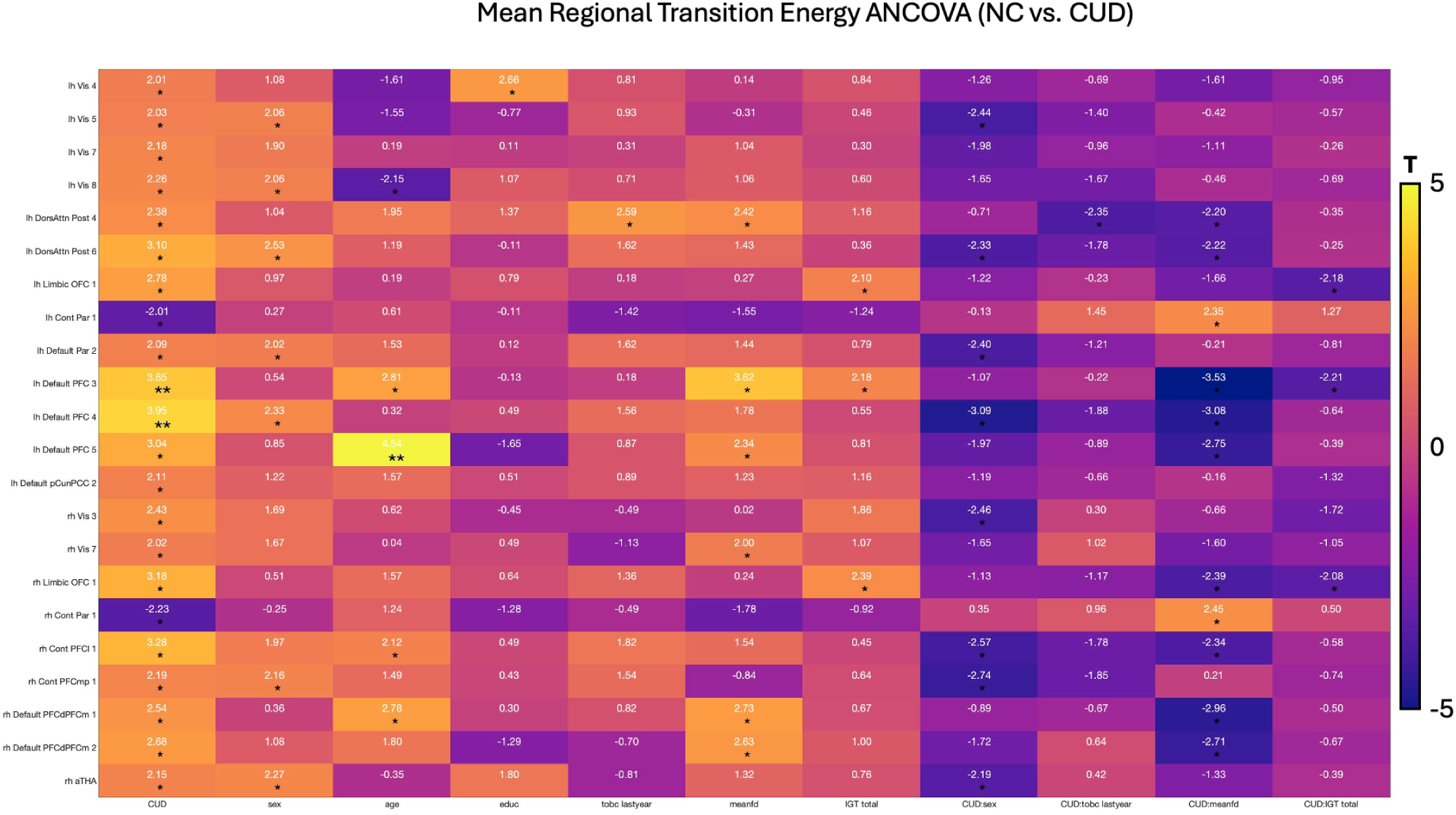
ANCOVA for comparing transition energies between regions for groups indicated that the energetic landscape was lower for people with CUD in higher order networks, with these effects being driven by regions in frontal half of the brain belonging to the default network.

**Figure S9.**
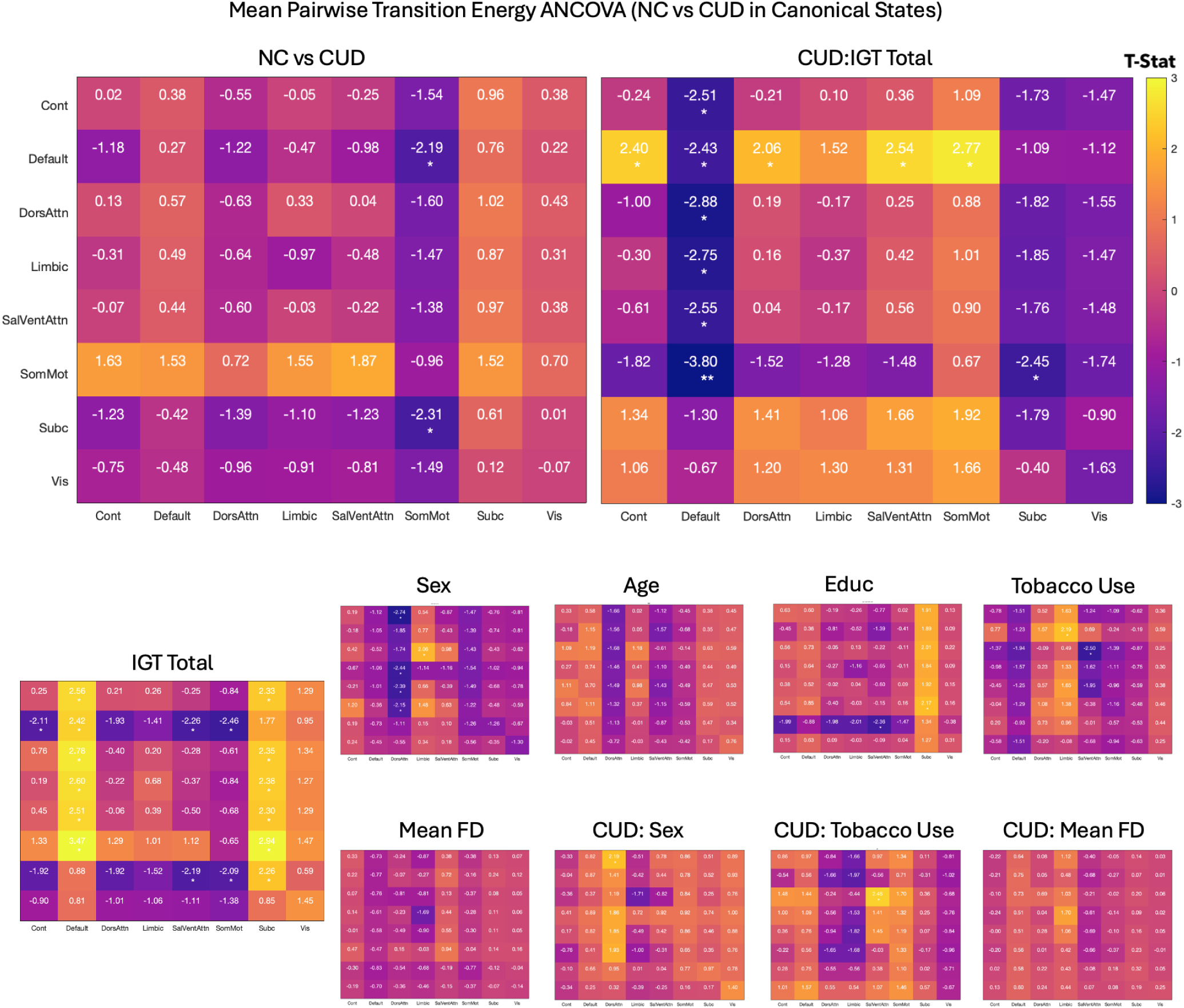
ANCOVAs for canonical state analysis underscored the impact of CUD on risk taking behaviors, with this impact altering energy dynamics from the SomMot to the default network

**Figure S10.**
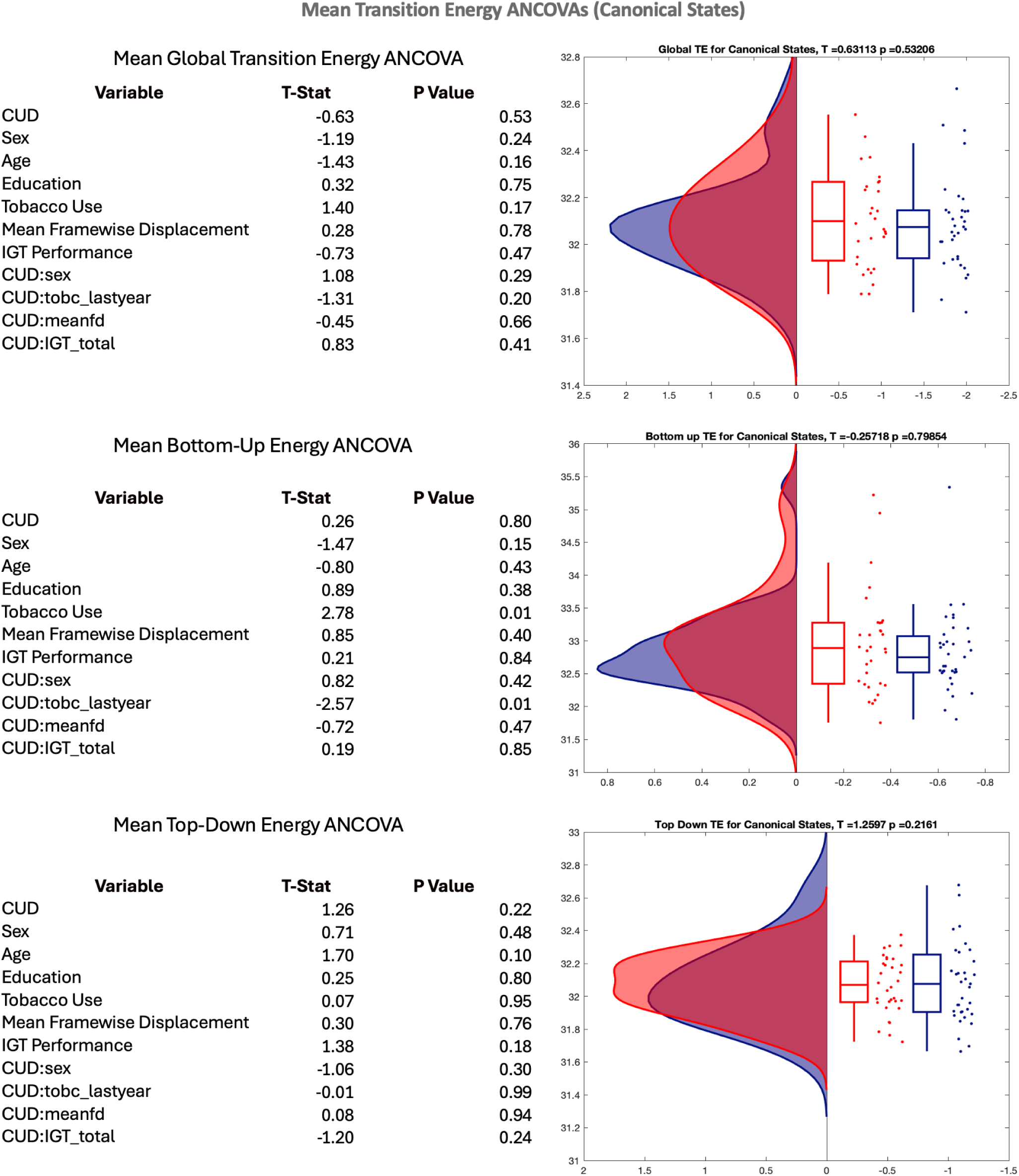
ANCOVAs for global transition, bottom up and top down energy metrics obtained from a canonical state analysis.

**Figure S11.**
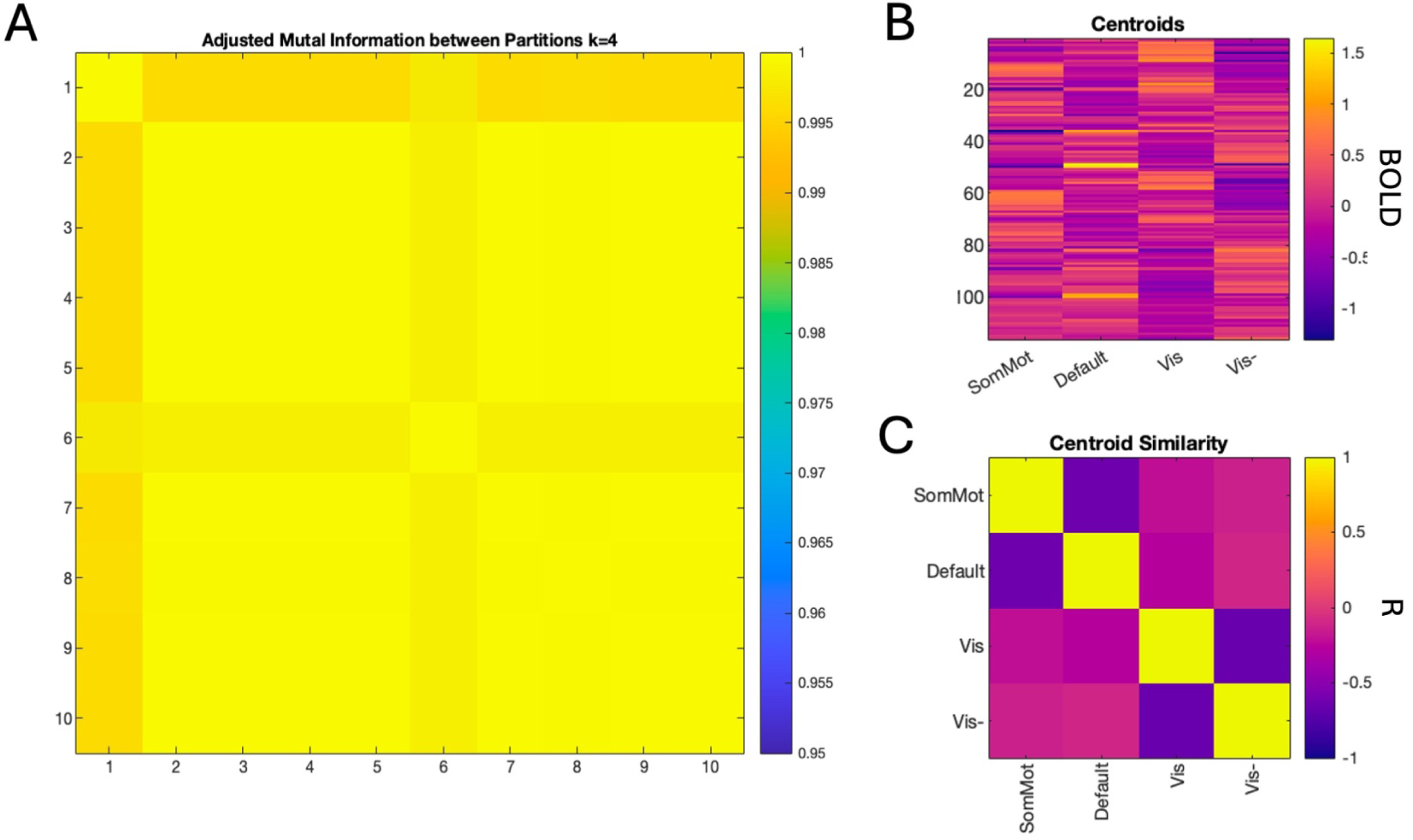
We replicated the main analysis with k = 4. We found that global energy dynamics were lower in people with CUD, and the same regions drove the difference for dynamics between people with CUD and NC. We did not find differences in temporal dynamics that survived correction. Results for the impact of CUD on risk taking behaviors and the cumulative impact of using cocaine across one’s lifetime remained similar across the different k’s chosen.

**Figure S12.**
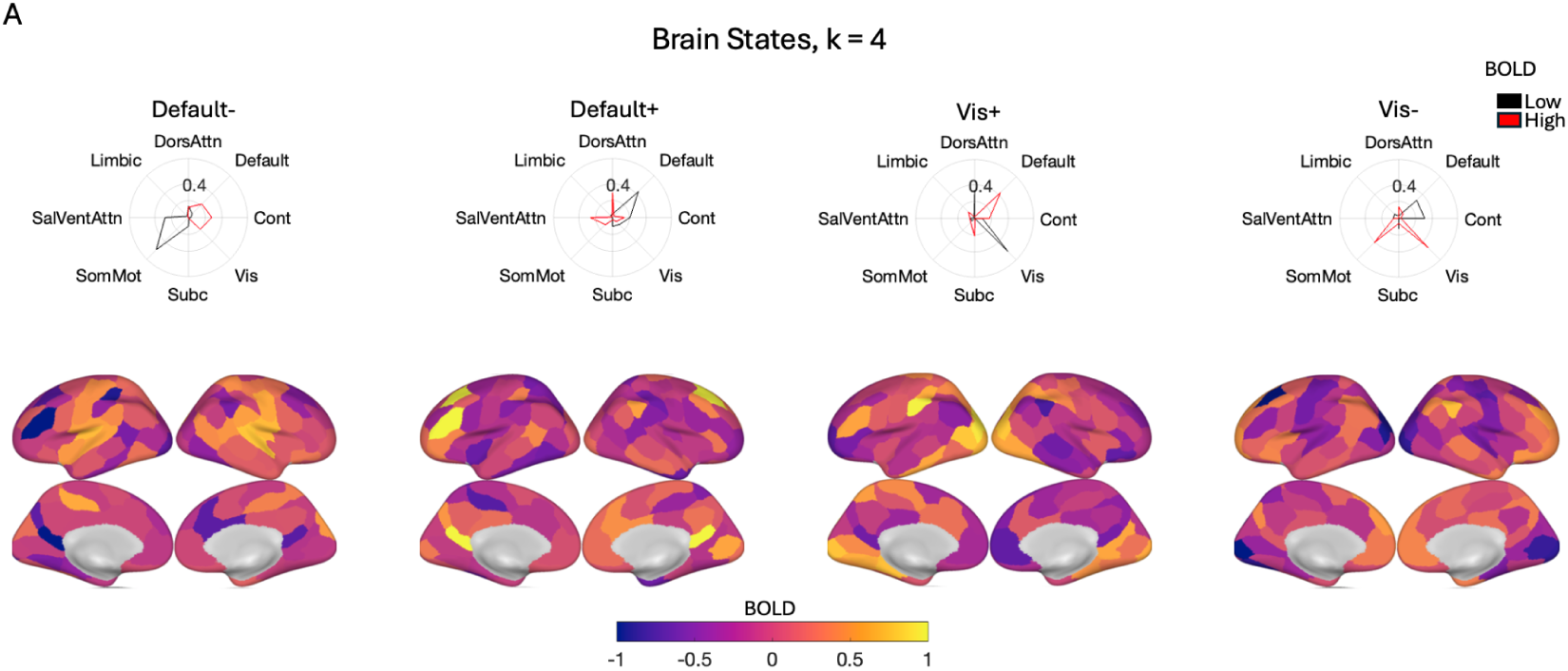
K means clustering revealed four brain states consisting of activation and deactivation of the somatomotor/default and visual networks. Temporal dynamics were analyzed, and it was found that people with cocaine use disorder had significantly fewer appearances of the default state.

**Figure S13.**
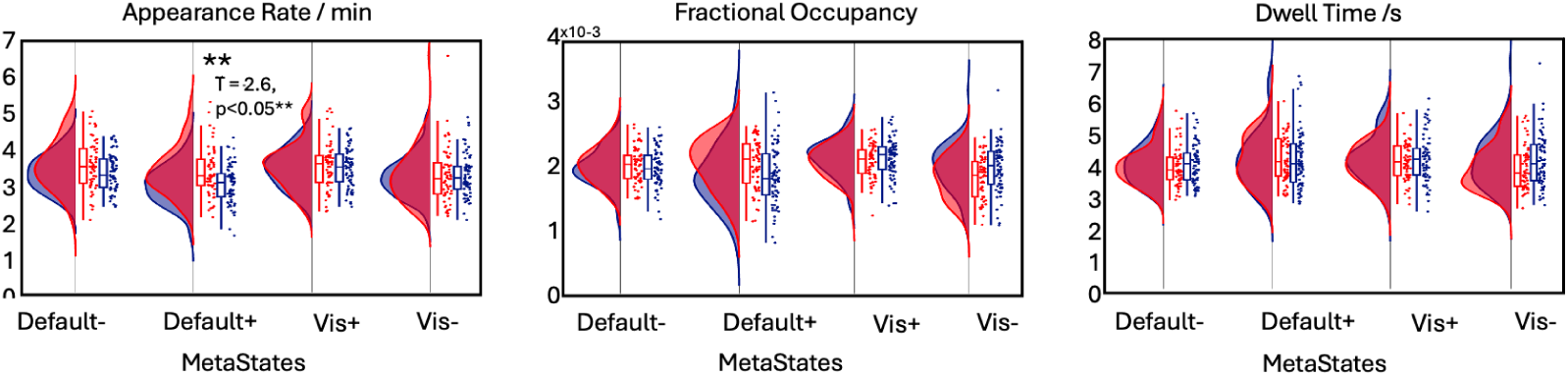
Temporal dynamics were analyzed, and it was found that people with cocaine use disorder had significantly fewer appearances of the default state.

**Figure S14.**
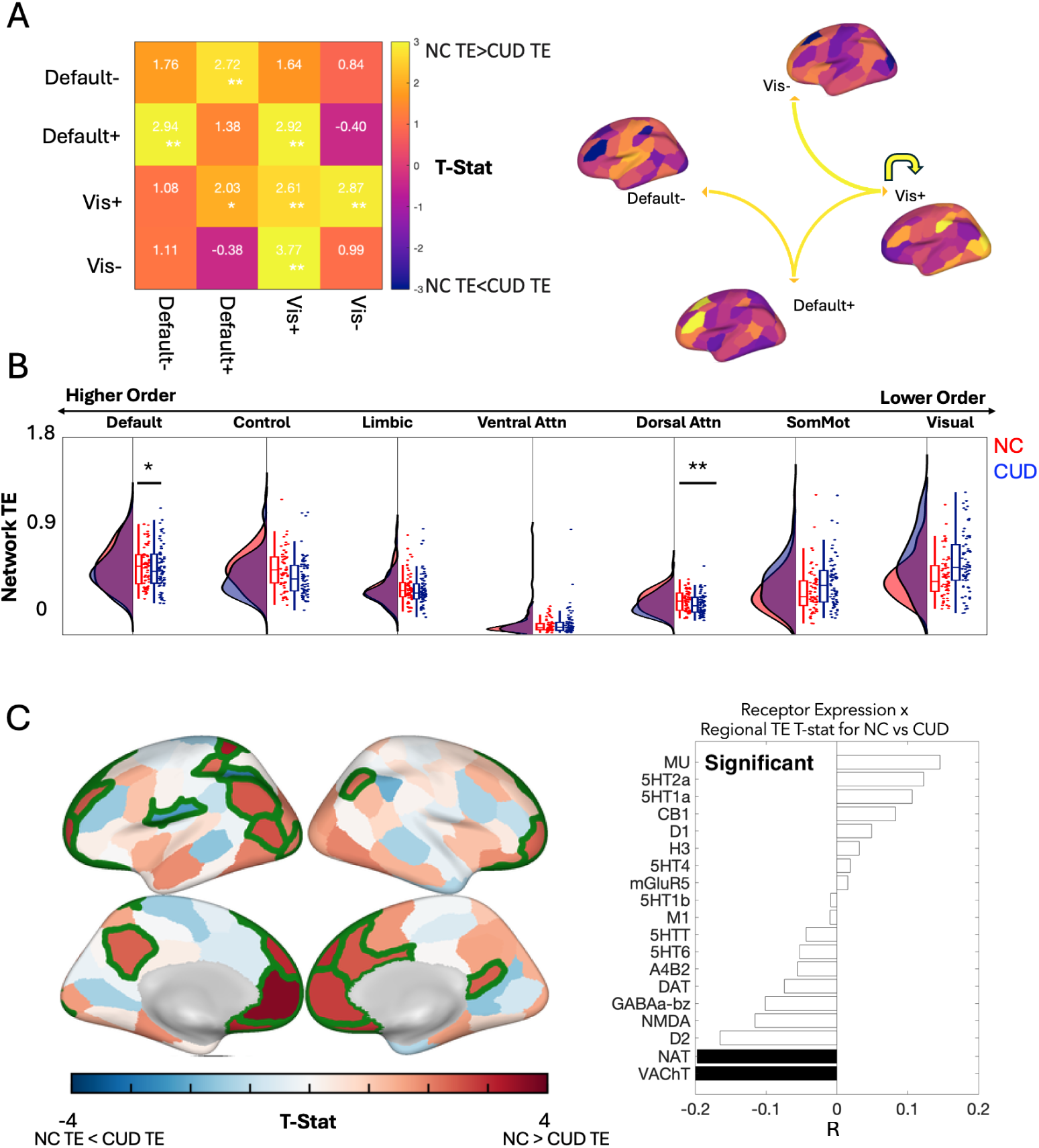
Recurrent States of Brain Activity underscore how Energy Dynamics differ for People with Cocaine Use Disorder (CUD) and Non-User Controls (NC). (**A**) Using an ANCOVA, we identified differences in energy dynamics for four brain states for people with CUD and NC, with significant differences that survive correction depicted in color in the spider-plot. (**B**) Average network transition energies showed a trend with transition energies being lower for people with CUD in the default mode network (pFDR <0.05, before correction) and dorsal attention networks (pFDR <0.05, after correction) in comparison to NC. (**C**) ANCOVAs for average regional TEs showed multiple regions belonging to higher order networks which had significantly lower transition energies in people with CUD. p-values were corrected for multiple comparisons (Benjamini-Hochberg) where indicated. **corrected p < 0.05. Statistical comparisons were made using ANCOVAs with covariates including age, sex, mean framewise displacement, tobacco use, performance on the IGT task and interaction terms for drug use with age, sex, tobacco use and performance on the IGT task. p-values were corrected for multiple comparisons (Benjamini-Hochberg) where indicated. **corrected p < 0.05.

**Figure S15.**
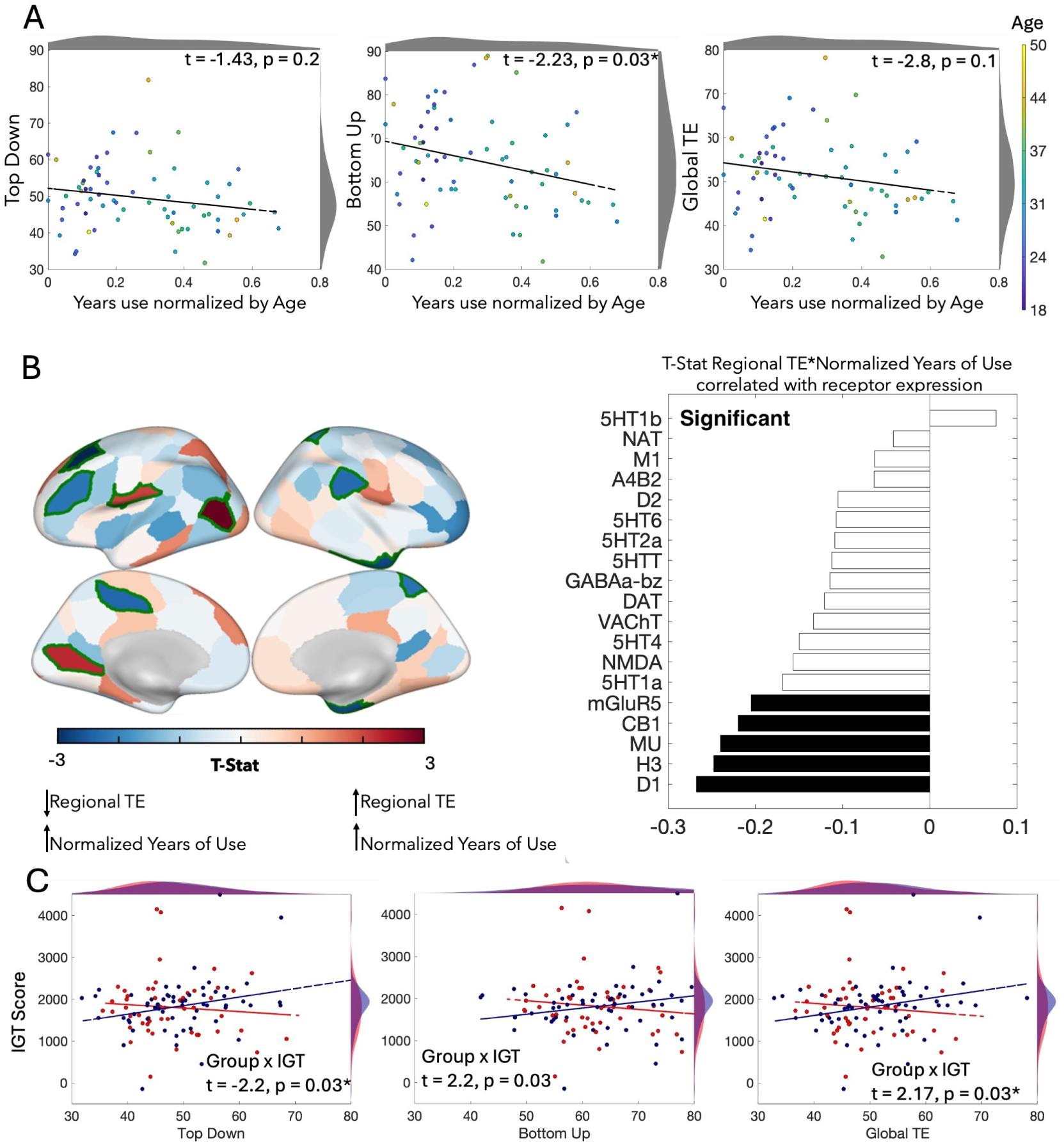
Global energy dynamics decrease across the years of cocaine use, and involve specific receptor families. (**A**) Global energy dynamics, categorized here by global transition energy, bottom up energy and top downn energy, trended towards declining over the course of CUD, with global transition energy and top-down energy being significant after correction. (**B**) Default and control network (highlighted in green) and regional transition energies correlated significantly with the numbers of years used normalized by age. We correlated the t-statistic for years of use normalized by age with receptor concentration for ANCOVAs that were ran for just people with CUD, and found that the decrease in transition energy over the years correlated positively with the receptor expression of excitatory receptors that are implicated in the glutamatergic toxicity that is central to coaine use disorder. p-values were corrected for multiple comparisons (Benjamini-Hochberg) where indicated. **corrected p < 0.05.

**Figure S16.**
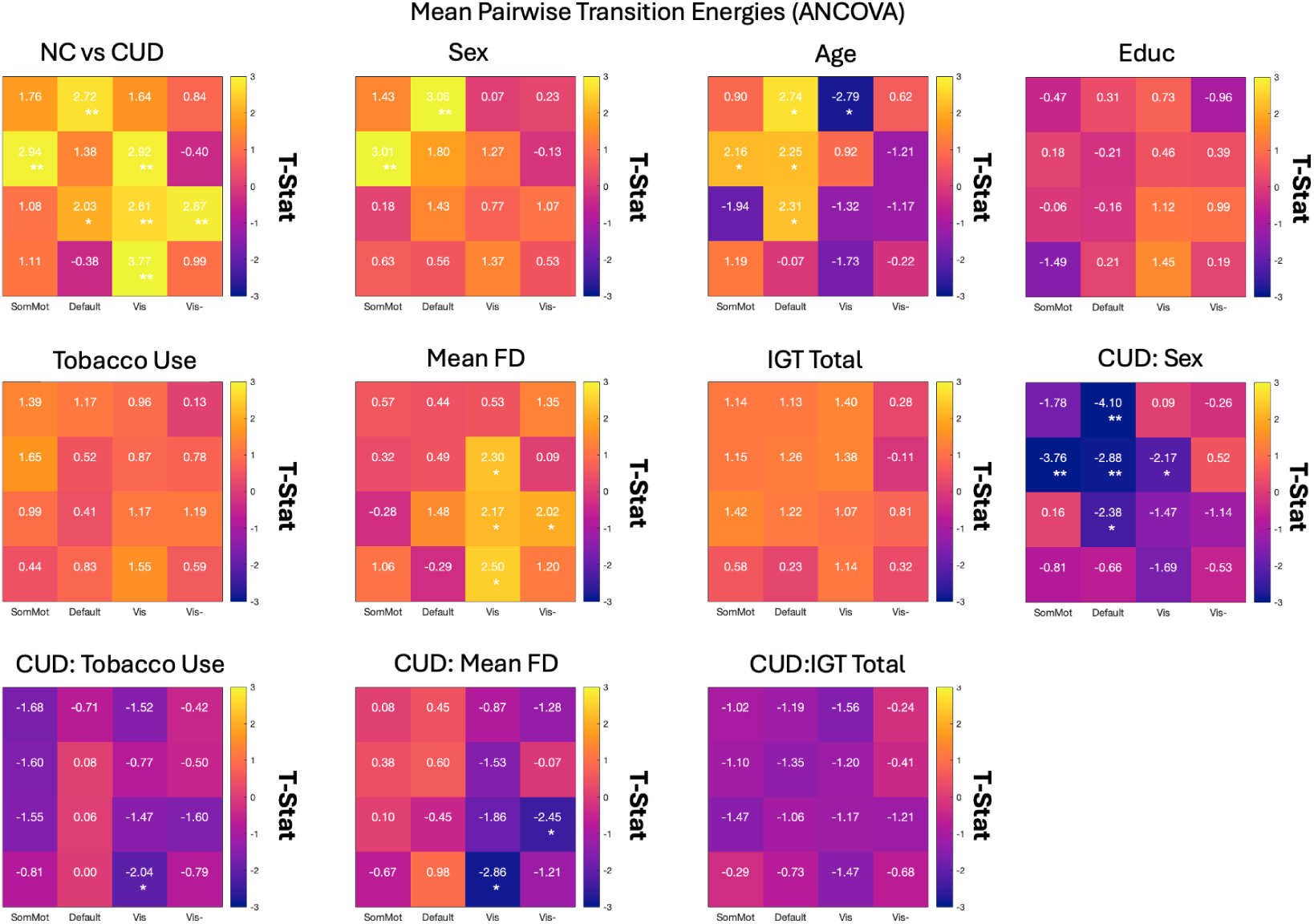
ANCOVA result for transition energies on a pairwise level between CUD and NC

**Figure S17.**
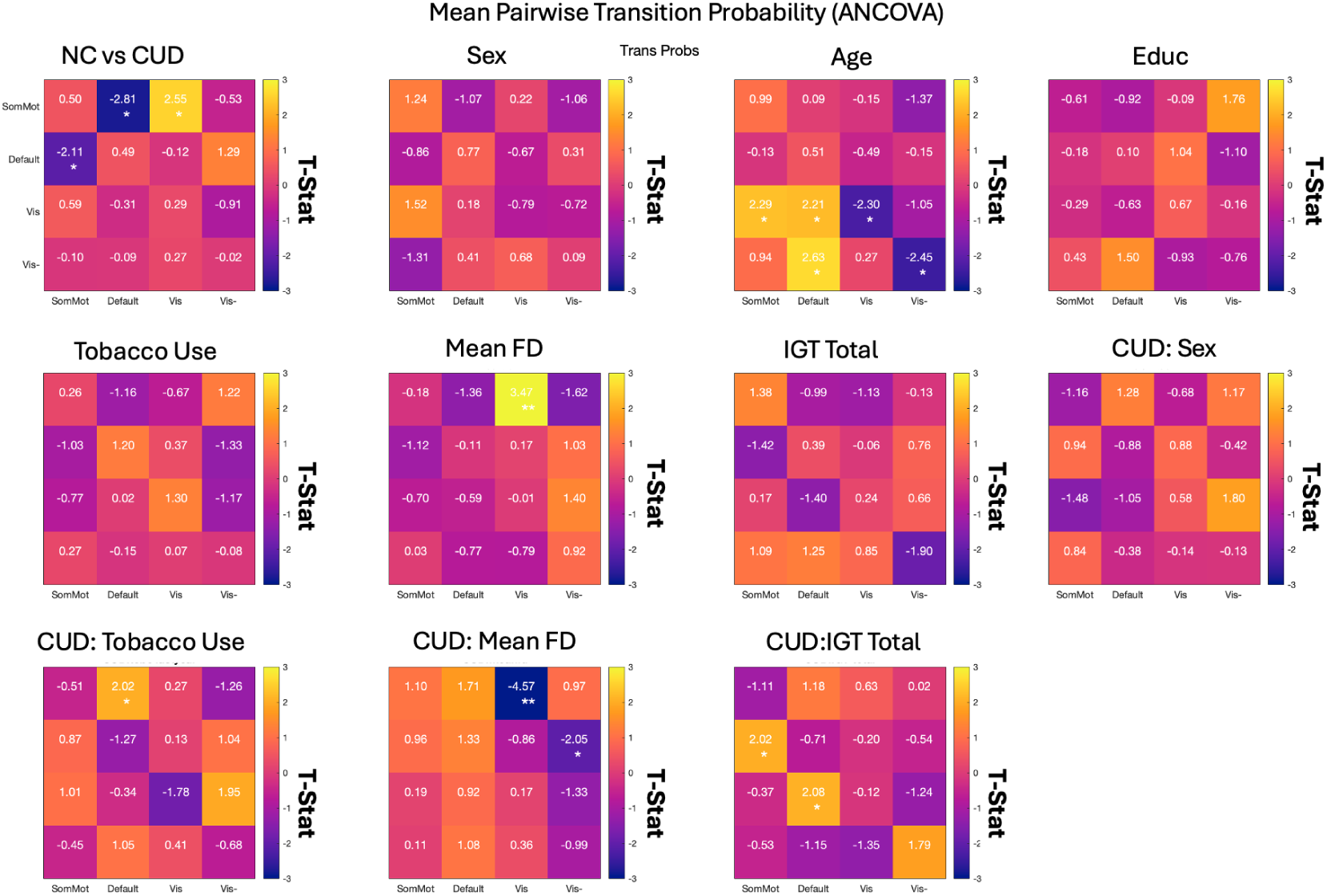
ANCOVA result for transition probabilities on a pairwise level between CUD and NC

**Table 6.**
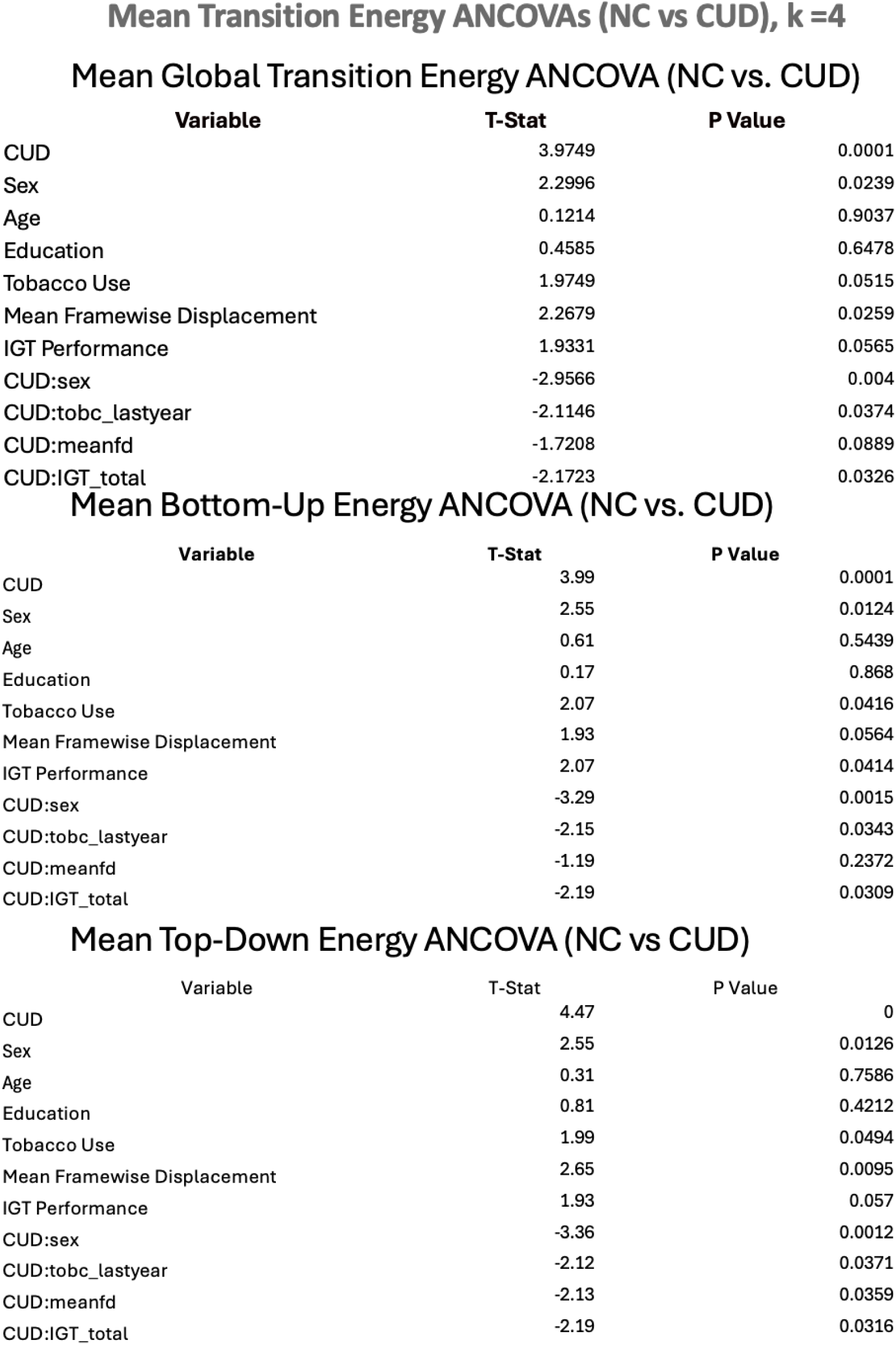
K =4, ANCOVA for comparing transition energies between groups on a global level, as well as to understand the difference between their top down and bottom up balance of signaling.

Mean Global Transition Energy ANCOVA (NC vs. CUD)
| Variable | T-Stat | P Value |
| --- | --- | --- |
| CUD | 3.9749 | 0.0001 |
| Sex | 2.2996 | 0.0239 |
| Age | 0.1214 | 0.9037 |
| Education | 0.4585 | 0.6478 |
| Tobacco Use | 1.9749 | 0.0515 |
| Mean Framewise Displacement | 2.2679 | 0.0259 |
| IGT Performance | 1.9331 | 0.0565 |
| CUD:sex | -2.9566 | 0.004 |
| CUD:tobc_lastyear | -2.1146 | 0.0374 |
| CUD:meanfd | -1.7208 | 0.0889 |
| CUD:IGT_total | -2.1723 | 0.0326 |

Mean Bottom-Up Energy ANCOVA (NC vs. CUD)
| Variable | T-Stat | P Value |
| --- | --- | --- |
| CUD | 3.99 | 0.0001 |
| Sex | 2.55 | 0.0124 |
| Age | 0.61 | 0.5439 |
| Education | 0.17 | 0.868 |
| Tobacco Use | 2.07 | 0.0416 |
| Mean Framewise Displacement | 1.93 | 0.0564 |
| IGT Performance | 2.07 | 0.0414 |
| CUD:sex | -3.29 | 0.0015 |
| CUD:tobc_lastyear | -2.15 | 0.0343 |
| CUD:meanfd | -1.19 | 0.2372 |
| CUD:IGT_total | -2.19 | 0.0309 |

| Variable | T-Stat | P Value |
| --- | --- | --- |
| CUD | 4.47 | 0 |
| Sex | 2.55 | 0.0126 |
| Age | 0.31 | 0.7586 |
| Education | 0.81 | 0.4212 |
| Tobacco Use | 1.99 | 0.0494 |
| Mean Framewise Displacement | 2.65 | 0.0095 |
| IGT Performance | 1.93 | 0.057 |
| CUD:sex | -3.36 | 0.0012 |
| CUD:tobc_lastyear | -2.12 | 0.0371 |
| CUD:meanfd | -2.13 | 0.0359 |
| CUD:IGT_total | -2.19 | 0.0316 |

**Table 7.**
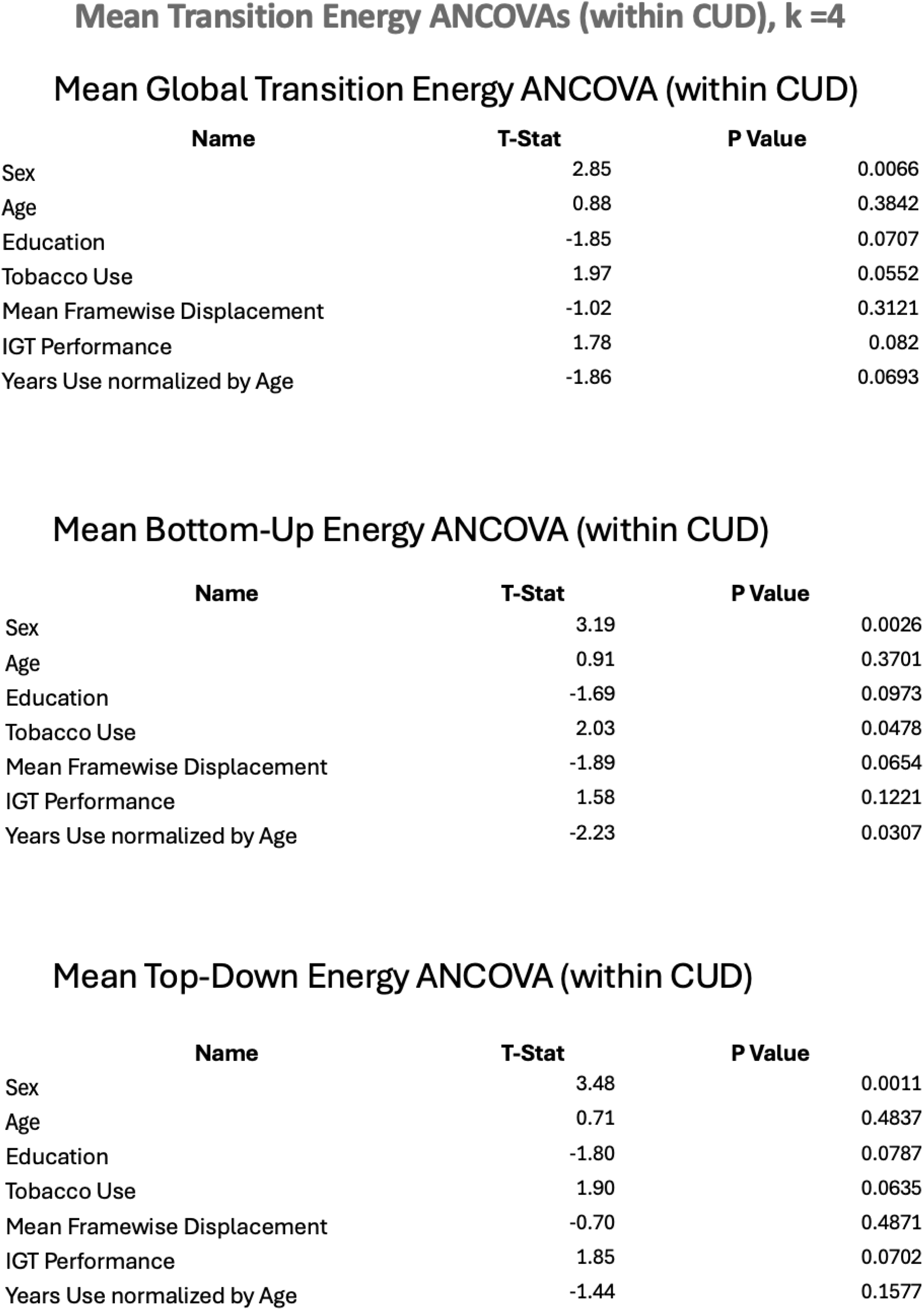
K =4, ANCOVA for understanding the impact of cocaine use over the years normalized by age on a global level, as well as on top down and bottom up signaling.

Mean Global Transition Energy ANCOVA (within CUD)
|  | Name | T-Stat | P Value |
| --- | --- | --- | --- |
| Sex |  | 2.85 | 0.0066 |
| Age |  | 0.88 | 0.3842 |
| Education |  | -1.85 | 0.0707 |
| Tobacco Use |  | 1.97 | 0.0552 |
| Mean Framewise Displacement |  | -1.02 | 0.3121 |
| IGT Performance |  | 1.78 | 0.082 |
| Years Use normalized by Age |  | -1.86 | 0.0693 |

Mean Bottom-Up Energy ANCOVA (within CUD)
|  | Name | T-Stat | P Value |
| --- | --- | --- | --- |
| Sex |  | 3.19 | 0.0026 |
| Age |  | 0.91 | 0.3701 |
| Education |  | -1.69 | 0.0973 |
| Tobacco Use |  | 2.03 | 0.0478 |
| Mean Framewise Displacement |  | -1.89 | 0.0654 |
| IGT Performance |  | 1.58 | 0.1221 |
| Years Use normalized by Age |  | -2.23 | 0.0307 |

|  | Name | T-Stat | P Value |
| --- | --- | --- | --- |
| Sex |  | 3.48 | 0.0011 |
| Age |  | 0.71 | 0.4837 |
| Education |  | -1.80 | 0.0787 |
| Tobacco Use |  | 1.90 | 0.0635 |
| Mean Framewise Displacement |  | -0.70 | 0.4871 |
| IGT Performance |  | 1.85 | 0.0702 |
| Years Use normalized by Age |  | -1.44 | 0.1577 |

**Table 8.**
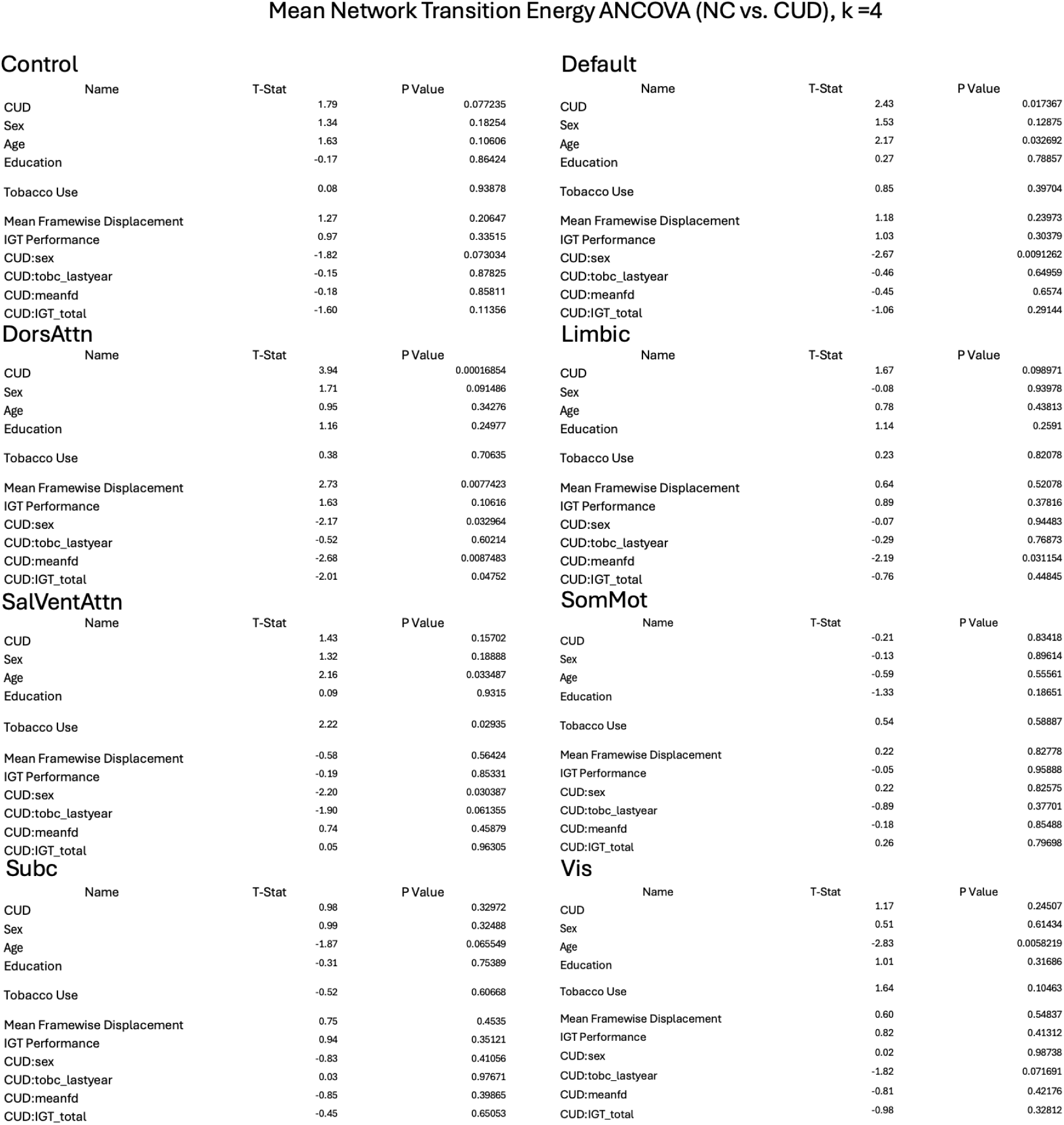
K = 4, ANCOVA for comparing transition energies between groups for Yeo Networks.

Control
| Name | T-Stat | P Value |
| --- | --- | --- |
| CUD | 1.79 | 0.077235 |
| Sex | 1.34 | 0.18254 |
| Age | 1.63 | 0.10606 |
| Education | -0.17 | 0.86424 |
| Tobacco Use | 0.08 | 0.93878 |
| Mean Framewise Displacement | 1.27 | 0.20647 |
| IGT Performance | 0.97 | 0.33515 |
| CUD:sex | -1.82 | 0.073034 |
| CUD:tobc_lastyear | -0.15 | 0.87825 |
| CUD:meanfd | -0.18 | 0.85811 |
| CUD:IGT_total | -1.60 | 0.11356 |

DorsAttn
| Name | T-Stat | P Value |
| --- | --- | --- |
| CUD | 3.94 | 0.00016854 |
| Sex | 1.71 | 0.091486 |
| Age | 0.95 | 0.34276 |
| Education | 1.16 | 0.24977 |
| Tobacco Use | 0.38 | 0.70635 |
| Mean Framewise Displacement | 2.73 | 0.0077423 |
| IGT Performance | 1.63 | 0.10616 |
| CUD:sex | -2.17 | 0.032964 |
| CUD:tobc_lastyear | -0.52 | 0.60214 |
| CUD:meanfd | -2.68 | 0.0087483 |
| CUD:IGT_total | -2.01 | 0.04752 |

SalVentAttn
| Name | T-Stat | P Value |
| --- | --- | --- |
| CUD | 1.43 | 0.15702 |
| Sex | 1.32 | 0.18888 |
| Age | 2.16 | 0.033487 |
| Education | 0.09 | 0.9315 |
| Tobacco Use | 2.22 | 0.02935 |
| Mean Framewise Displacement | -0.58 | 0.56424 |
| IGT Performance | -0.19 | 0.85331 |
| CUD:sex | -2.20 | 0.030387 |
| CUD:tobc_lastyear | -1.90 | 0.061355 |
| CUD:meanfd | 0.74 | 0.45879 |
| CUD:IGT_total | 0.05 | 0.96305 |

Subc
| Name | T-Stat | P Value |
| --- | --- | --- |
| CUD | 0.98 | 0.32972 |
| Sex | 0.99 | 0.32488 |
| Age | -1.87 | 0.065549 |
| Education | -0.31 | 0.75389 |
| Tobacco Use | -0.52 | 0.60668 |
| Mean Framewise Displacement | 0.75 | 0.4535 |
| IGT Performance | 0.94 | 0.35121 |
| CUD:sex | -0.83 | 0.41056 |
| CUD:tobc_lastyear | 0.03 | 0.97671 |
| CUD:meanfd | -0.85 | 0.39865 |
| CUD:IGT_total | -0.45 | 0.65053 |

Default
| Name | T-Stat | P Value |
| --- | --- | --- |
| CUD | 2.43 | 0.017367 |
| Sex | 1.53 | 0.12875 |
| Age | 2.17 | 0.032692 |
| Education | 0.27 | 0.78857 |
| Tobacco Use | 0.85 | 0.39704 |
| Mean Framewise Displacement | 1.18 | 0.23973 |
| IGT Performance | 1.03 | 0.30379 |
| CUD:sex | -2.67 | 0.0091262 |
| CUD:tobc_lastyear | -0.46 | 0.64959 |
| CUD:meanfd | -0.45 | 0.6574 |
| CUD:IGT_total | -1.06 | 0.29144 |

Limbic
| Name | T-Stat | P Value |
| --- | --- | --- |
| CUD | 1.67 | 0.098971 |
| Sex | -0.08 | 0.93978 |
| Age | 0.78 | 0.43813 |
| Education | 1.14 | 0.2591 |
| Tobacco Use | 0.23 | 0.82078 |
| Mean Framewise Displacement | 0.64 | 0.52078 |
| IGT Performance | 0.89 | 0.37816 |
| CUD:sex | -0.07 | 0.94483 |
| CUD:tobc_lastyear | -0.29 | 0.76873 |
| CUD:meanfd | -2.19 | 0.031154 |
| CUD:IGT_total | -0.76 | 0.44845 |

SomMot
| Name | T-Stat | P Value |
| --- | --- | --- |
| CUD | -0.21 | 0.83418 |
| Sex | -0.13 | 0.89614 |
| Age | -0.59 | 0.55561 |
| Education | -1.33 | 0.18651 |
| Tobacco Use | 0.54 | 0.58887 |
| Mean Framewise Displacement | 0.22 | 0.82778 |
| IGT Performance | -0.05 | 0.95888 |
| CUD:sex | 0.22 | 0.82575 |
| CUD:tobc_lastyear | -0.89 | 0.37701 |
| CUD:meanfd | -0.18 | 0.85488 |
| CUD:IGT_total | 0.26 | 0.79698 |

| Name | T-Stat | P Value |
| --- | --- | --- |
| CUD | 1.17 | 0.24507 |
| Sex | 0.51 | 0.61434 |
| Age | -2.83 | 0.0058219 |
| Education | 1.01 | 0.31686 |
| Tobacco Use | 1.64 | 0.10463 |
| Mean Framewise Displacement | 0.60 | 0.54837 |
| IGT Performance | 0.82 | 0.41312 |
| CUD:sex | 0.02 | 0.98738 |
| CUD:tobc_lastyear | -1.82 | 0.071691 |
| CUD:meanfd | -0.81 | 0.42176 |
| CUD:IGT_total | -0.98 | 0.32812 |

**Table 9.**
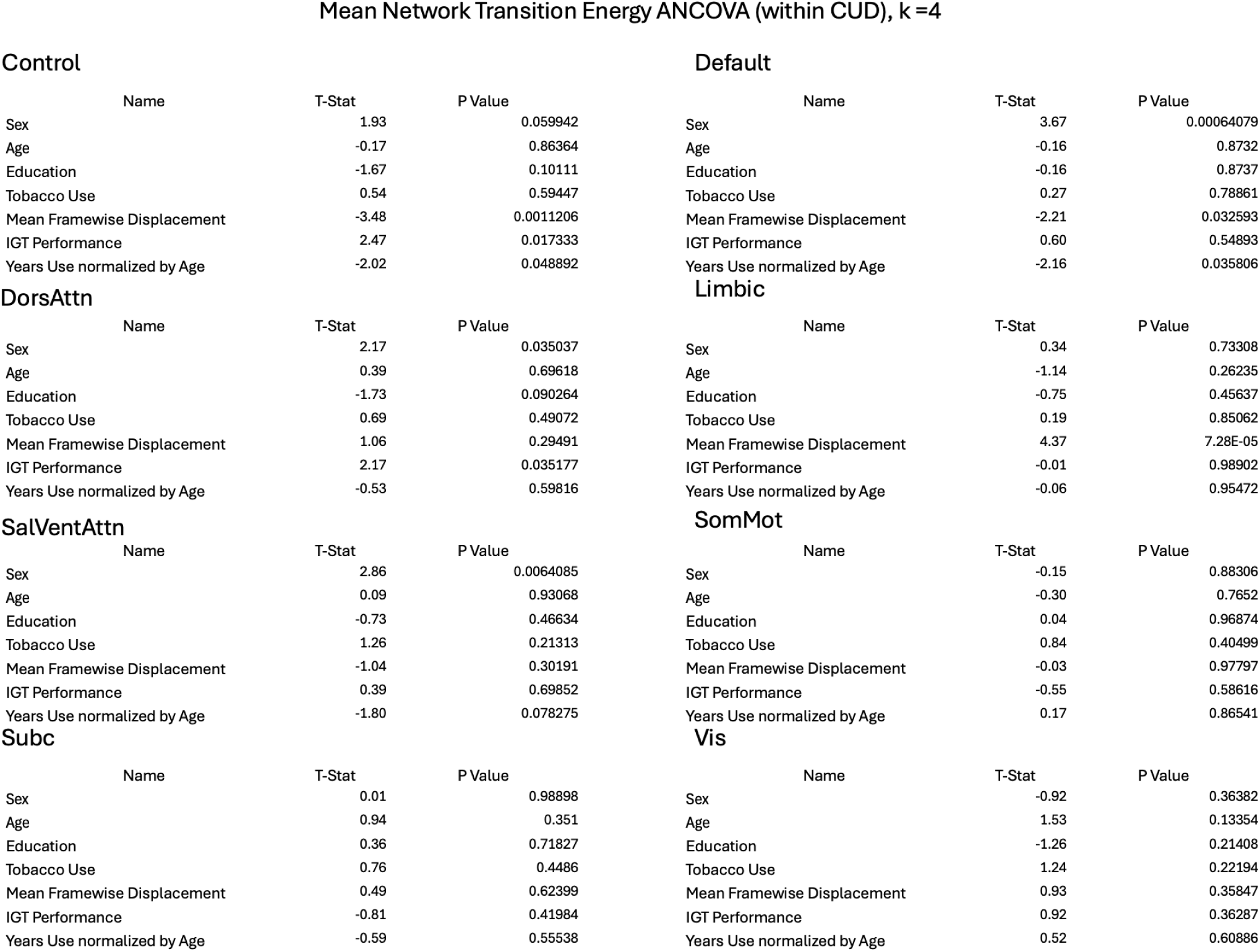
K = 4, ANCOVA for understanding the impact of cocaine use over the years normalized by age on transition energies for Yeo Networks.

Control
| Name | T-Stat | P Value |
| --- | --- | --- |
| Sex | 1.93 | 0.059942 |
| Age | -0.17 | 0.86364 |
| Education | -1.67 | 0.10111 |
| Tobacco Use | 0.54 | 0.59447 |
| Mean Framewise Displacement | -3.48 | 0.0011206 |
| IGT Performance | 2.47 | 0.017333 |
| Years Use normalized by Age | -2.02 | 0.048892 |

DorsAttn
| Name | T-Stat | P Value |
| --- | --- | --- |
| Sex | 2.17 | 0.035037 |
| Age | 0.39 | 0.69618 |
| Education | -1.73 | 0.090264 |
| Tobacco Use | 0.69 | 0.49072 |
| Mean Framewise Displacement | 1.06 | 0.29491 |
| IGT Performance | 2.17 | 0.035177 |
| Years Use normalized by Age | -0.53 | 0.59816 |

SalVentAttn
| Name | T-Stat | P Value |
| --- | --- | --- |
| Sex | 2.86 | 0.0064085 |
| Age | 0.09 | 0.93068 |
| Education | -0.73 | 0.46634 |
| Tobacco Use | 1.26 | 0.21313 |
| Mean Framewise Displacement | -1.04 | 0.30191 |
| IGT Performance | 0.39 | 0.69852 |
| Years Use normalized by Age | -1.80 | 0.078275 |

Subc
| Name | T-Stat | P Value |
| --- | --- | --- |
| Sex | 0.01 | 0.98898 |
| Age | 0.94 | 0.351 |
| Education | 0.36 | 0.71827 |
| Tobacco Use | 0.76 | 0.4486 |
| Mean Framewise Displacement | 0.49 | 0.62399 |
| IGT Performance | -0.81 | 0.41984 |
| Years Use normalized by Age | -0.59 | 0.55538 |

Default
| Name | T-Stat | P Value |
| --- | --- | --- |
| Sex | 3.67 | 0.00064079 |
| Age | -0.16 | 0.8732 |
| Education | -0.16 | 0.8737 |
| Tobacco Use | 0.27 | 0.78861 |
| Mean Framewise Displacement | -2.21 | 0.032593 |
| IGT Performance | 0.60 | 0.54893 |
| Years Use normalized by Age | -2.16 | 0.035806 |

Limbic
| Name | T-Stat | P Value |
| --- | --- | --- |
| Sex | 0.34 | 0.73308 |
| Age | -1.14 | 0.26235 |
| Education | -0.75 | 0.45637 |
| Tobacco Use | 0.19 | 0.85062 |
| Mean Framewise Displacement | 4.37 | 7.28E-05 |
| IGT Performance | -0.01 | 0.98902 |
| Years Use normalized by Age | -0.06 | 0.95472 |

SomMot
| Name | T-Stat | P Value |
| --- | --- | --- |
| Sex | -0.15 | 0.88306 |
| Age | -0.30 | 0.7652 |
| Education | 0.04 | 0.96874 |
| Tobacco Use | 0.84 | 0.40499 |
| Mean Framewise Displacement | -0.03 | 0.97797 |
| IGT Performance | -0.55 | 0.58616 |
| Years Use normalized by Age | 0.17 | 0.86541 |

| Name | T-Stat | P Value |
| --- | --- | --- |
| Sex | -0.92 | 0.36382 |
| Age | 1.53 | 0.13354 |
| Education | -1.26 | 0.21408 |
| Tobacco Use | 1.24 | 0.22194 |
| Mean Framewise Displacement | 0.93 | 0.35847 |
| IGT Performance | 0.92 | 0.36287 |
| Years Use normalized by Age | 0.52 | 0.60886 |

**Figure S18.**
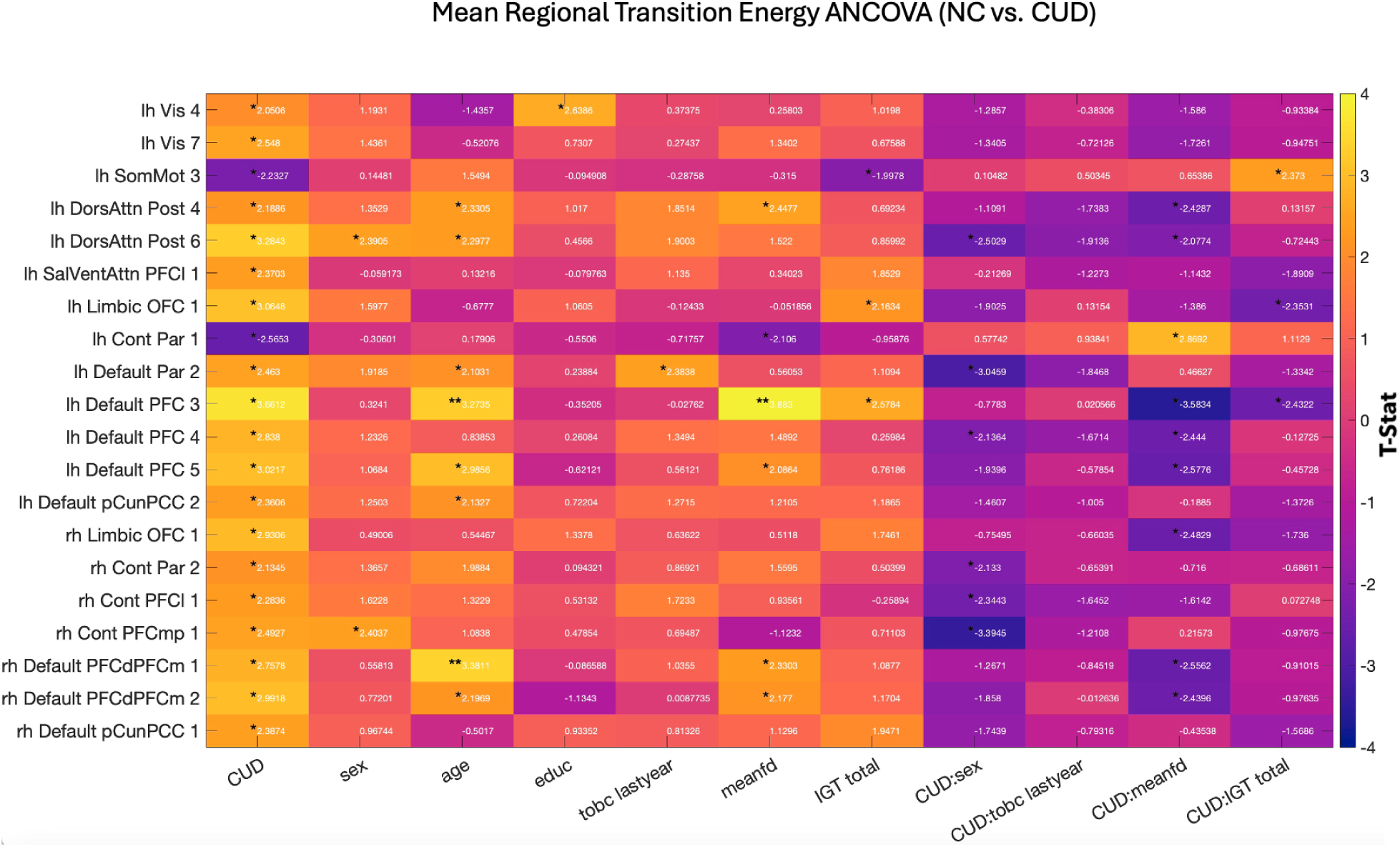
K=4, ANCOVA for comparing transition energies between regions for groups indicated that the energetic landscape was lower for people with CUD in higher order networks, with these effects being driven by regions in frontal half of the brain belonging to the default network.

## Notes

### Competing Interest Statement

The authors have declared no competing interest.

