## Supplementary figures and images for "Cocaine use disorder is associated with lower brain state transition energy particularly in higher order and excitatory networks"

### Supplemental Image 1

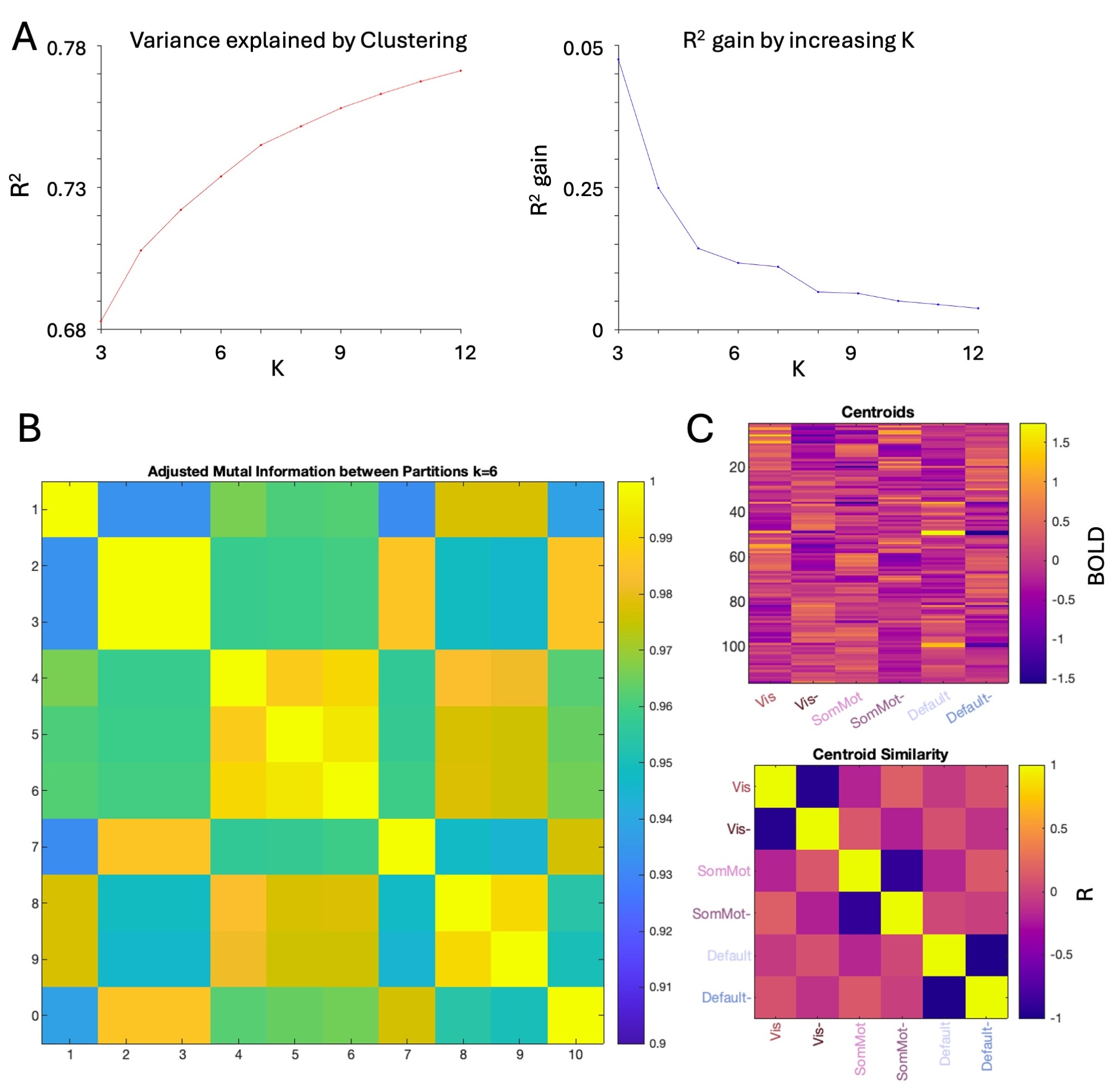

### Supplemental Image 2

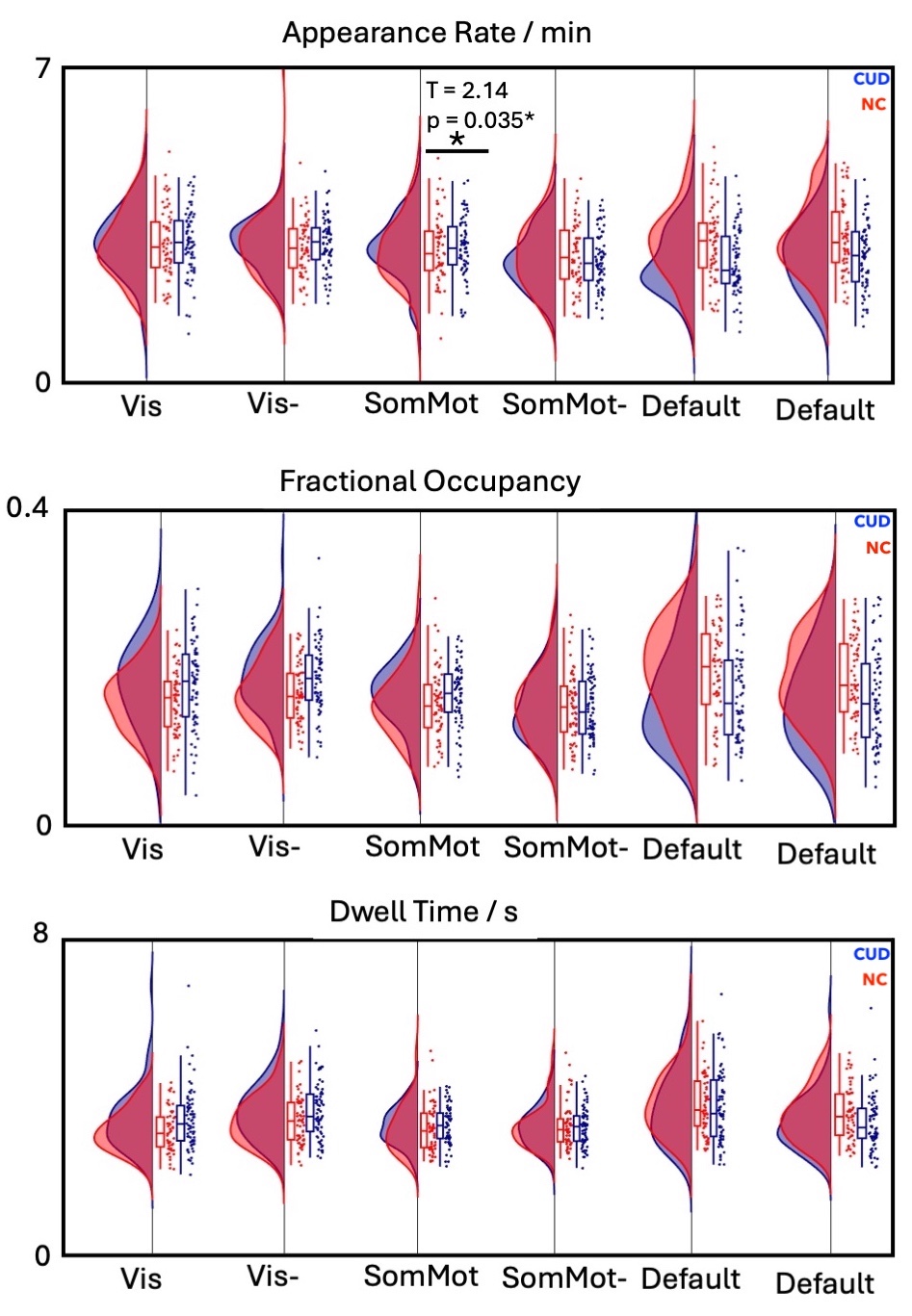

### Supplemental Image 3

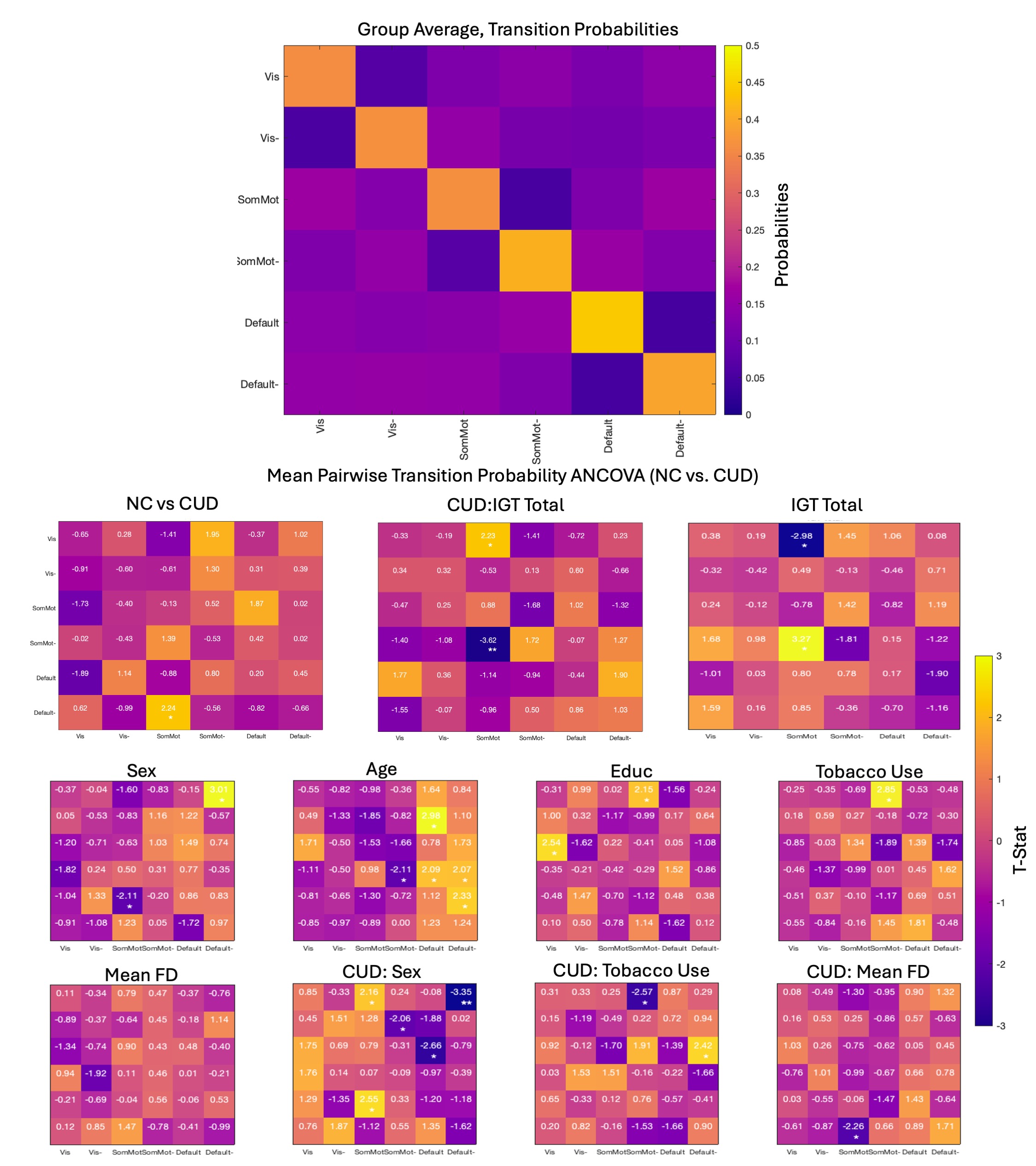

### Supplemental Image 4

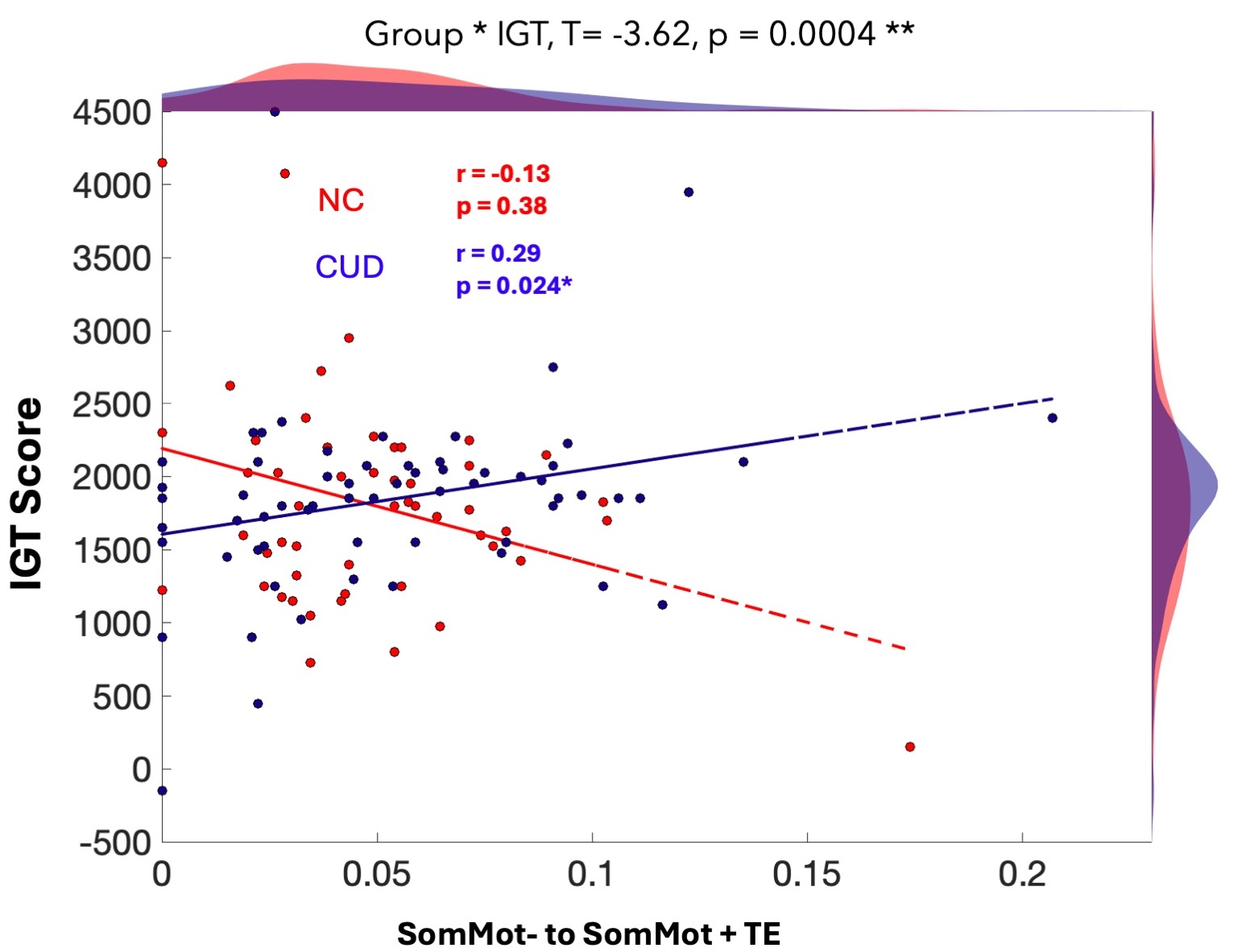

### Supplemental Image 5

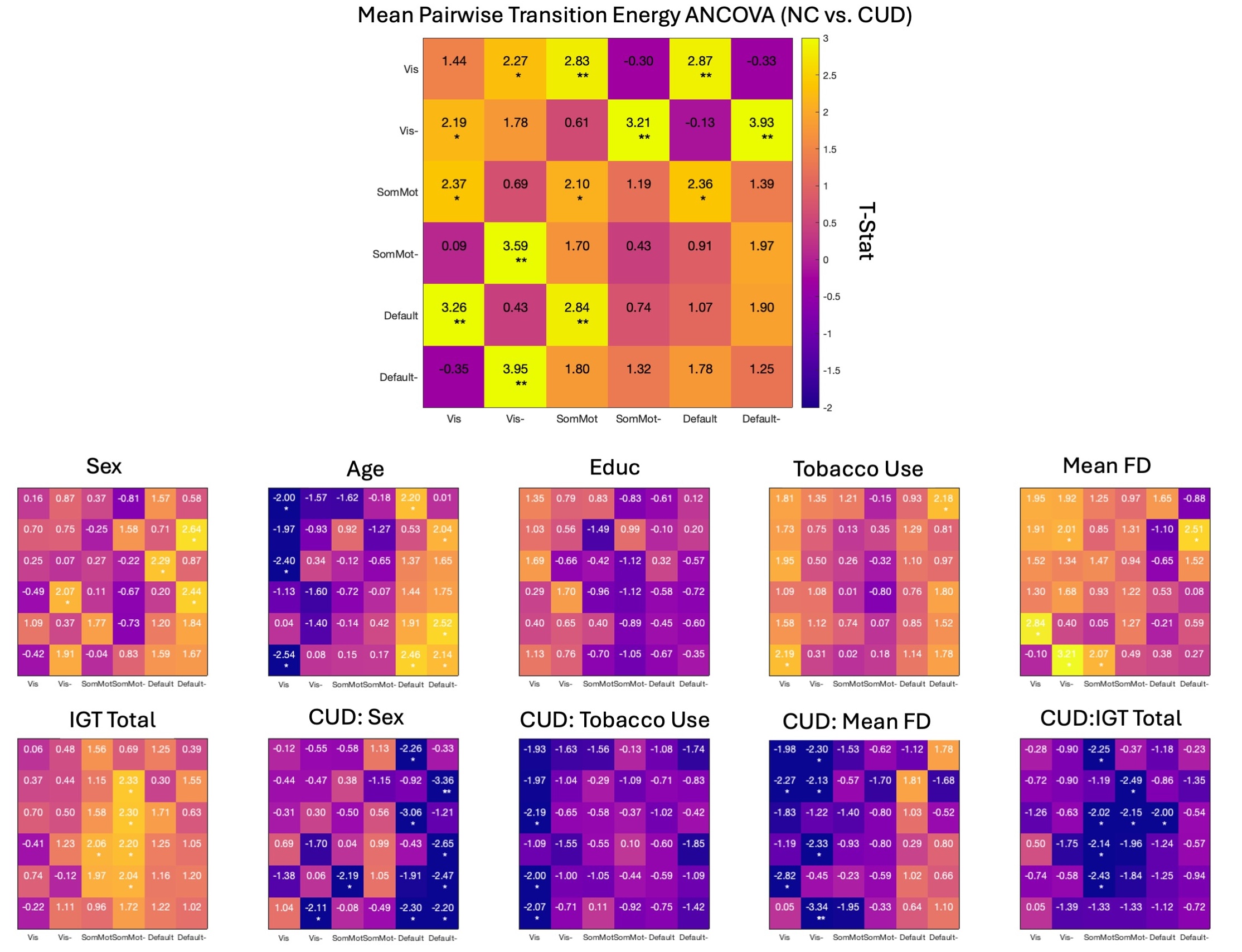

### Supplemental Image 6

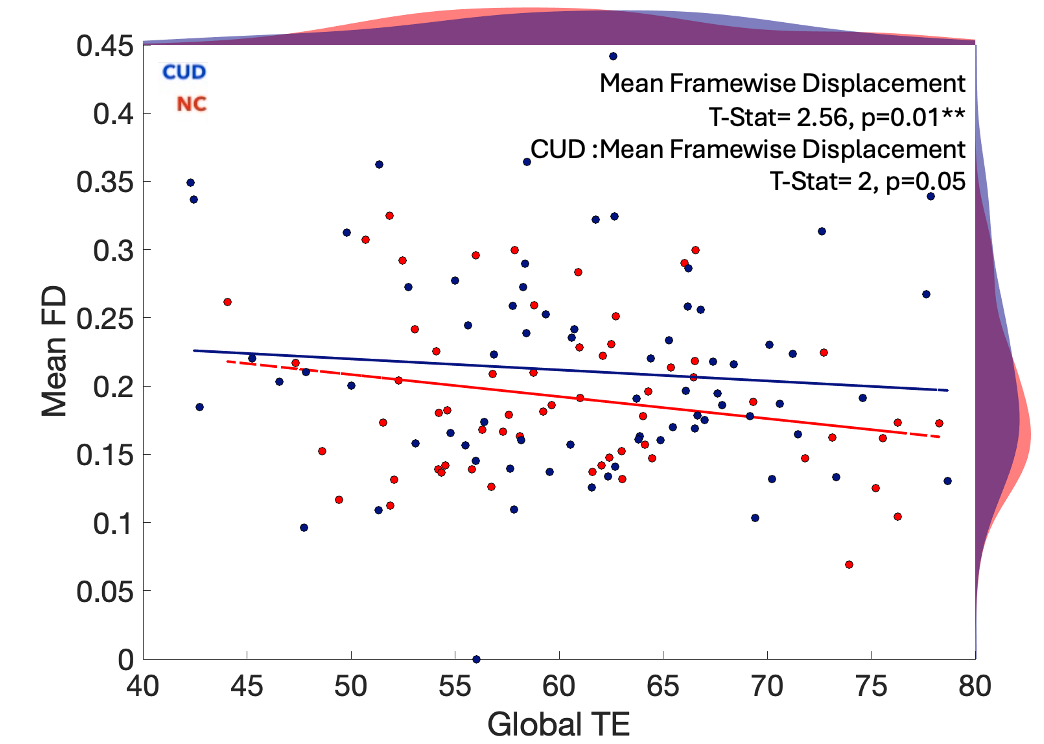

### Supplemental Image 7

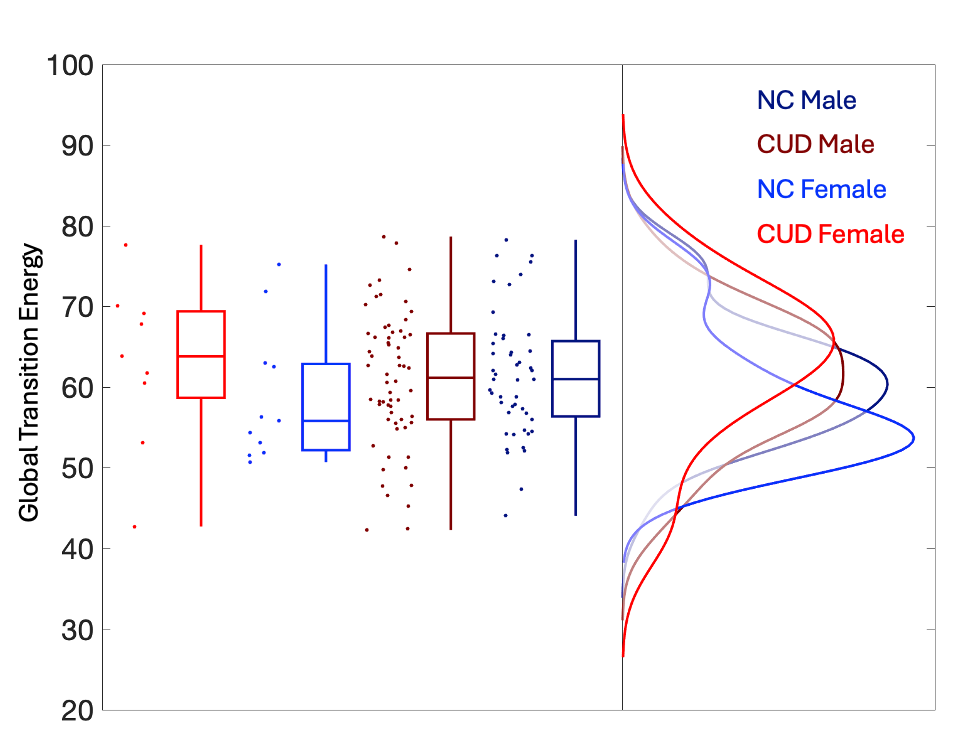

### Supplemental Image 8

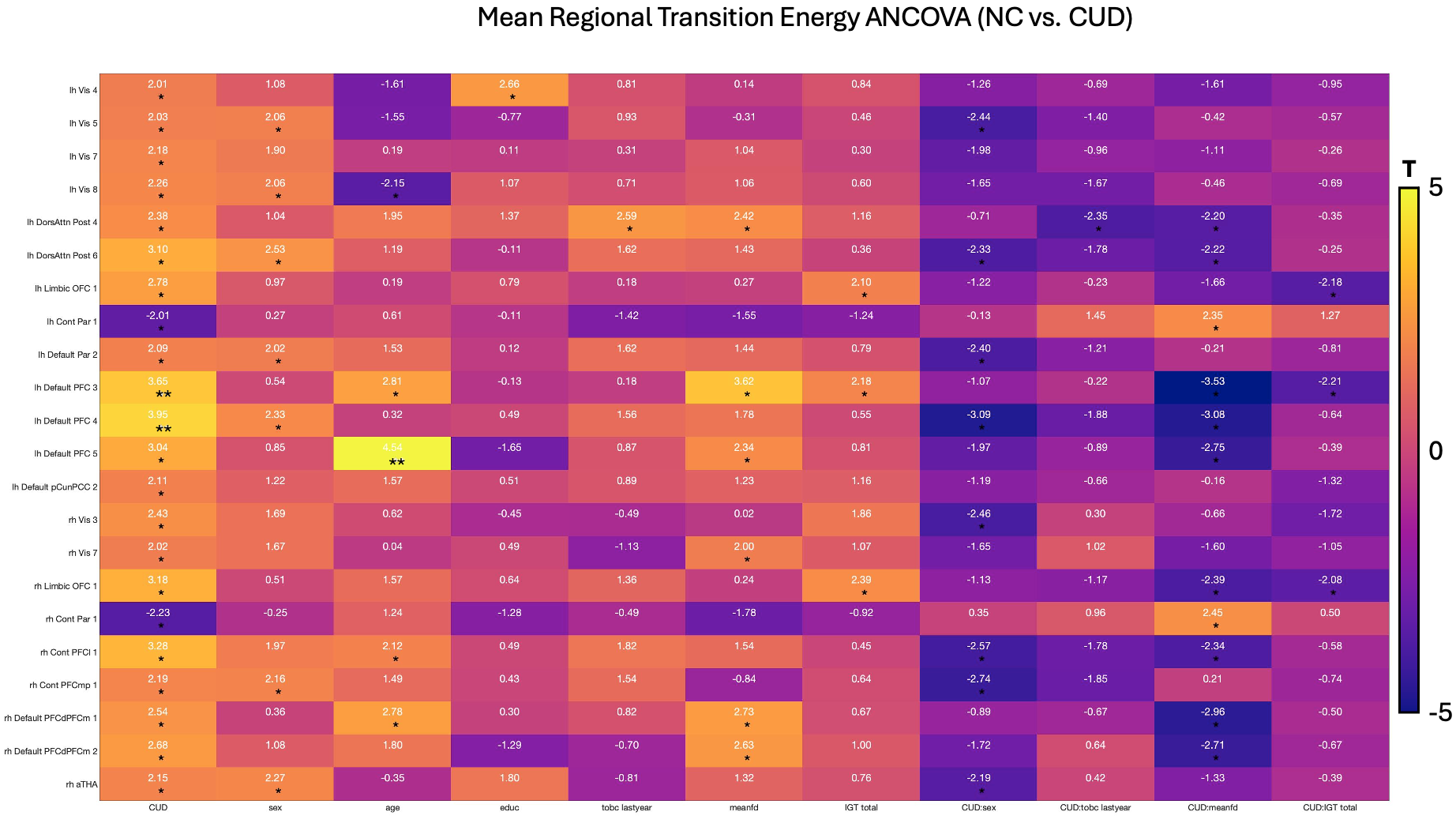

### Supplemental Image 9

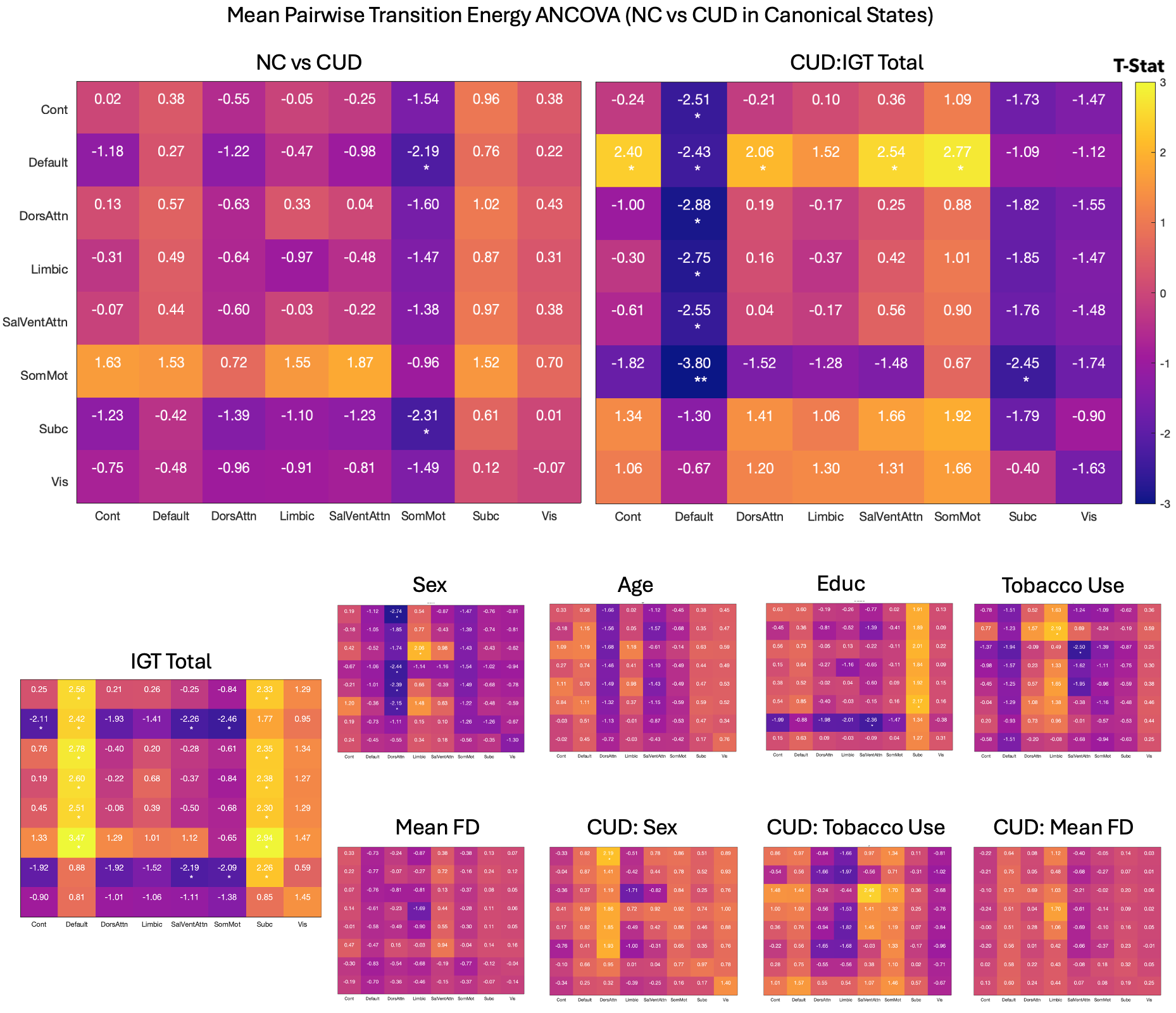

### Supplemental Image 10

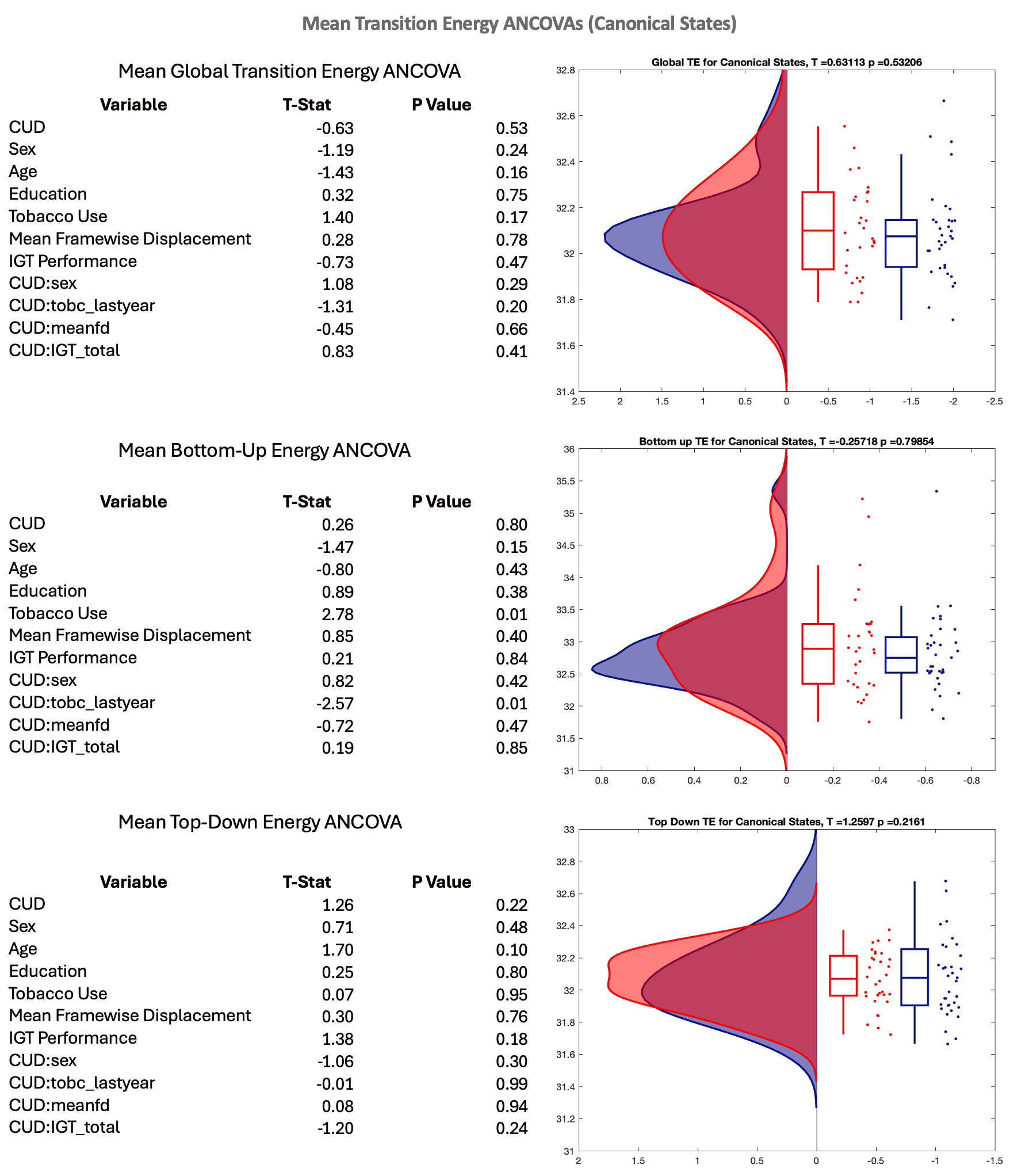

### Supplemental Image 11

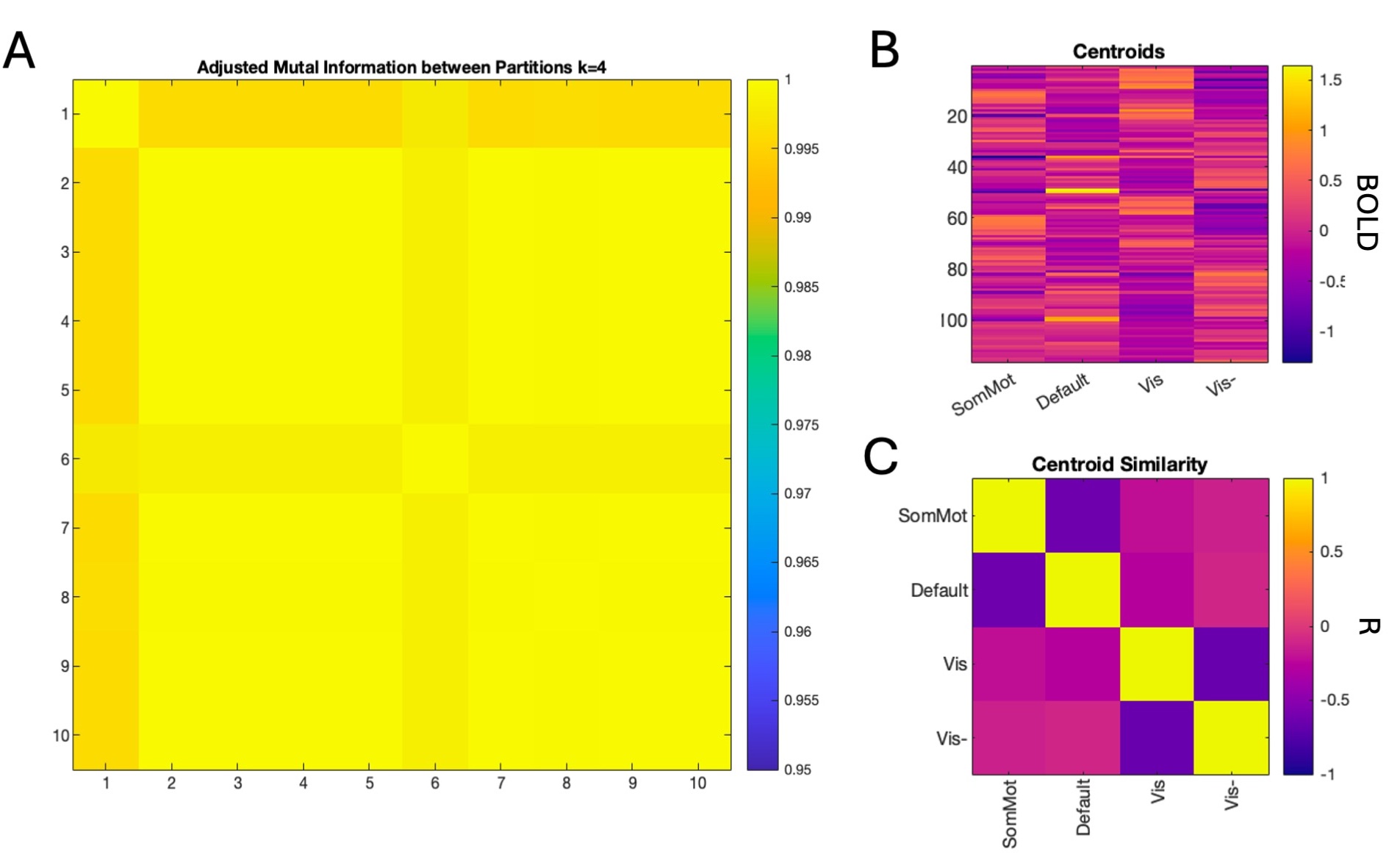

### Supplemental Image 12

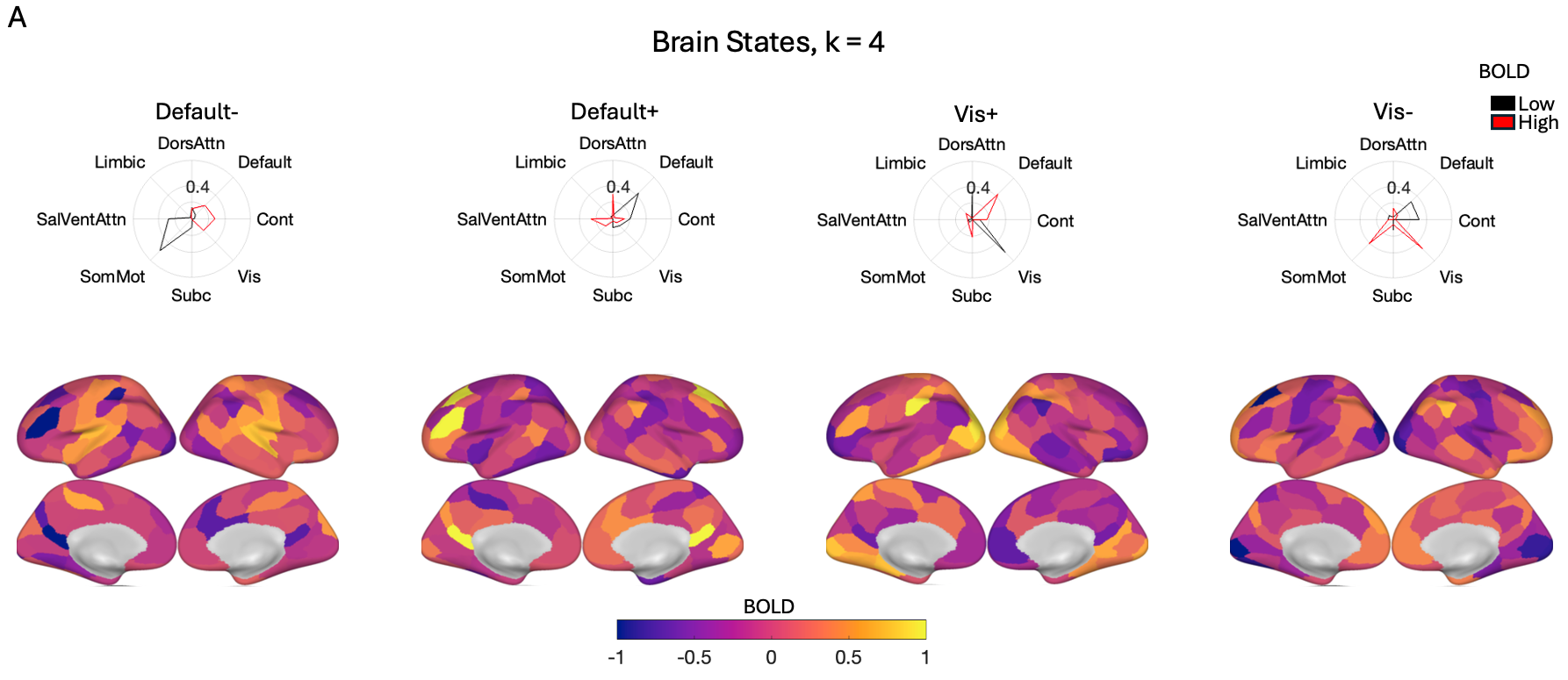

### Supplemental Image 13

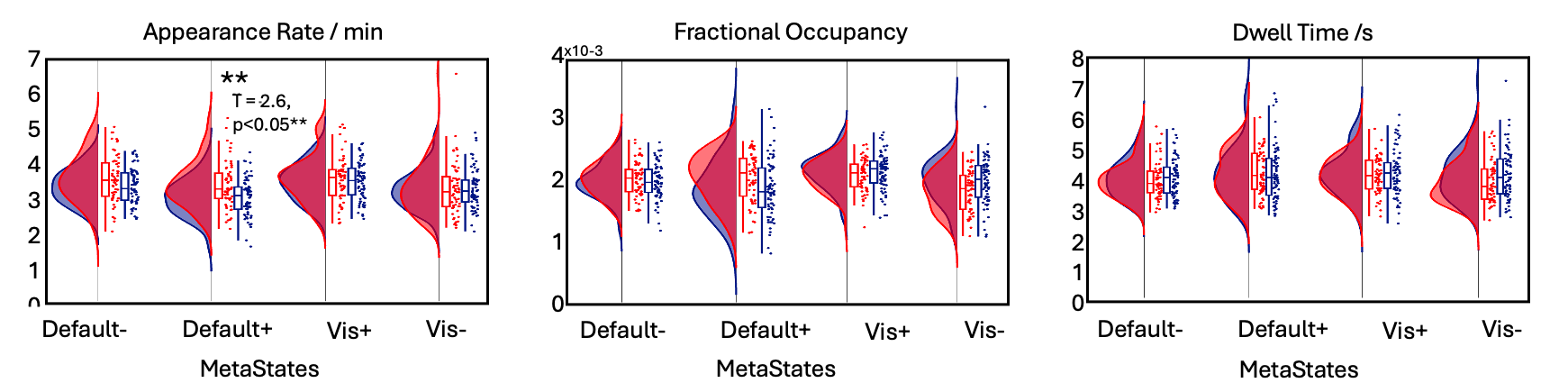

### Supplemental Image 14

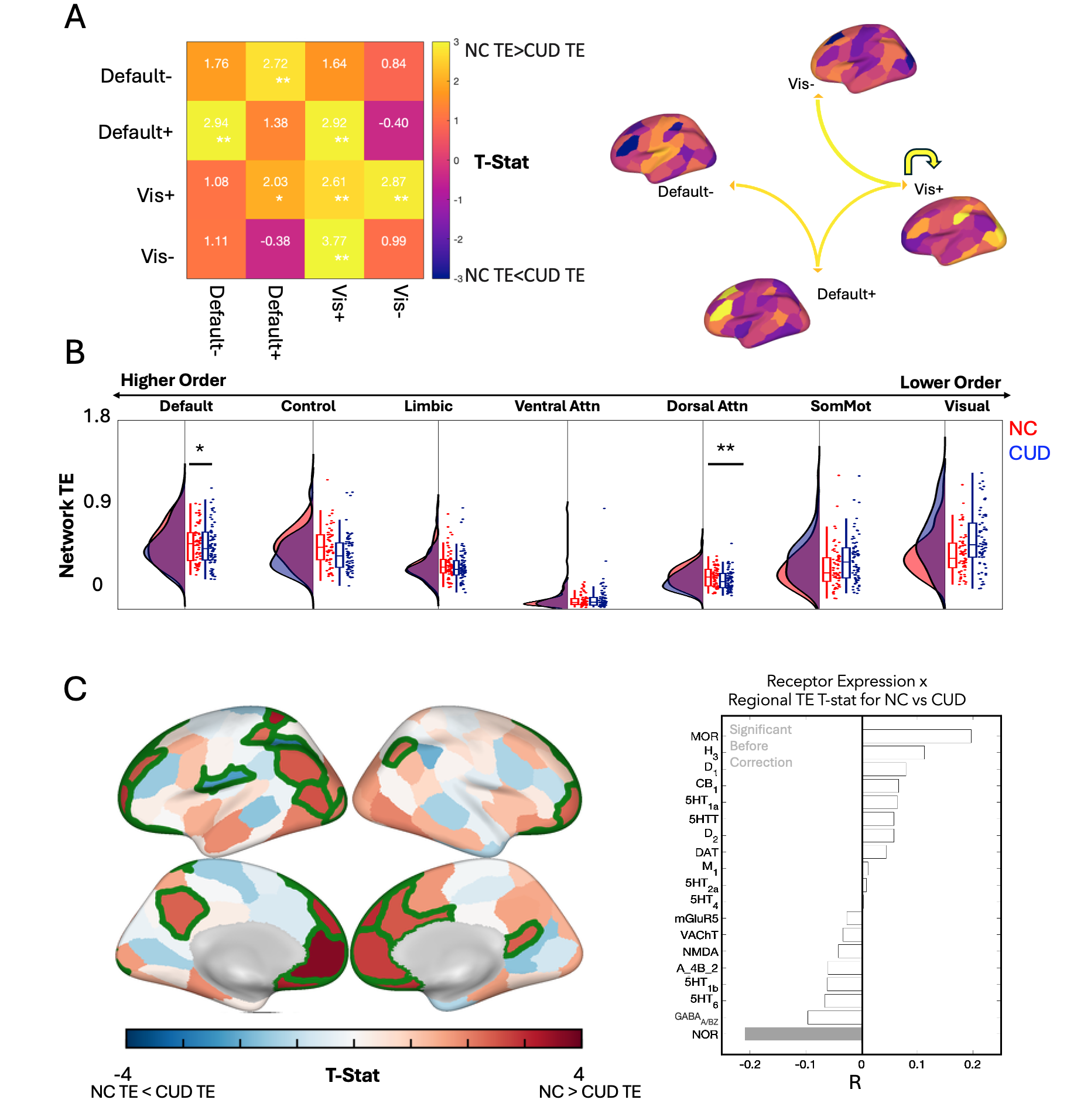

### Supplemental Image 15

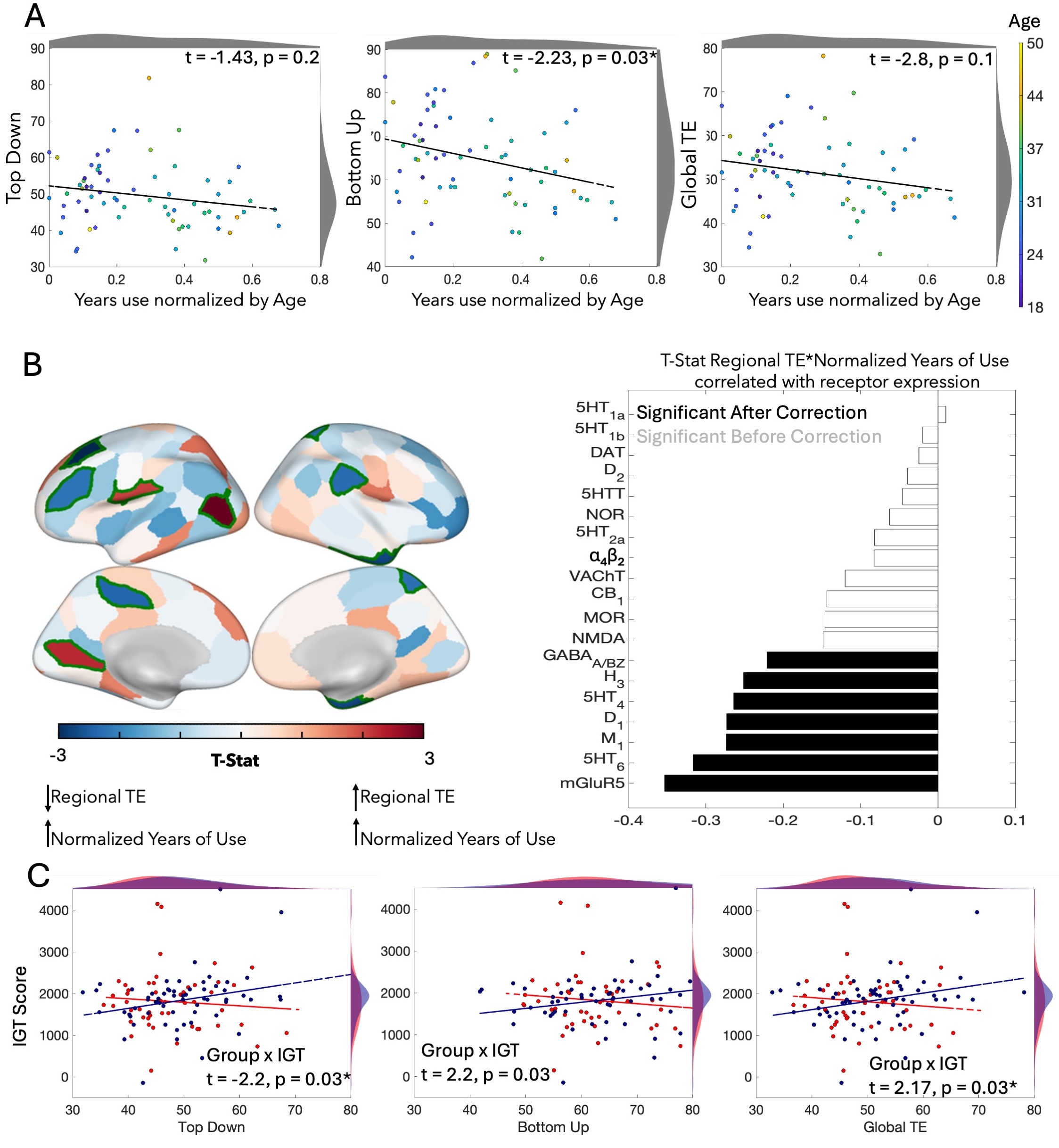

### Supplemental Image 16

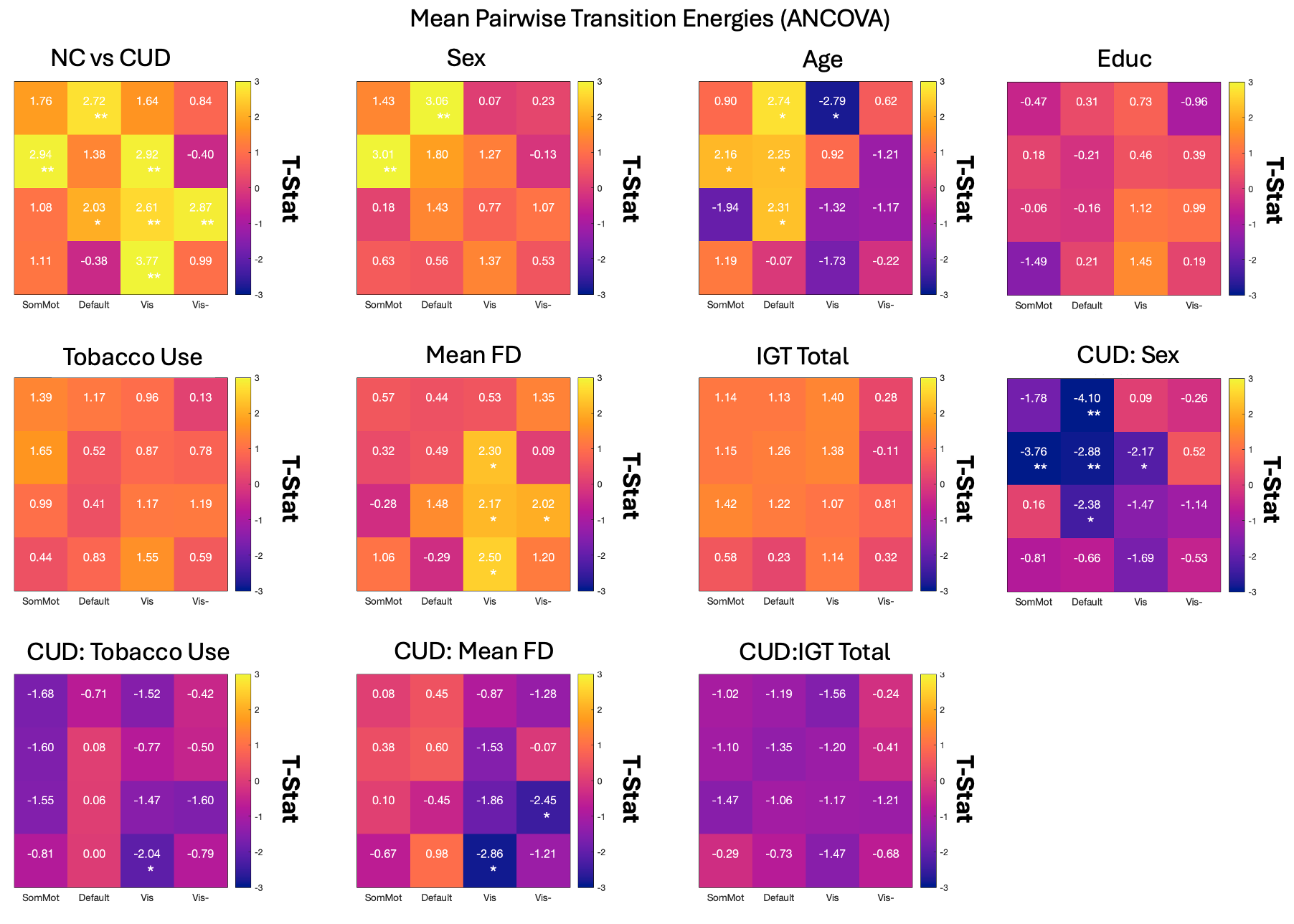

### Supplemental Image 17

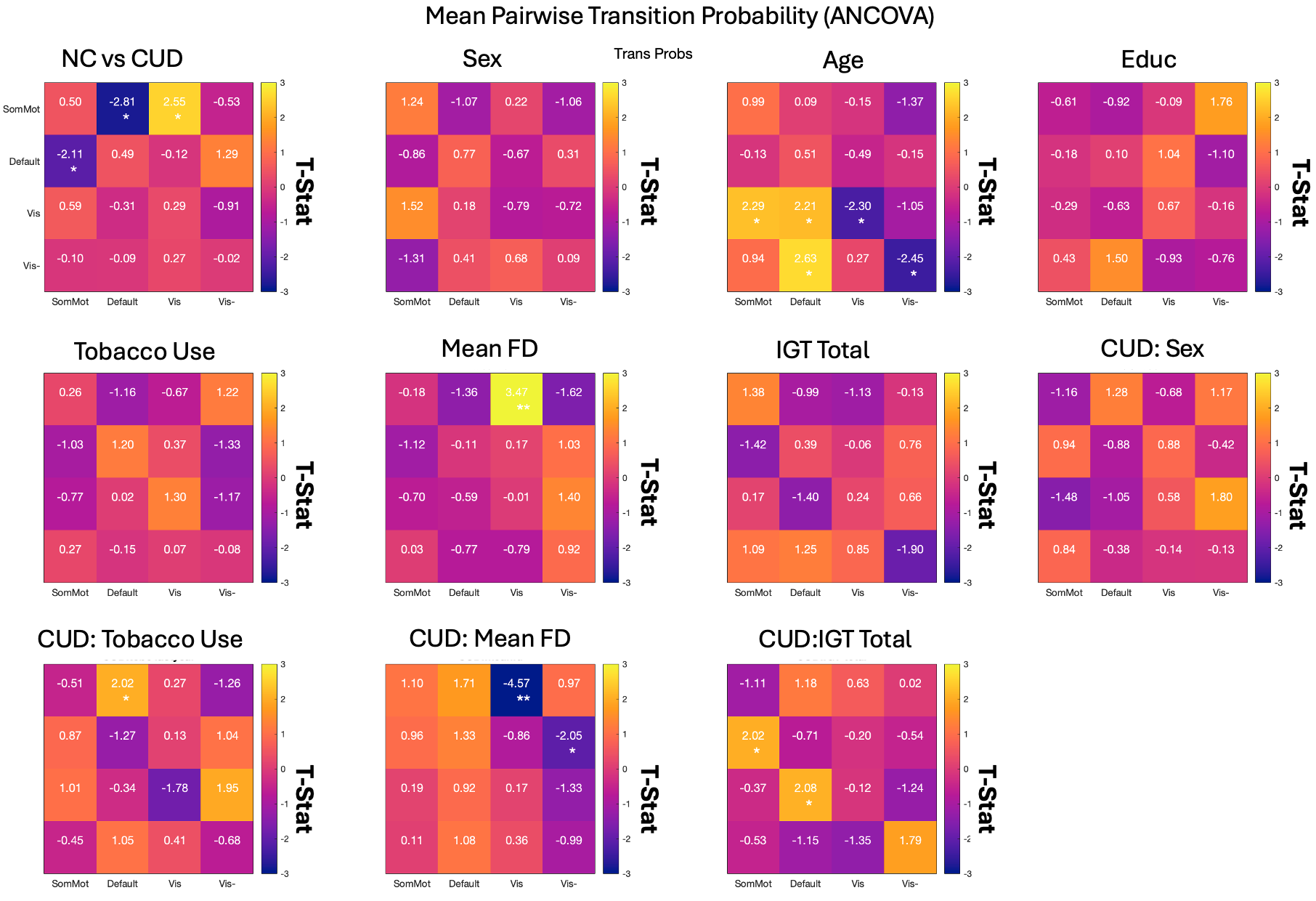

### Supplemental Image 18

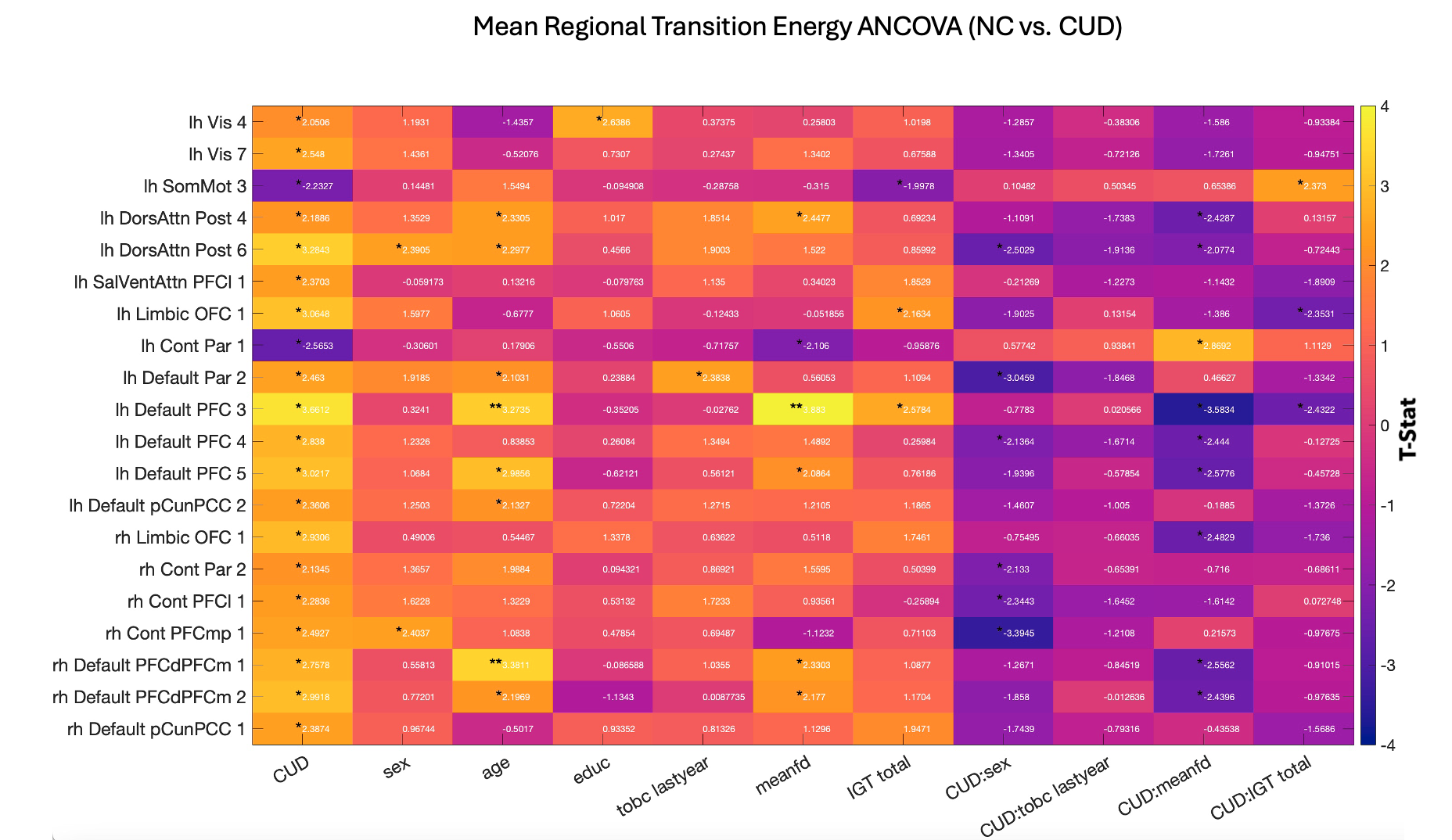

### Supplemental Table 1

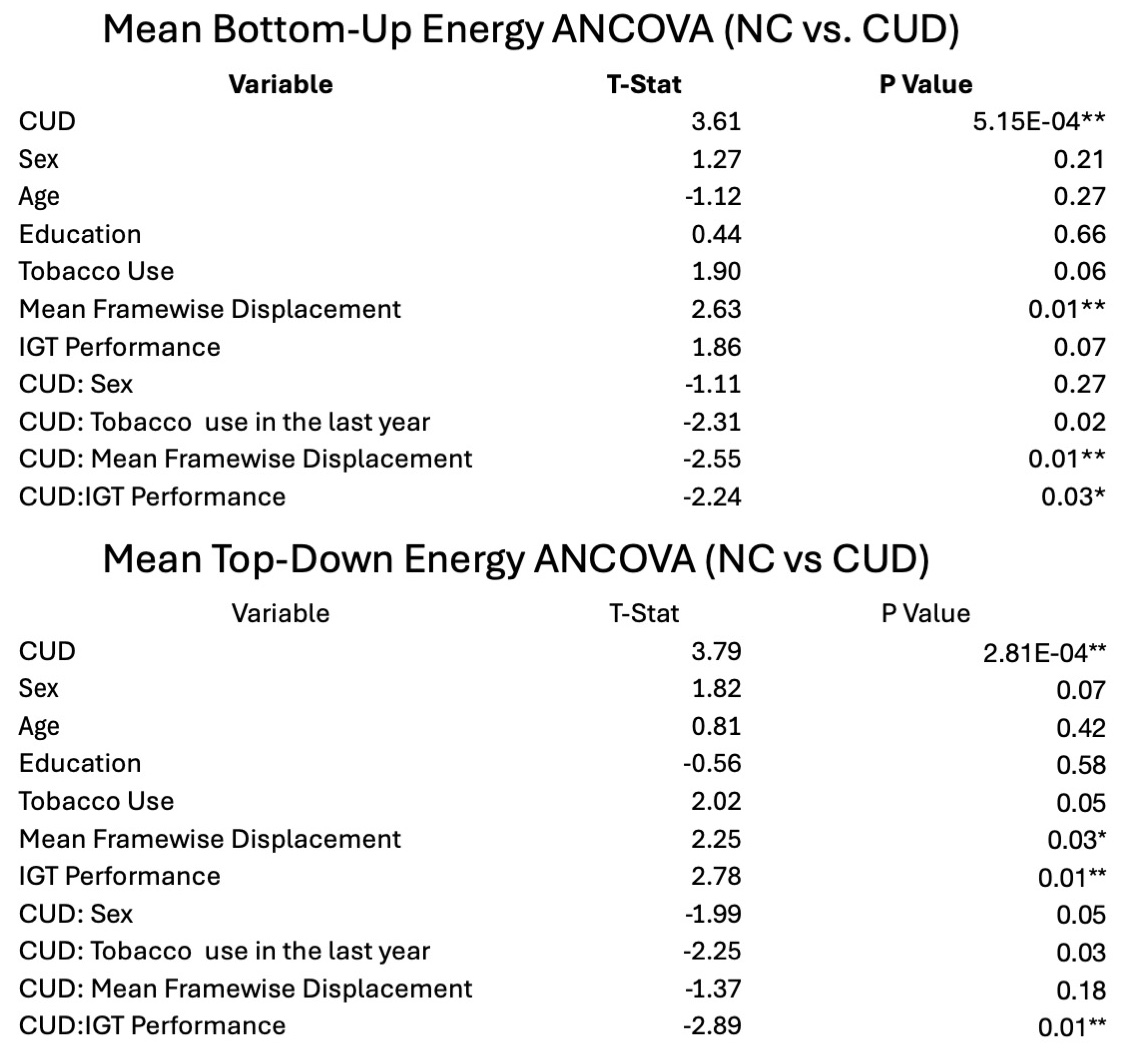

### Supplemental Table 2

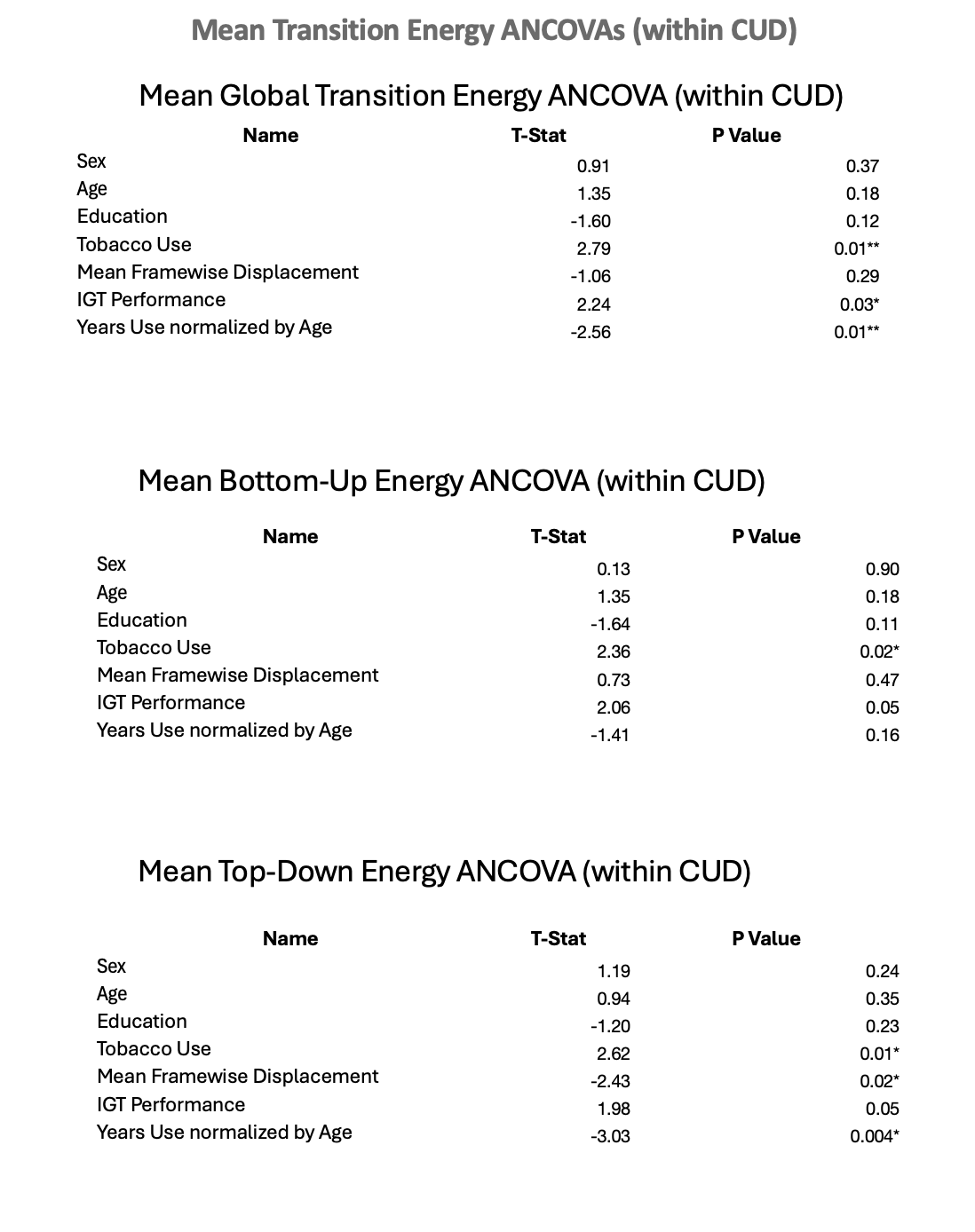

### Supplemental Table 3

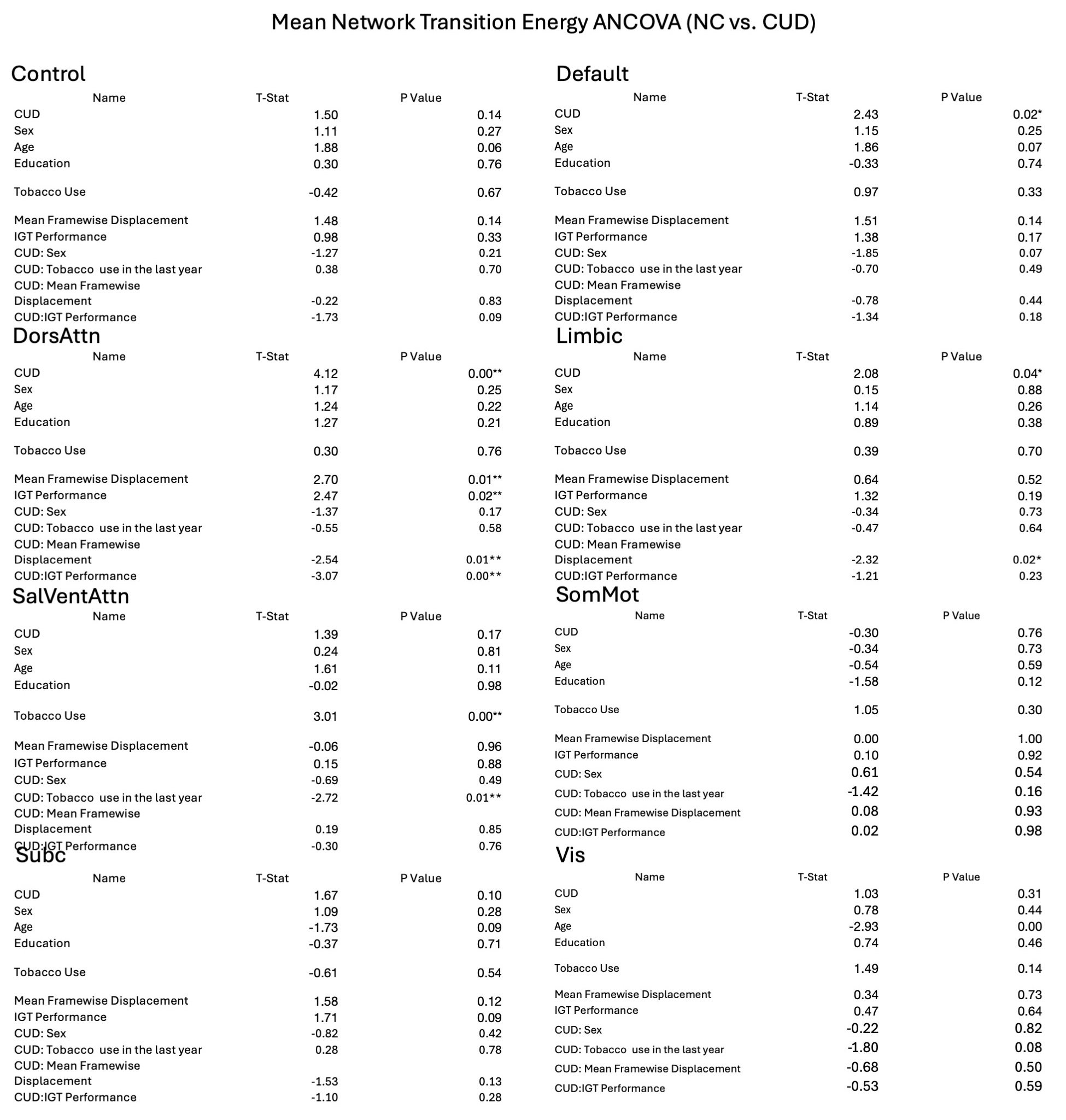

### Supplemental Table 4

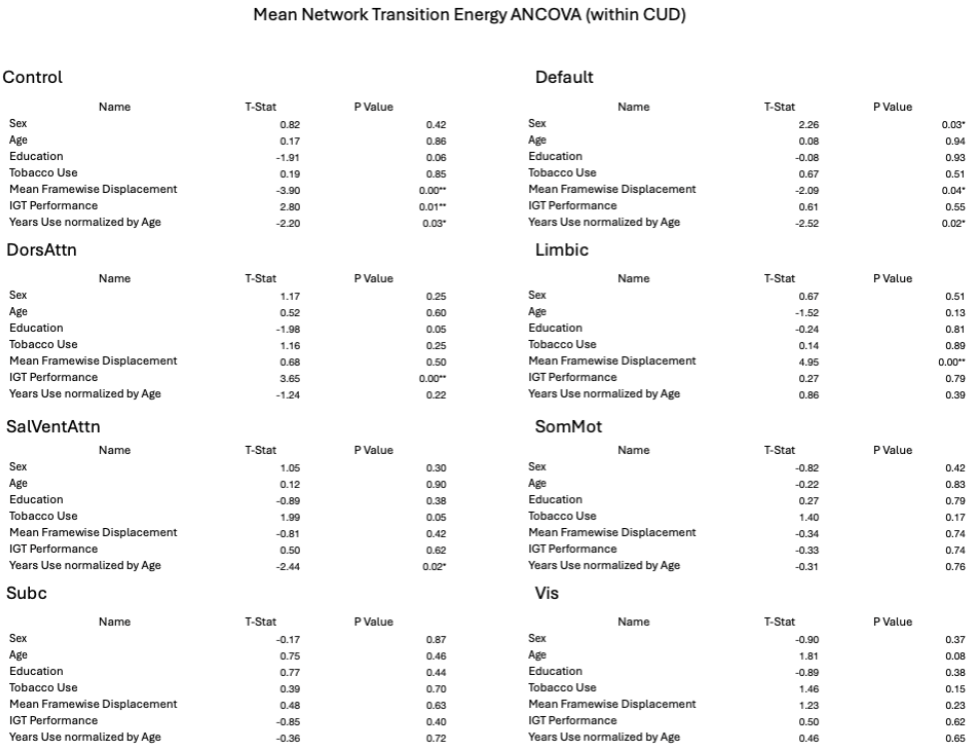

### Supplemental Table 5

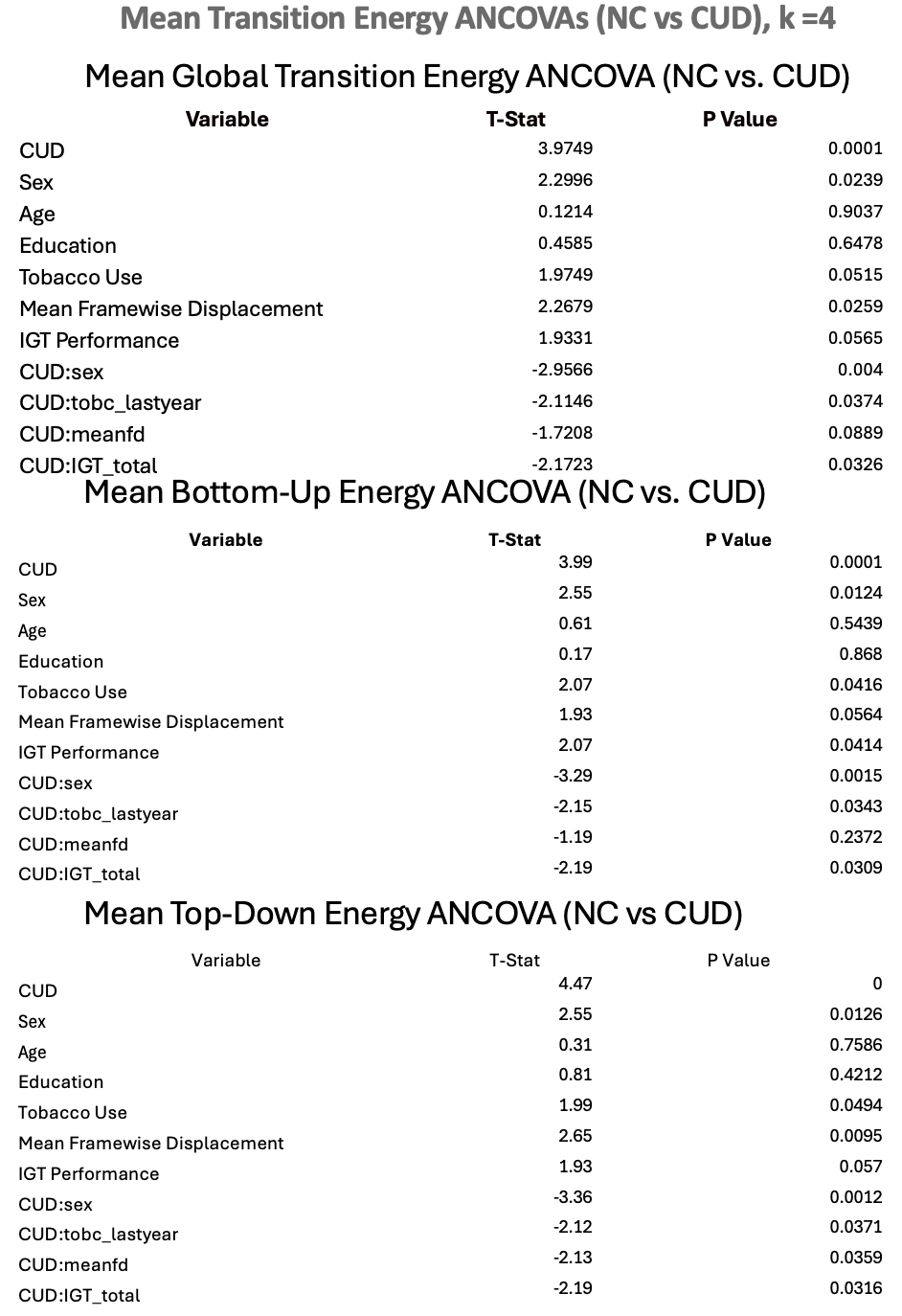

### Supplemental Table 6

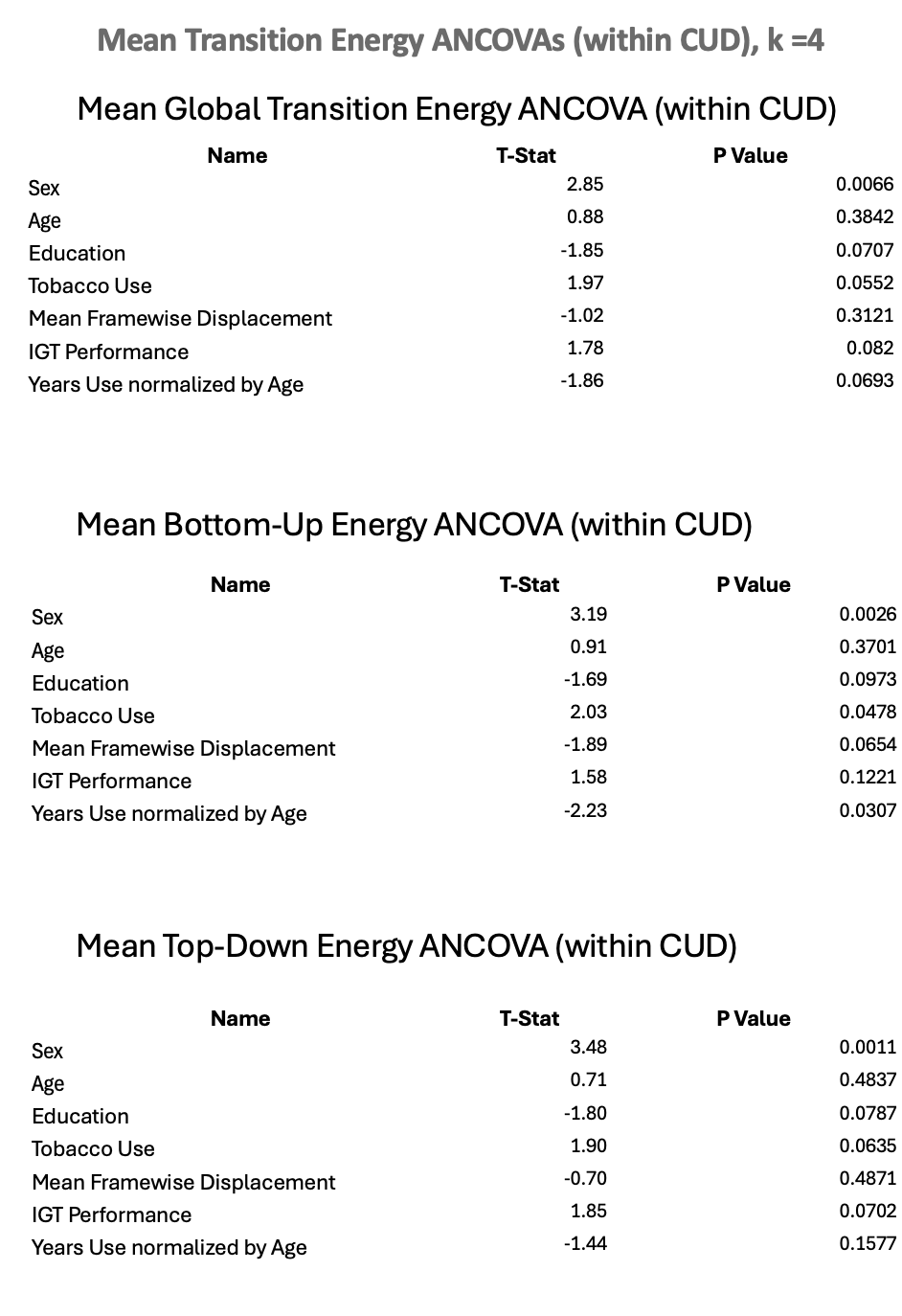

### Supplemental Table 7

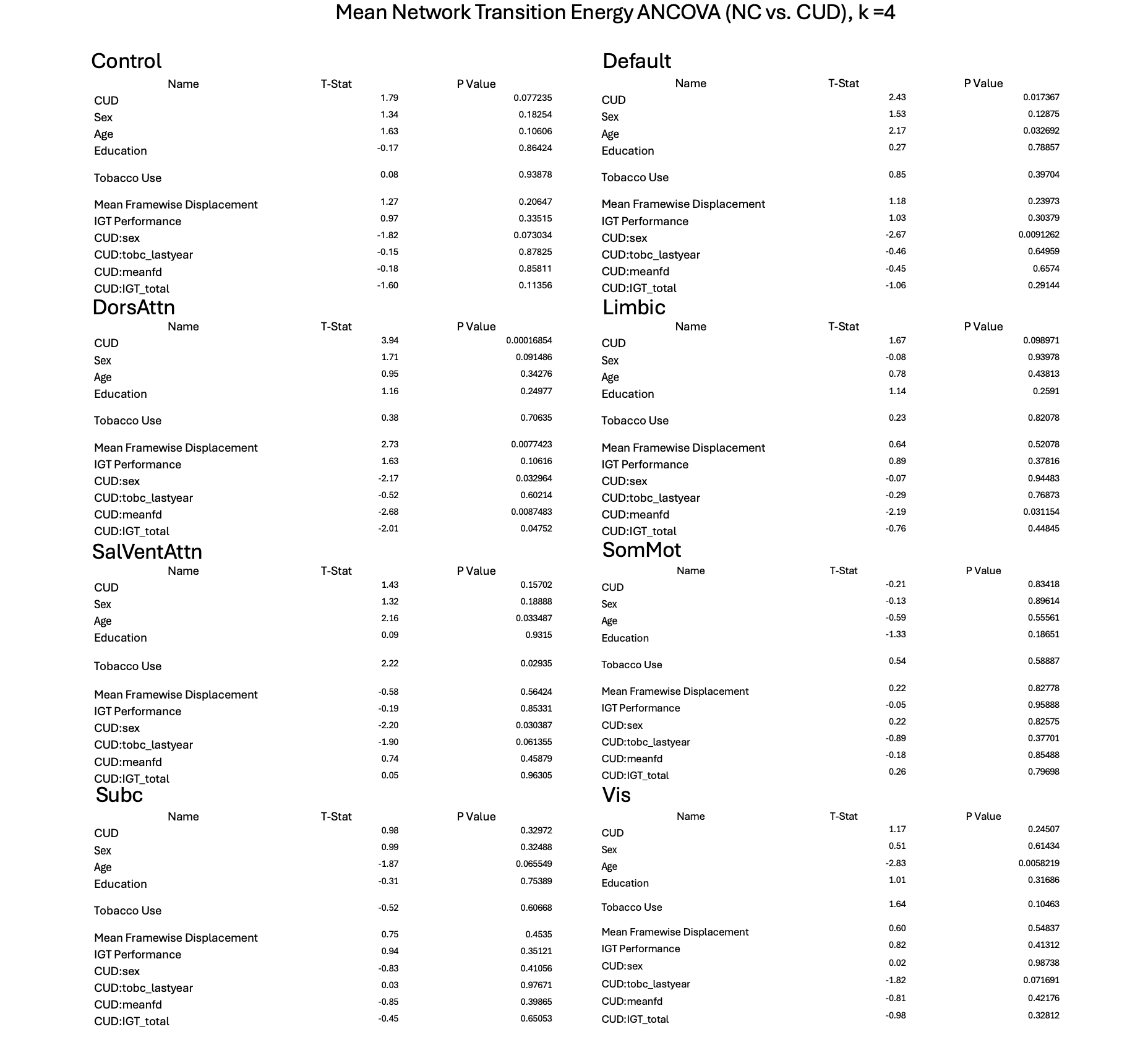

### Supplemental Table 8

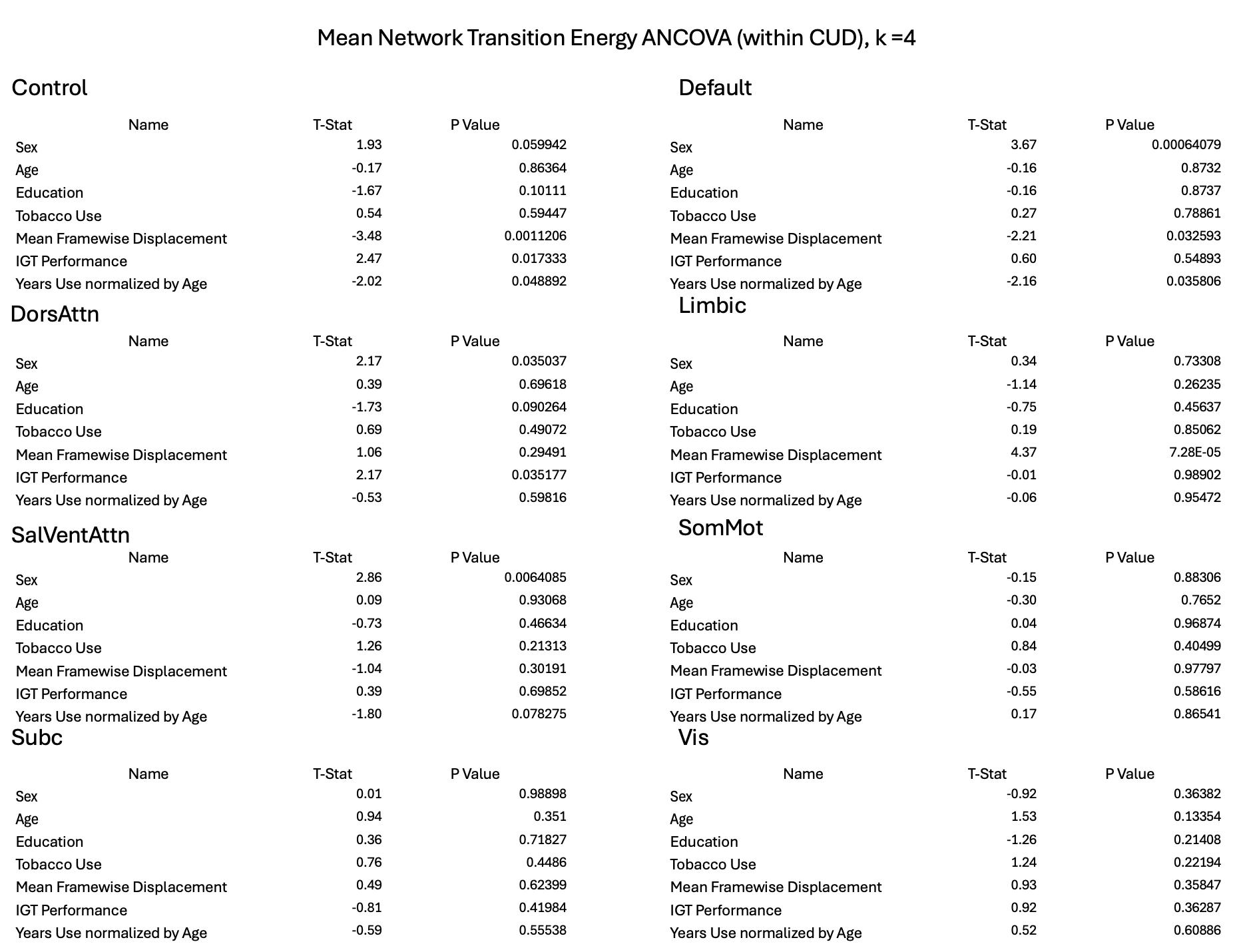
